# Inferring long-term and short-term determinants of genetic diversity in honey bees: Beekeeping impact and conservation strategies

**DOI:** 10.1101/2024.09.04.611184

**Authors:** Thibault Leroy, Pierre Faux, Benjamin Basso, Sonia Eynard, David Wragg, Alain Vignal

**Affiliations:** GenPhySE, Université de Toulouse, INRAE, ENVT, 31326, Castanet Tolosan, France; Abeilles et Environnement, INRAE, Avignon, France; Beebytes Analytics CIC, Roslin Innovation Centre, Easter Bush Campus, Midlothian, UK

**Keywords:** genetic diversity, genomic landscapes, demographic modeling, recombination, *Apis*, honey bees

## Abstract

Bees are vital pollinators in natural and agricultural landscapes around the globe, playing a key role in maintaining flowering plant biodiversity and ensuring food security. Among the honey bee species, the Western honey bee (*Apis mellifera*) is particularly significant, not only for its extensive crop pollination services but also for producing economically valuable products such as honey. Here, we analyzed whole-genome sequence data from four *Apis* species to explore how honey bee evolution has shaped current diversity patterns. Using Approximate Bayesian Computation, we first reconstructed the demographic history of *A. mellifera* in Europe, finding support for postglacial secondary contacts, therefore predating human-mediated transfers linked to modern beekeeping. However, our analysis of recent demographic changes then reveals significant bottlenecks due to beekeeping practices, which have notably affected genetic diversity. Black honey bee populations from conservatories, particularly those on islands, exhibit considerable genetic loss, raising concerns about the long-term effectiveness of current conservation strategies. Additionally, we observed a high degree of conservation in the genomic landscapes of nucleotide diversity across the four species, despite a divergence gradient spanning over 15 million years, consistent with a long-term conservation of the recombination landscapes. Taken together, our results provide the most comprehensive assessment of diversity patterns in honey bees to date and offer insights into the optimal management of resources to ensure the long-term persistence of honey bees and their invaluable pollination services.

## Introduction

Genetic diversity is a fundamental unit of biodiversity, enabling species to withstand threats such as diseases, climate changes, and other evolutionary challenges, ultimately reducing extinction risks. In December 2022, the Kunming-Montreal global biodiversity framework (post-2020) of the UN Convention on Biological Diversity was enacted during the UN Biodiversity Conference (COP15), including important objectives regarding the maintenance of global genetic diversity in order to safeguard the adaptive potential of all species (Goal A & Target 4; CBD, 2022). Recent large-scale multinational evaluations of the evolution of the genetic diversity of wild species through direct estimates (*e.g.* Leigh et al. 2019: 91 species) or indirect indicators (*e.g.* Mastretta-Yanes et al. 2024: 919 eukaryotic species or subspecies) have contributed to report rapid erosion of genetic diversity, thus questioning the long-term sustainability of the current trajectory (see also Esposito-Alonso et al. 2022). Albeit often overlooked in such studies, investigating the evolution of genetic diversity in domesticated species is also crucial, firstly, because it is by definition a part of the global biodiversity, but more importantly; because of the potential consequences of this loss of genetic diversity regarding food security. In that respect, the Kunming-Montreal Global Biodiversity Framework emphasizes the need for equal efforts in preserving the genetic diversity of both wild and domesticated species. Species-specific investigations have however reported extremely rapid genetic diversity loss, including on domesticated (wheat: Pont et al. 2019) and semi-domesticated species (*e.g.* roses, Leroy et al. 2023). Given its central importance in the human diet, the example of wheat is noticeable with an estimated genetic diversity loss of 30% during the 20th century (Pont et al. 2019). In this species, the decline is therefore already far greater than the milestone initially proposed of preserving at least 90% of within-species genetic diversity (CBD, 2021).

Investigating the evolution of genetic diversity in pollinators is especially crucial, given their central role for plant biodiversity, since that around 90% of the >350,000 flowering plant species are pollinated by animals, around 82% of which exclusively by insects including bees (Ollerton et al. 2011; Tong et al. 2023). Currently reported pollinator declines represent a major source of vulnerability for flowering plant reproduction and therefore persistence in ecosystems (*e.g.* Rodger et al. 2021). Bees are not only essential for safeguarding wild plant biodiversity, but also have a profound impact on agriculture. In the late 2000s, it has been estimated that around 10% of the value of the world agricultural production used for human food is dependent on insect pollinators, representing an estimated annual value of more than 150 billion US$ annually (Gallai et al. 2009). For EU countries alone, the economical value of this pollination service was estimated to be around 15 billion US$ (Gallai et al. 2009; Leonhardt et al. 2013), around one third of which could be attributed to a single species, the Western honey bee *Apis mellifera* (Breeze et al. 2011). In addition to this, honey bees - especially Western honey bees - produce economically important products, including honey, beeswax, propolis and royal jelly. Hive products represent significant economic sectors, with increasing demands. For instance, the global honey market alone is estimated to be nearly 10 billion US$ annually and is continuously growing driven by consumers’ expectations for healthy food and natural alternatives to chemical sweeteners. To meet such production needs, modern beekeeping has relied on technical advancements, such as the development of moveable-comb hives in the latter half of the 19th century. Jointly with the production of queens and the control of their fertilization, either by geographical isolation in mating stations or even by artificial insemination (Plate et al. 2019), these advancements have contributed to major changes in the dynamics of the domestication process in honey bees (Crane, 1999; Oxley & Oldroyd, 2010; Weber, 2013). Although Western honey bees are still considered a semi-domesticated species (Oldroyd, 2012), several breeding programs have been implemented, sometimes including full control of queen fertilization through artificial insemination, questioning the recent impact of the intensification of beekeeping practices on the evolution of genetic diversity of honey bees.

Honey bees belong to the genus *Apis*, which comprises around ten eusocial species with a haplodiploid sex-determination system (Ruttner, 1988; Arias & Sheppard, 2005). All *Apis* species are native to Southeast Asia, with the notable exception of *A. mellifera* whose original distribution area covers Europe, Africa, and the Middle East. Within this native range, about thirty subspecies of *A. mellifera* have been identified both morphologically and genetically, grouped into five main lineages: African (A), Arabian (Y), West European (M), Central European (C), and Caucasian (O) (Ruttner; 1988; Cornuet & Garnery, 1991; Hall & Smith, 1991; Whitfield et al. 2006; Alburaki et al. 2013; Wallberg et al. 2014; Ilyasov et al. 2020). The evolutionary history of Western honey bees is a subject of ongoing study. Major uncertainties remain regarding the exact origin of the species in Asia or Africa (The Honeybee Genome Sequencing Consortium, 2006; Whitfield et al. 2006; Han et al. 2012; Cridland et al. 2017; Dogantzis et al. 2021) and the pathways of honey bee colonization into Western Europe (Near East or North Africa, Han et al. 2012). However, whatever the exact origin of *A. mellifera,* its current worldwide distribution is entirely mediated by man, due to interests in honey and beeswax production. For instance, introductions were reported in North-America and Oceania during the 17th and 19th century, respectively (Hopkins, 1911; Carpenter & Harpur, 2021). Comprehensive studies have highlighted the relative contributions of the M-, C-, and O-lineages in shaping the current genetic diversity used by beekeepers (Wragg et al. 2016; 2022; Carpenter & Harpur, 2021). In the recent decades, C-lineage bees, including the *A. m. carnica* and *A. m. ligustica* subspecies, have become predominantly used by professional beekeepers, due to their gentleness and high honey production (e.g. Ruttner, 1988; Hoppe et al. 2020), resulting in the dominance of this lineage over the other genetic backgrounds (*e.g.* Harpur et al. 2015; Carpenter & Harpur, 2021). Intentional crossbreeding between subspecies has given rise to new breeds for honey production, such as the ‘Buckfast’ (Adam, 1996), which rapidly became one of the most popular breeds among beekeepers. Lines have also been specifically selected for increased production of royal jelly (Wragg et al. 2016).

Recently, population genomics approaches by whole genome sequencing hundreds of Western honey bee samples, have contributed to substantially increase the amount of publicly available genomic resources for this species, with several thousands of genomes now available (Harpur et al. 2014; Wallberg et al. 2014; Chen et al. 2018; Wragg et al. 2022; Cao et al. 2023; Eynard et al. 2024, among others). Recently, our research group at INRAE Toulouse, France, inferred the population structure of honey bees present in Western Europe, with a particular focus on France, based on the whole-genome sequencing of over 800 *A. mellifera* drones (Wragg et al. 2022). By including specific human-managed populations, including the study of hybrids, those used for honey and royal jelly production, as well as black *A. m. mellifera* honey bees kept in conservatories, this work contributed to a detailed description of the genetic makeup of different natural subspecies and of the influence of beekeeping. Among the three main lineages (M, C and O), nine genetic backgrounds were identified (Fig. 1A). Four clusters belongs to the M-lineage, including a cluster composed of *A. m. iberiensis* samples (referred to as ‘Iberiensis’) and three *A. m. mellifera* clusters from black honey bee conservatories (one French Mediterranean population referred to as ‘Mellifera’, and the two isolated Atlantic islands in Brittany and Scotland, referred to as ‘Ouessant’ and ‘Colonsay’). Similarly, two clusters belonging to the C-lineage were identified, composed of *A. m. ligustica* and *A. m. carnica* samples (referred to as ‘Ligustica’ and ‘Carnica’, respectively). A single population of the O-lineage was also identified composed of *A. m. caucasia* (‘Caucasia’). In addition, an eighth background was identified in samples from the honey bees selected by the French royal jelly breeders’ organization (GPGR) for the production of royal jelly (‘RoyalJelly’) and a ninth appears in populations that were assumed to be a type of Buckfast bees (‘Buckfast’). Other honey bee species have been studied by whole genome sequencing and available datasets including a few hundreds of individuals of the Eastern honey bee *A. cerana* (Chen et al. 2018; Wang et al. 2024) and more than fifty giant honey bee samples (*A. laboriosa* and *A. dorsata*, Cao et al. 2023). These datasets have contributed to better describe the genetic structure within these three other *Apis* species.

**Figure 1:**
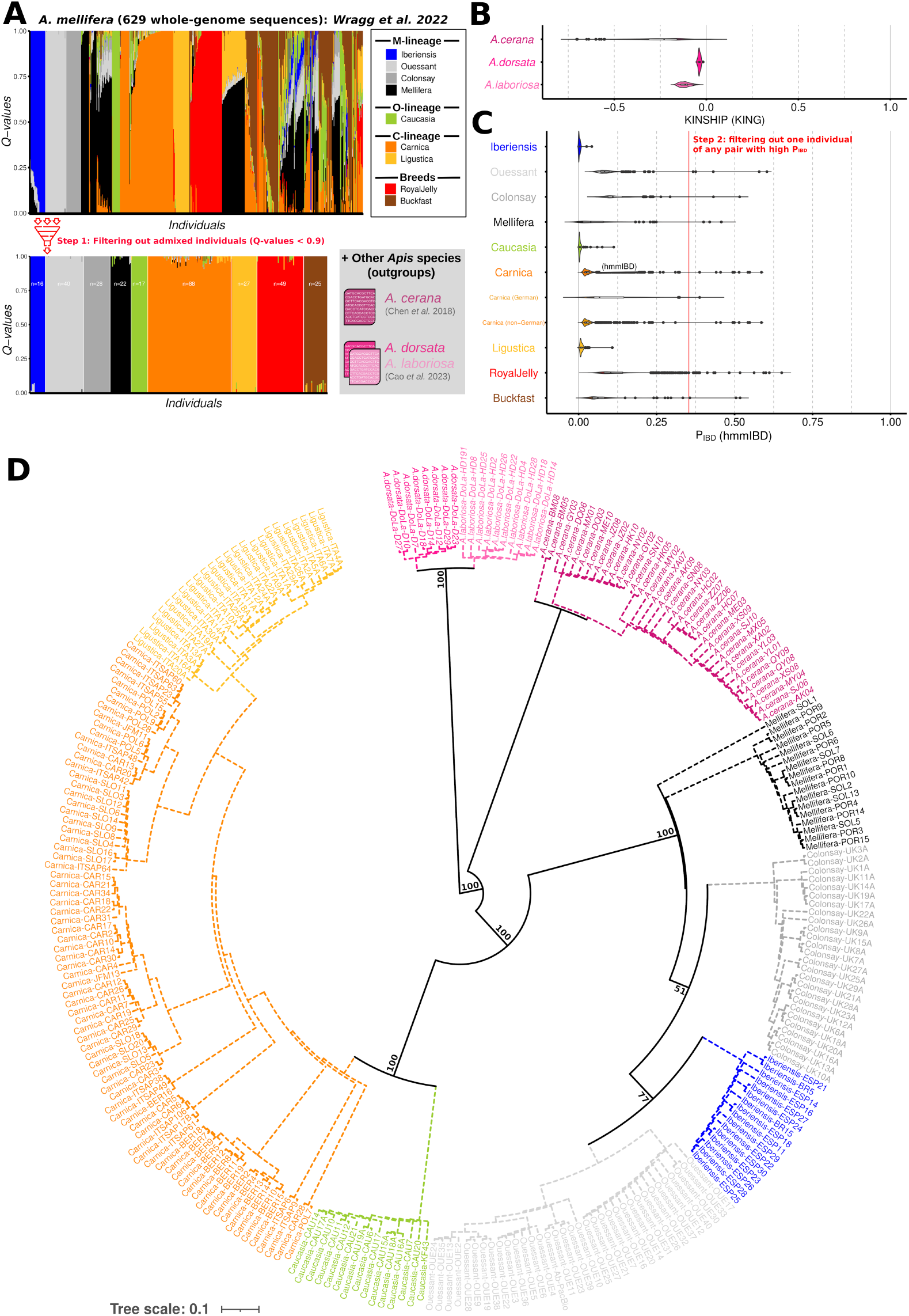
Population structure and kinship of the honey bees investigated in this study. **A.** Population structure of the Apis mellifera samples used in the study. Filtering for individuals with elevated memberships to one of the 9 clusters identified by Wragg et al. 2022 among the 629 individuals, allowing to identify 312 well-assigned individuals. Values correspond to Table S7 from Wragg et al. 2022. **B.** Kinship estimates based on all workers (diploid) of the three outgroup species, as inferred with KING. **C.** Kinship values for all haploid drones as computed with hmmIBD. Individuals with elevated P_IBD_ (above the red threshold) were subsequently discarded from the dataset (see Table S2). **D.** Consensus tree from 200 replicates generated with the Kmer approach implemented in PanTools. Bootstrap values are only shown for the main nodes (solid lines; for details, see Fig. S2).

Despite its importance, the levels of genetic diversity has been less studied in honey bees, although some preliminary results were published on global values (Mikheyev et al. 2015; Espregueira Themudo et al. 2020), or on determinants of its variation along the genome (Wallberg et al. 2015). Moreover, the impacts of modern beekeeping on the levels of genetic diversity is an example of a topic of substantial ongoing debate among bee geneticists (Harpur et al. 2012; 2013; De la Rua, 2013).

In this study, we leveraged extensive genomic resources to investigate both short-term and long-term trajectories of nucleotide diversity. Specifically, we aimed to: (i) elucidate the evolutionary history of honey bees, including the presence and timing of gene flow between lineages; (ii) assess global nucleotide diversity levels in the nine *A. mellifera* subspecies and three other *Apis* species; (iii) examine recent demographic changes linked to modern beekeeping and their effects on current honey bee diversity; (iv) analyze the correlations of nucleotide diversity variations across the genome (referred to as genomic landscapes) among the different genetic clusters and species, covering a divergence gradient of more than 15 million years; and (v) explore the role of recombination in shaping these genomic landscapes of nucleotide diversity.

## Results

### Population structure and kinship

We gathered publicly available whole-genome sequence data of a total of 683 individuals from four honey bee species of the *Apis* genus, spanning more than 15 million years of divergence (TimeTree, median: 16.8, CI: 15.1 – 17.0 ; see also Engel 2006; Ramírez et al. 2010; Martins et al. 2015). Specifically, we analyzed 629 haploid drones from *A. mellifera*, which were previously sequenced in our lab (Wragg et al. 2022), plus 54 diploid workers from three other species, 8 from *A. dorsata*, 10 from *A. laboriosa* and 36 from *A. cerana* (see Table S1 & File S1).

Among the *A. mellifera* samples, Wragg et al. (2022) previously identified 9 genetic clusters, corresponding to 3 main lineages: the Western European lineage M (4 clusters: ‘Iberiensis’, ‘Mellifera’, ‘Iberiensis’, ‘Colonsay’ and ‘Ouessant’), the Central European lineage C (2 clusters: ‘Ligustica’ and ‘Carnica’) and the Caucasian lineage O (1 cluster: ‘Caucasia’), plus two genetic clusters that were considered as two breeds, respectively selected for the specific production of royal jelly (‘RoyalJelly’) and for honey production (‘Buckfast’). To know more about the history of the different genetic clusters identified by Wragg and collaborators (2022), we first excluded admixed individuals by discarding those exhibiting Q-values lower than 0.9 (Fig. 1A), leaving 312 haploid individuals for further analyses. To further check the absence of remaining substructure within the previously inferred clusters, we randomly sampled around 200,000 biallelic SNPs among the 8,949,504 high-quality SNPs in our dataset and then use PCAs to investigate the within-group population structuration (Fig. S1). Assuming panmixia, one might expect each individual to contribute to an independent PC, with a variance explained of roughly 1/n individuals included in the analysis. Based on this criteria, we excluded all individuals acting as simple outliers with an individual contribution exceeding 50% more than this simple expected value (see Table S2 and Fig. S1). Assuming the same criteria, we excluded 6 individuals from the Buckfast cluster that formed a small cluster on the PC1 (PC1: 10.22% of variance explained; expectation: 4.00%; enrichment: 2.56). For the Carnica, two subgroups were also identified, separating all except two samples from Germany (hereafter referred to as the German samples) from all other samples (PC1: explained variance 2.83%, expected: 1.14%, enrichment: 2.48; all other Carnica samples are then referred to as non-German samples).

To perform unbiased inferences and estimates of nucleotide diversity, we also excluded individuals with close family relationships by performing kinship analyses. To take into account the variable ploidy levels among the samples (haploid drones *vs.* diploid workers), two different methods have been used (hmmIBD and KING for haploid and diploid samples, respectively). For the haploid samples, we identified 89 pairs of individuals with elevated kinship values, corresponding to an average of 0.29 pairs per individual (min=0, max=4), for which we subsequently filtered out one individual per pair. Based on this analysis, we excluded a total of 33 additional samples, hence eventually retaining 267 haploid drones. Among the 54 diploid workers, we found no evidence for close family relationships (Fig. 1B). Importantly, close exploration of the kinship results provides important practical information for beekeepers, conservatory managers, and other honey bee experts that are discussed in detail in Sup. Note 1.

### Phylogenetic reconstruction

For all subsequent analyses, we reconstructed whole-genome sequences from a VCF containing both variant and non-variant positions, focusing only on observed variation at SNPs (for details, see Materials and Methods). To gain insights into the phylogenetic relationships in the dataset, we performed a neighbor-joining (NJ) tree reconstruction using a pangenomic approach based on a K-mer distance matrix (Figs. 1D & S2). Among the different *Apis* species, we observed a first split between the commonly used honey-producing honey bees (*i.e.* the Eastern and Western honey bees, *A. cerana* and *A. mellifera*) from the so-called giant honey bees, namely *A. laboriosa* and *A. dorsata*, the Himalayan and the Southeast Asian giant honey bees, respectively. Subsequently, the tree is consistent with a second split between the Eastern and Western honey bees. Within the Eurasian lineages of *A. mellifera*, we found unambiguous bootstrap support (100%) for a first split between the M-lineage (*A. mellifera mellifera* and *A. mellifera iberiensis*) and the ancestor of the C- and O-lineages, followed by a similarly well supported split between the C-(*A. mellifera carnica* and *A. mellifera ligustica*) and O-lineages (*A. mellifera caucasia*). Overall, the genetic clusters are largely monophyletic, with the notable exception of the Carnica cluster (in orange, Fig. 1D). Surprisingly, we found substantial evidence, albeit with bootstrap support lower than 80% (51% and 77%, Fig. 1D), for the closest proximity between the genetic clusters corresponding to conservatories located on the Atlantic coast (Colonsay, Scotland and Ouessant, Northwestern France) and the Iberiensis cluster (*A. mellifera iberiensis*), compared to samples from the Mediterranean conservatory (*M. mellifera*, Southwestern France). Given that Colonsay, Ouessant, and Mellifera are expected to represent different samples of a single subspecies (*A. mellifera*), this result raises questions about the specific evolutionary history of the M-lineage, particularly regarding levels and spatial heterogeneity in gene flow between *A. mellifera iberiensis* and *A. mellifera mellifera*.

### Demographic modeling supports recent and massive secondary contacts

To shed light on the recent evolutionary history of honey bees, we performed demographic inferences between pairs of subspecies using DILS (Fraïsse et al. 2021), a computationally-efficient ABC modeling approach that accounts for the effects of linked selection. A total of 84 independent demographic analyses between pairs of genetic clusters, including constant vs. variable *N_e_* after the divergence of the two lineages (“Var. Ne”, Figure 2A) and simulations using or not the Site Frequency Spectrum (SFS, “SFS used”, Figure 2A) as summary statistics. Each of the 84 analyses were based on summary statistics calculated from independent samplings of 200 windows of 100-kbp of sequence.

**Figure 2:**
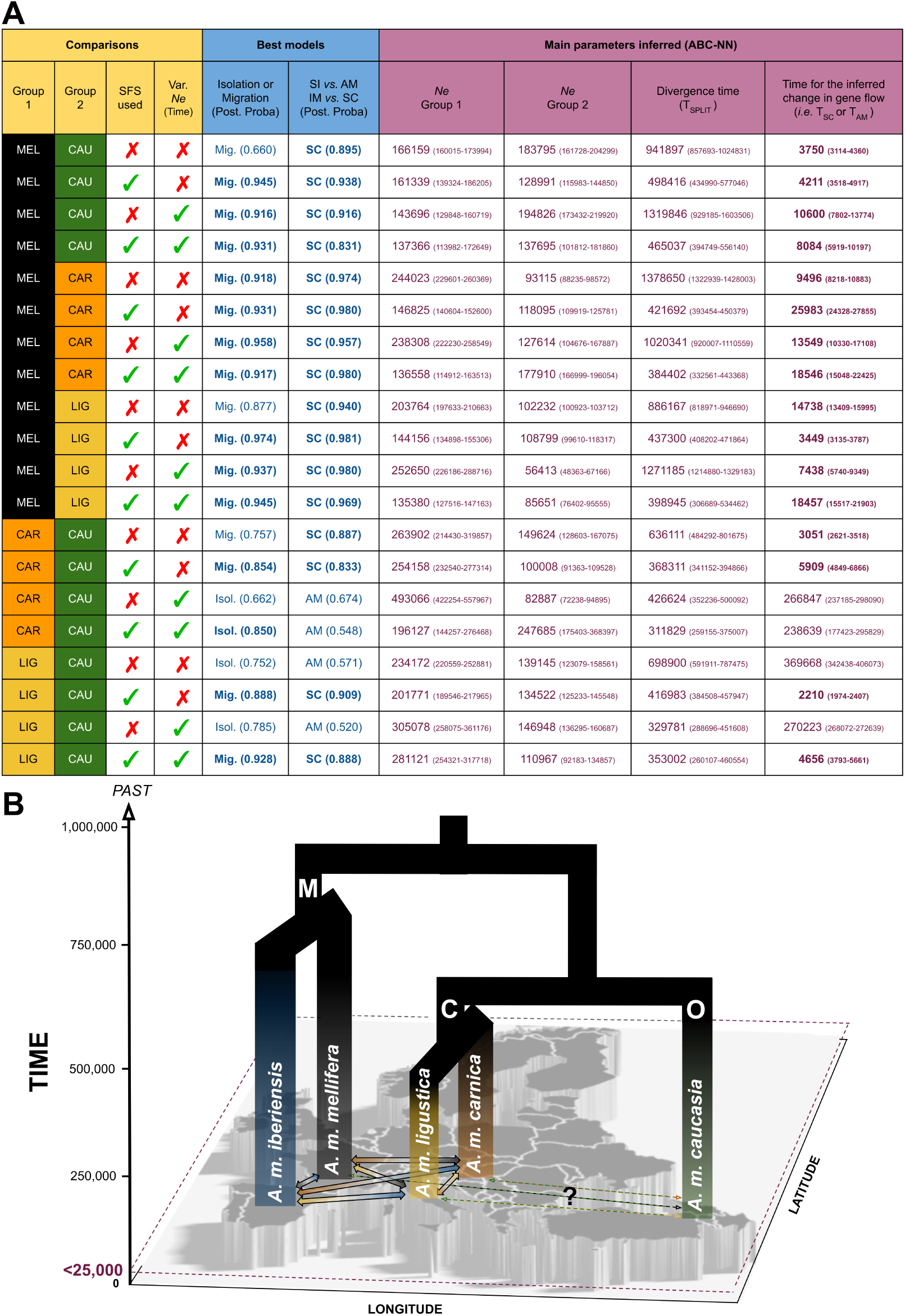
Summary of the ABC inferences. **A:** Best models and parameters for a series of inferences among subspecies from the 3 lineages (for extended information, see File S2). Models with posterior probabilities exceeding 0.80 are shown in bold. Abbreviations. ABC-NN: ABC Neural Networks (optimized set); AM: Ancient Migration; IM: Isolation with Migration; Isol.: Isolation; Mig.: Migration; Post. Proba.: Posterior Probability; SC: secondary Contacts; SI: Strict Isolation; Var. Ne: simulations assuming a temporal variation of Ne (e.g. bottleneck). **B:** schematic drawing of the most likely scenario of divergence for the Eurasian honeybees (lineages M, C and O) based on the ABC inferences, especially highlighting the recent postglacial secondary contacts (<25,000 generations). Time is expressed in generations past.

Model choice performed based on all these comparisons are consistent with ongoing gene flow between the different clusters (68/84 comparisons, including 47 with posterior probability exceeding 80%, Fig. 2A and File S2). Importantly, for all these comparisons supporting a scenario of ongoing gene flow, secondary contact scenarios perform better than those assuming isolation-with-migration (68/68, including 43 with a posterior probability exceeding 80%). Our demographic inferences provide considerable support for ongoing migration and, more specifically, secondary contacts among clusters from the C-and M-lineages, since all pairwise comparisons (32/32) support such a scenario (32/32, 30/32 and 18/32 with posterior probabilities exceeding 0.8, 0.9 and 0.95, respectively). Inferred time for secondary contacts (T_SC_) are mostly consistent with recent events (30/32, 20/32 and 13/32 with median T_SC_ estimates < 50,000, < 25,000 and <12,500 generations ago, respectively) between the M- and C-lineages (Fig. 2).

Among the 84 performed comparisons, the only 16 supporting no ongoing gene flow (including 8 with posterior probabilities > 0.8) almost exclusively include the Caucasian cluster (15/16). Ancient migration scenarios appear to better fit the data than strict isolation ones, albeit this comparison is poorly statistically supported (0/16 with posterior probabilities > 0.8). Surprisingly, some other simulations regarding comparison between the O-lineage and either the M-lineage or the C-lineage exhibit considerable support for ongoing migration (9/24, including 7/9 with posterior probabilities > 0.8) and secondary contacts (9/24, all with posterior probabilities > 0.8), especially for pairwise comparisons between Mellifera and Caucasia (4/4 supporting SC with posterior probabilities > 0.8, Fig. 2A), questioning the existence and intensity of gene flow with Caucasia (Fig. 2B).

Although variable among replicates, the inferred divergence times (T_SPLIT_) for the among-lineages comparisons are mostly consistent with the phylogenetic tree previously reconstructed (Fig. 1D), with more ancient median estimates for T_SPLIT_ for comparisons between the M-lineages on one hand, and the C- and O-lineages on the other hand (48 among-lineage comparisons, median=914,032 generations, mean=900,100 generations, SD=533,191 generations). Similarly, T_SPLIT_ was consistently inferred to be more recent for the comparisons between the C- and O-lineages (8 among-lineage comparisons, median=392,647; mean=442,693, SD=145,122). Our inferences are also consistent with a split between the two subspecies of the C-lineage (‘Carnica’ & ‘Ligustica’) occurring relatively soon after the divergence of the C- and O-lineage (4 within-lineage comparisons, median=324,590; average=383,735, SD=335,380). The divergence of the different M-lineages is inferred to be more ancient. Among the genetic groups corresponding to the M-lineage (‘Mellifera’, ‘Colonsay’, ‘Ouessant’ and ‘Iberiensis’), T_SPLIT_ was inferred to be ancient (24 within-lineage comparisons, median=693,002, mean=635,173, SD=408,235). Unlike the phylogeny (Fig. 1D), inferences are consistent with more recent divergence among the 3 clusters belonging to *A. mellifera* (12 comparisons, median=625,863, mean=598,006, SD=411,304) than between pairs of these 3 clusters of *A. mellifera* with *A. mellifera iberiensis* (12 comparisons, median=765,167, mean=672,340, SD=419,854; for details, see File S2).

To check the robustness of the demographic inferences to the effects of background selection, the analysis were also performed on 50 windows of 100-kbp that are specifically in low or high recombining regions. Indeed, this background selection is expected to negatively impact the performance of demographic inference methods (Pouyet et al. 2018). These results were remarkably consistent with the previous results, reinforcing the support for recent postglacial secondary contacts between the M- and C-lineages (lowly-recombining: 28/32 supporting SC, including 27/28, 15/28 and 4/28 with posterior probabilities exceeding 0.8, 0.9 and 0.95, respectively; highly-recombining: 32/32 supporting SC, including 32/32, 25/32 and 3/32 with posterior probabilities exceeding 0.8, 0.9 and 0.95). The secondary contacts were similarly found to be extremely recent for both the inferences in low recombination regions (25/28, 20/28 and 14/28 with median T_SC_ estimates < 50,000, < 25,000 and <12,500 generations ago, respectively) and high recombination regions (30/32, 27/32 and 18/32 with median T_SC_ estimates < 50,000, < 25,000 and <12,500 generations ago).

### Investigating global nucleotide diversity levels and impacts of recent demography

Using non-overlapping 100-kbp sliding windows spanning the genome, we reported variable levels of nucleotide diversity (*π*) among the different species and genetic clusters (Fig. 3A). Lower levels of diversity were observed among the samples from the two giant honey bee species (median=1.85×10^-3^ and 1.45×10^-3^ for *A. dorsata* and *A. laboriosa*, respectively) as compared to those from *A. cerana* (2.48×10^-3^) and *A. mellifera* clusters (1.91-3.48×10^-3^). Within *A. mellifera*, the lowest diversity was observed in the two conservatories of the black honey bee (*A. mellifera mellifera*) located on Atlantic islands (Colonsay, Scotland: 1.91×10^-3^; Ouessant, France: 1.94×10^-3^). Higher levels of diversity were observed for the ‘Iberiensis’ (*A. m. iberiensis* samples, 2.70×10^-3^) and ‘Mellifera’ genetic clusters (2.66×10^-3^). The latter group corresponds to a genetic cluster composed of samples from two Mediterranean conservatories in Southeastern France: Porquerolles island and the nearby continental conservatory at Solliès. Among samples from the C-lineage, we reported higher levels of diversity in the Carnica (2.77×10^-3^) than in Ligustica (2.13×10^-3^). Within the two Carnica subgroups, we reported a marked difference in nucleotide diversity between non-German (2.75×10^-3^) and German Carnica (2.24×10^-3^). The Caucasia genetic cluster (O-lineage), composed of samples that were collected in France from importers, exhibit one of the weakest diversity levels (1.99×10^-3^). Two genetic clusters assumed to be associated to to two recent breeds, respectively selected for royal jelly (‘RoyalJelly’) and honey production (‘Buckfast’), surprisingly exhibit the two highest levels of nucleotide diversity (‘RoyalJelly’: 2.85×10^-3^; ‘Buckfast’: 3.84×10^-3^).

**Figure 3:**
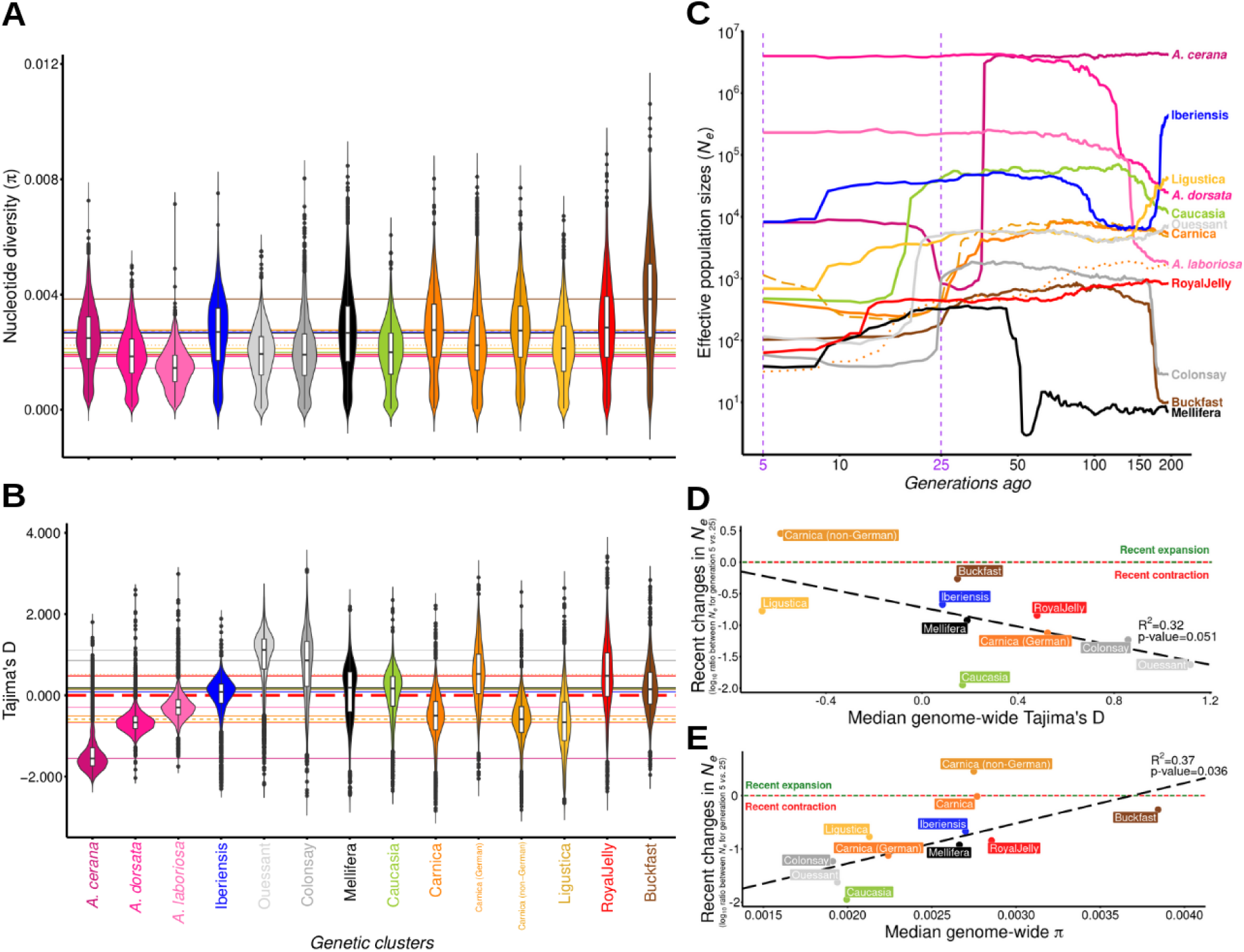
Genetic diversity and impacts of recent demographic changes. **A.** Per-population distributions of nucleotide diversity estimates in non-overlapping 100-kb sliding windows spanning the honeybee genome. **B.** Tajima’s D estimates for the same windows. **C.** Linkage disequilibrium (LD)-based inferences of effective population sizes (N_e_) over the last 200 generations as inferred with GONE. Median values over 40 GONE replicates are shown (see Fig. SX for details). The dotted and dashed orange lines correspond to the German and non-german samples from the Carnica genetic group, respectively. **D & E.** Scatterplots and linear correlations between the ratio of GONE-inferred N_e_ for generation 5 vs. generation 25 (as shown in purple in C) as compared to the median of Tajima’s D (**D**) and nucleotide diversity (**E**) across the whole genome for all A. mellifera genetic clusters. Recent changes were calculated as log_10_(N_e(5gen)_/N_e(25gen)_). The entire Carnica population (including both German and non-German samples) was excluded prior to performing the linear tests.

To understand recent demographic changes, we used two independent strategies: Tajima’s D, a population genetic summary statistic that contrasts π and θ to identify genome-wide departures from mutation-drift equilibrium, such as bottlenecks and expansions (with positive and negative D values, respectively), and GONE, a linkage disequilibrium-based method that infers effective population sizes (*N_e_*) from linked markers, revealing *N_e_* changes over up to 200 generations (Santiago et al. 2020). Median of Tajima’s D across all the 100-kbp windows highlighted negative Tajima’s D values over the three other *Apis* species: *A. laboriosa* (-0.30), *A. dorsata* (-0.67) and *A. cerana* (-1.55, Fig. 3B). Among the *A. mellifera* genetic clusters, median Tajima’s D were highly variable, ranging from -0.67 to 1.12 (Fig. 3B). Most positive Tajima’s D values are observed in the two Atlantic conservatories of black honeybee (Colonsay, Scotland: 1.12; Ouessant, France: 0.85), the clusters that already exhibited the lowest levels of nucleotide diversity (Fig. 3A). In contrast, the other M-lineage genetic clusters exhibit less positive values, respectively 0.09 and 0.19 for Iberiensis (Continental Spain) and Mellifera (Southeastern France). As a contrast, most negative values observed for clusters from the C-lineages (Ligustica, -0.67) and Carnica (-0.50), albeit with a profound difference when German (0.52) and non-German Carnica samples (-0.59) are contrasted (Δ_Tajima’s D_ = 1.11). For recent breeds, we observed slight to moderate positive values of Tajima’s D for both ‘RoyalJelly’ (0.48) and ‘Buckfast’ (0.15), consistent with the general expectation of population contraction in breeds. A slightly positive median Tajima’s D value was also reported for Caucasia (0.17), which corresponds to Caucasian samples recently imported to France due to the current interest in this genetic lineage among French beekeepers producing honey, a value which therefore appears to be relatively similar to that reported for ‘Buckfast’.

Inferences of *N_e_* changes over the last 200 generations, as inferred with the approach implemented in GONE (Santiago et al. 2020) revealed important trajectories among samples (Fig. 3C). Based on median values of *N_e_* over 40 independent GONE runs for each cluster (see Sup. Note 2 for details), we reported (Fig. 3C) expansions in both *A. dorsata* and *A. laboriosa* occurring between 100 and 200 generations ago, followed by around 100 generations of constant population size. In *A. cerana*, the inference is more consistent with a massive drop in *N_e_* around 50 generations ago, followed by a partial recovery that started around 25 generations ago. More interestingly, among the *A. mellifera* genetic clusters, important differences in trajectories were also reported (Fig. 3C). Regarding the M-lineage clusters, considerable support for recent and massive bottlenecks were inferred for the two conservatories in the Atlantic islands (Colonsay, Scotland & Ouessant, France), occurring around 25 and 20 generations ago, respectively. As compared to these two conservatories, Mellifera, which correspond to two spatially close Mediterranean conservatories in Southeastern France, exhibit a very different trajectory, albeit more variable among the different GONE runs than for the other clusters (see Sup. Note 2), with a relatively long evolution at low *N_e_* (between generations 200 to 50 generations ago), followed by a sudden increase in *N_e_* around 50 generations ago and a slight but constant bottleneck afterward. As compared to the other M-lineage, Iberiensis exhibits larger effective population size, with relatively limited changes in *N_e_*, at least over the last 100 generations. Among the two genetic clusters from the C-lineage, we identified clear bottlenecks over at least the last 100 generations. In Carnica samples, we observed two distinct trajectories between the German (dotted orange line, Fig. 3C) and non-German samples (dashed orange line), with clear evidence of a contraction and an expansion, respectively. The divergence in these trajectories appears to have begun approximately 15 generations ago.

Although initial introductions may have occurred since the late 19th century, the globalization of beekeeping, marked by the massive imports of C-lineage bees for commercial breeding, primarily took place after World War II, with significant intensification around the 1980s. To investigate the impacts of modern beekeeping on effective population sizes, we considered the log_10_-ratio of changes in GONE-inferred *N_e_* between generations 5 and 25. We used generation 5 to exclude potential remaining recent kinship in the data. Given that most samples were collected in 2014-2015, and assuming a generation time of 3 years in modern breeds (queen lifespan: 2-5 years), our summary statistic provides a contrast between inferred *N_e_* around the years 1940 and 2000. Marginally significant negative correlation (p=0.051) between Tajima’s D and this latter summary statistic suggests that it captures well the genomic impacts of demography across this period of time. Overall, all clusters exhibit signatures consistent with contraction, albeit variable in intensity, with the notable exception of non-German Carnica for which the signature is more consistent with an expansion across the time frame (Fig. 3D). In detail, we observed marked negative values for yellow honey bees (C-lineage) which are genetic backgrounds or breeds widely used in modern beekeeping such as German Carnica (-1.12), ‘RoyalJelly’ (-0.84), Ligustica (-0.77). Black bees have also been particularly impacted. Albeit the most intense contraction were observed in the two conservatories of black bees from Atlantic islands of Colonsay, Scotland (-1.23) and Ouessant, France (-1.62), the two other genetic clusters associated with black honey bees (C-lineage) also exhibit marked negative values (Mellifera: -0.92; Iberiensis: -0.67). Across the whole dataset the population that exhibit the strongest contraction over the period is Caucasia (-1.95).

We subsequently tested whether the GONE-associated metric capturing changes in *N_e_* during the period of intensification and globalization of beekeeping, is consistent with the observed differences in present-day nucleotide diversity levels (Fig. 3E). Remarkably, we found a significant positive linear correlation (R^2^=0.37, p-value=0.036, Fig. 3E) between recent changes in *N_e_* and the nucleotide diversity levels, suggesting that modern beekeeping has had a substantial impact on present-day nucleotide diversity. The most striking evidence is the difference between German Carnica samples, which have diverged from non-German ones over just 15 generations (Fig. 3C), yet already exhibiting profound differences in their levels of diversity (Fig. 3A and 3E).

### Conserved genomic landscapes of nucleotide diversity among *Apis*

Investigating variation of population genomic summary statistics along the genome (hereafter referred to as genomic landscapes) is known to be crucial to understand how the evolutionary processes, such as natural selection and recombination, have shaped genetic variation across the genome. By first investigating the correlations of the genomic landscapes of nucleotide diversity (Fig. 4A), we uncovered extremely high correlations among the different *A. mellifera* genetic clusters (median=0.776; min=0.628 (Ligustica-Colonsay); max= 0.922 (Ouessant-Iberiensis)). More surprisingly, we also revealed particularly high correlations for the among-species comparison (median=0.633; min: 0.534 (*A. laboriosa*-Colonsay); max: 0.907 (*A.laboriosa*-*A.dorsata*)), revealing a high correlation of the genomic landscapes across a long evolutionary timescale. Despite the significant variation in observed diversity between low and high recombination regions of the genome (Fig. S3, see next section), the correlation of genomic landscapes remains comparably well-recovered in both low and high recombination regions (Fig. S4). A neighbor-joining tree based on the correlation matrix of nucleotide diversity (Fig. 4B) captures the long-term history of divergence among the species and subspecies, with remarkable consistency with our earlier analyses (Fig. 1, Fig. 2A), including the sequence of splits among the different M-, C- and O-lineages.

**Figure 4:**
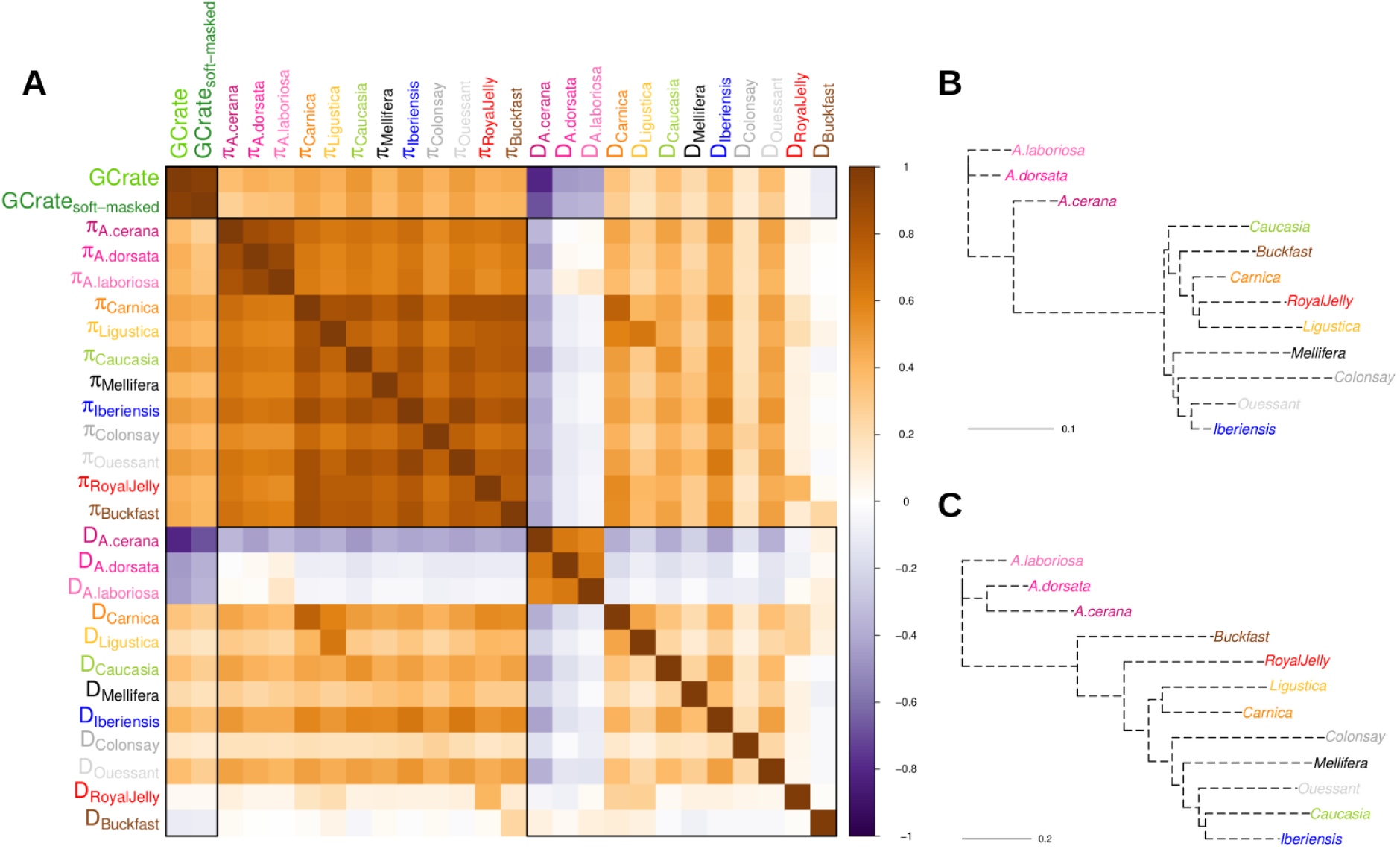
Correlations of genomic landscapes of nucleotide diversity, Tajima’s D and recombination. **A.** Correlation matrix of the genomic landscapes of nucleotide diversity and Tajima’s D among the different species and subspecies A. GC%, π and Tajima’s D values are based on non-overlapping 100-kbp genomic windows spanning the whole genome. GC % was calculated on the A. mellifera reference genome (HAv3.1) and is used as a proxy of the recombination rate. GCrate_soft-masked_ corresponds to the GC% in soft-masked regions only. See also Fig. S4 and S7 for correlations of π and Tajima’s D docussing on highly or lowly recombining regions. **B & C.** Neighbor-joining trees using 1-covariances of π (**B**) and Tajima’s D (**C**) as distance matrices. Trees are rooted on the ancestors of A. laboriosa and A. dorsata (B) or A. laboriosa, A. dorsata and A. cerana (C).

Similarly, we investigated the correlations of the genomic landscapes of Tajima’s D (Fig. 4A). Globally, we observed weak negative correlations of the Tajima’s D landscapes between *A. mellifera* and the other species (median=-0.110, min=-0.429 (*A.cerana*-Iberiensis), max=0.097 (*A.laboriosa*-Buckfast)), but remarkably strongly positive correlations among the three other species (median=0.632; min=0.575 (*A.laboriosa*-*A.cerana*), max=0.639 (*A.dorsata*-*A.laboriosa*)). The singularity of the genomic landscapes of *A. mellifera* as compared to the other species seems mostly driven by low recombination regions, in which Tajima’s D are mostly negative in *A. mellifera* while they are mostly positive in the other species (Figs. S5-6), such that the landscapes become either entirely uncorrelated or slightly positively correlated when considering only high recombination regions (Fig. S7). Comparing genomic landscapes among the *A. mellifera* genetic clusters, Tajima’s D appears substantially correlated median=0.162; min=-0.074 (Mellifera-Buckfast); max=0.490 (Ouessant-Iberiensis)), albeit less correlated than those of nucleotide diversity, especially in highly recombining regions (Fig. S7). Tajima’s D genomic landscapes are even more correlated, when the recent breeds are not considered (median=0.256; min=0.087 (Colonsay-Ligustica); max=0.490 (Ouessant-Iberiensis)). Indeed, Tajima’s D genomic landscape of Buckfast is slightly negatively correlated with all clusters (median=-0.030), except with those from the C-lineage (RoyalJelly=0.021, Ligustica=0.084, Carnica=0.104). Albeit positive, the correlation coefficients are similarly low for ‘RoyalJelly’ (median=0.061), except when compared with Carnica (0.263) and Ligustica (0.272). Tajima’s D genomic landscapes of Caucasia (O-lineage) are surprisingly highly correlated with both Iberiensis (0.484) and Ouessant (0.388), such that Caucasia falls within the M-lineage on a Neighbor-joining tree based on the correlation matrix of Tajima’s D (Fig. 4C).

### Recombination shapes highly heterogeneous conserved landscapes of nucleotide diversity

To better understand the genomic landscapes of nucleotide diversity, we investigate whether the recombination rate could have been one of the key evolutionary processes at play in shaping that variation. First, we investigated the correlations of the GC content and levels of diversity over all 100-kbp windows spanning the whole genome. Base composition in DNA is indeed associated with recombination rates, as a by-product of GC-biased gene conversion (gBGC), a recombination-associated meiotic repair bias favoring G and C over A and T alleles at recombination sites (Duret & Galtier, 2009). We observed strongly positive correlations between the genomic landscapes of GC content and diversity for all genetic clusters (median=0.447, with min=0.402 and max=0.519 for Mellifera and Caucasia, respectively, Fig. 4A, see also Fig. S3). Second, we benefited from the uniquely high recombination rates in honey bees (*e.g.* Beye et al. 2006; Kent et al. 2012; Liu et al. 2015; Wallberg et al. 2015; Conlon et al. 2023) to visually compare empirical estimates of recombination rates estimated from a genetic map (internal circle, Fig. 5) with the diversity levels of the different genetic groups. Although less precise than using GC content due to the necessity of larger window sizes resulting from the limited number of sequenced individuals (Liu et al. 2015), this approach still allows us to more directly reveal the link between the recombination and diversity landscapes.

**Figure 5:**
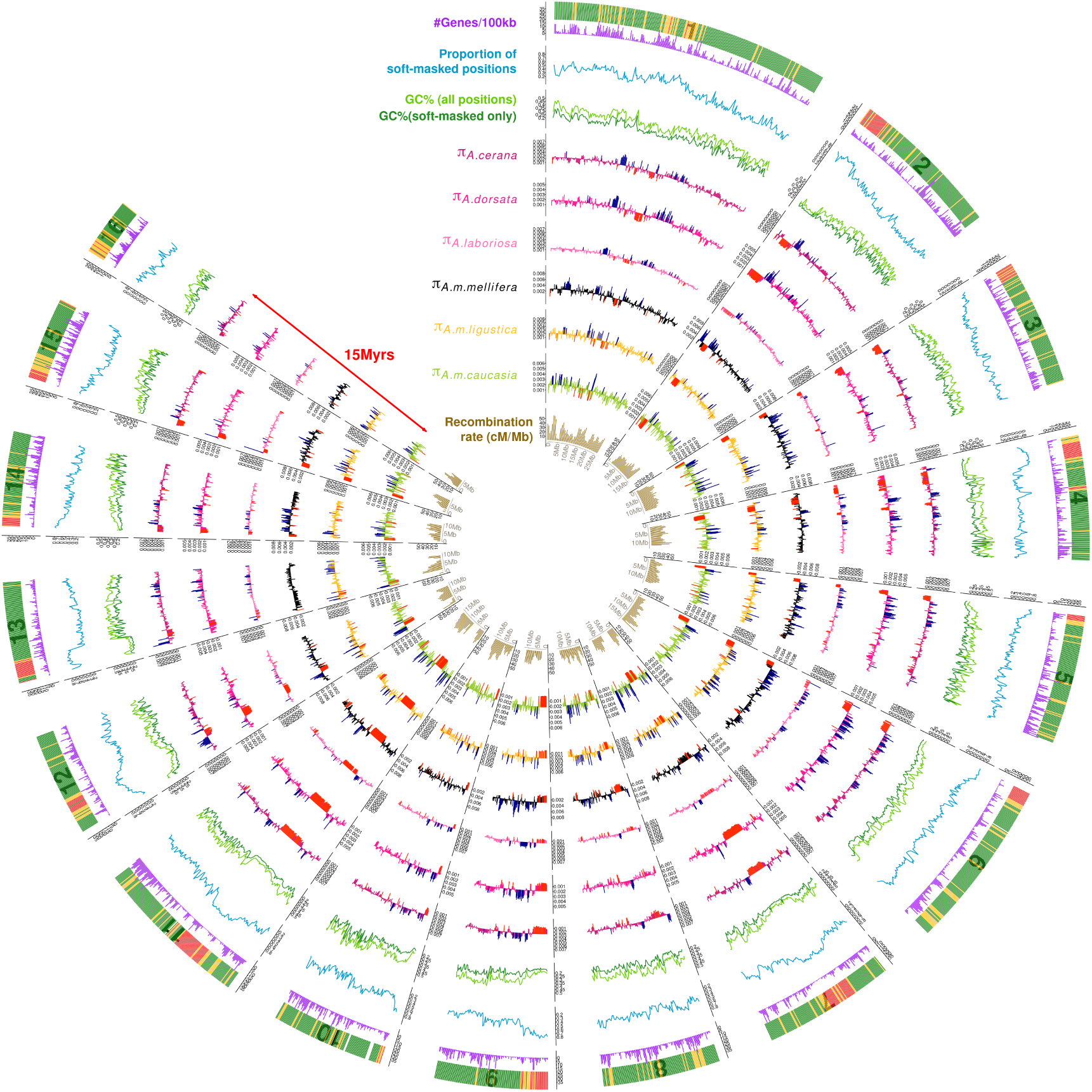
Circular visualization of the correlated genomic landscapes of recombination and diversity in Apis, covering more than 15 million years of divergence. From external to internal: gene density, proportion of soft-masked position on the reference genome, GC-content and GC-content (dark green) at soft-masked (green) positions only, nucleotide diversity in A. cerana, A. dorsata, A. laboriosa, Mellifera, Ligustica and Caucasia and recombination rate (cM/Mb). The most external track highlights the genomic windows that are consistent with high-recombination (green), intermediate (orange) or low-recombination (red). The chromosome number is also indicated, from 1 to 16

Even more remarkably, we not only observed high correlations of the GC content in the reference genome of *A. mellifera* with the nucleotide diversity of the different *A. mellifera* genetic clusters, but also with the diversity landscapes of the other *Apis* species, reaching 0.361, 0.405 and 0.423 for *A. cerana*, *A. laboriosa* and *A. dorsata*, respectively (Fig. 4A). Given that GC content was solely estimated based on the reference genome of *A. mellifera*, such correlations suggest that the recombination landscapes have evolved little over the last 15 million years. To investigate whether such a hypothesis is plausible, we evaluated the conservation of gene order at orthologous genes (1:1 copies) as well as the GC content (Fig. S8A). We also performed whole-genome alignments of *A. cerana* (ACSNU2.0, Park et al. 2015) and *A. mellifera* reference (HAv3.1, Wallberg et al. 2019) assemblies with D-genies to investigate large-scale synteny (Fig. S8B). Our analysis is consistent with a near-perfect collinearity of the two genomes (see also Wang et al. 2020 for similar results), with few potential inversions. Given that the other *Apis* reference genomes remain quite fragmented (e.g. *A. laboriosa*, Cao et al. 2023), we evaluated the level of synteny between *A. mellifera* and *Bombus terrestris*, being the closest well-annotated whole genome sequence. Consequently, we investigated the synteny of orthologous genes using the method implemented in Genespace. Despite that the divergence time between bumblebees and honey bees is estimated to be around 82 million years ago (CI=77-85, ToL), we observed remarkable large-scale synteny (Fig. S8C), with little evidence for major inter-chromosomal rearrangements, with the notable exception of the chromosomes 1 and 7 of *B. terrestris*, which are fused in *A. mellifera* (chromosome 1), suggesting either a Robertsonian fusion in *A. mellifera* lineage, or a centric fission in the *Bombus* lineage. We therefore observed a remarkable one-to-one correspondence between the chromosomes, with evidence only for within-chromosome inversions. Taken together, our results suggest that recombination is a major driver of whole-genome variation in nucleotide diversity, and that the minimal reshuffling of genomes has contributed to maintaining conserved recombination landscapes, thereby maintaining landscapes of nucleotide diversity over long evolutionary timescales.

## Discussion

### Tracking long-term divergence of honey bees

Combining phylogenetic approaches with demographic modeling, we reconstructed the sequences of speciation events between four different *Apis* species, as well as several *Apis mellifera* subspecies. The two approaches are complementary, since demographic modeling has very little power to infer very ancient events (>5-10 *N_e_* generations ago, *i.e.* 1.5-3 million generations ago, assuming 300,000 haplodiploid individuals as the upper boundary for the post-T_SPLIT_ *N_e_* as inferred in DILS (File S2, for theoretical expectations regarding *N_e_*, see Charlesworth, 2009 among others). Consequently, we reconstructed phylogenetic trees offering a window on ancient events, such as the divergence among the different species. Consistent with the literature, our reconstruction provides considerable support (bootstrap=100%) for a first split between the two giant honey bees species *A. dossata* and *A. laboriosa* on one side and the ancestor of Eastern *A. cerana* and Western *A. mellifera* honey bees on the other, an event which is assumed to have occurred more than 15 million years ago (Engel 2006; Ramírez et al. 2010; Martins et al. 2015). Since this initial event, as seen on our phylogenetic trees and according to expectations, the split between the Eastern and Western honey bees predates that between the two giant honey bee species. This is consistent with the long taxonomical debate regarding the best species delimitations in giant honey bees (Ruttner, 1988).

When looking at more recent time periods, we compared the sequences of splits between subspecies within *A. mellifera* as reconstructed through the phylogeny from the demographic inferences. Both analyses are consistent with a first split between the Western European (M) lineage and the ancestor of the Central (C) and Eastern (O) lineage, consistent with the literature (Han et al. 2012; Wallberg et al. 2014; Dogantzis et al. 2021). Regarding the timing, the event is inferred to have occurred around 900k generation ago, an estimation which is difficult to scale in real time due to the uncertainties regarding the generation time in natural habitats. Assuming a generation time of 1 year in natural environments (as assumed by Wallberg et al. 2014), such a divergence time of 0.9 Myr is relatively consistent with the divergence times of 0.7 and 1.0 Myr calculated by Arias & Sheppard (1996) and Garnery et al. (1992) and, more broadly, falls in the range of divergence time values generally assumed for the main honey bees lineages, from 0.30-1.00 Myr (Garnery et al. 1992; Arias & Sheppard 1996; Wallberg et al. 2014), but departs from the 5.54 Myr estimate provided by Dogantzis et al. (2021). As compared to the split within the M-lineage, the divergence of the C and O-lineages appears to be far more recent, inferred to have occurred around 450k generations. The ABC-reconstructed relationships among lineages (Fig. 2A) is not only consistent with the phylogenetic tree (Fig. 1), but also with the literature (Han et al. 2012; Wallberg et al. 2014; Dogantzis et al. 2021), therefore (i) consolidating the general backbone for the different *A. mellifera* lineage and (ii) building confidence in the ABC-based inferences. Within the M-lineage, we inferred relatively ancient splits between the four clusters studied here, going up to 675k generations BP. Given the wide distribution of the M-lineage in Europe, covering a large region spanning from Spain (*A. m. iberiensis*), all of Western and Northern Europe including Scandinavia, as well as Western Russia (*A. m. mellifera*), one might expect that the M-lineage has more complex history than previously reported, in which different groups could have been isolated in different locations during glacial periods. Demographic reconstruction accounting for the past periods of gene flow, as well as other sources of confounding factors, could have contributed to provide robust estimates of divergence times. Altogether (Fig. 1, 2, but see also Fig. 4), our analysis suggests that the evolutionary history of the M-lineage could be more complex than commonly thought. Regarding the C-lineage and contrary to Wallberg et al. 2016 for which the inferred split between Carnica and Ligustica was inferred to be fairly recent (∼25,000 years BP), our analysis is consistent with more ancient divergence times between the two C-lineages Ligustica and Carnica (∼380k generations BP). Such a time frame appears more consistent with the hypothesis of a divergence at the onset of a glacial period, while the estimate by Wallberg and collaborators (2014) requires that the honey bees have only started to diverge in the middle of the last glacial period, at a time very close to the last glacial maximum (21,000-23,000 BP). A more ancient divergence would be more consistent with the general hypothesis that divergence occurred at the beginning of an ice age period, the latest one or one of the former ice age periods. However, our analysis also suffers from important limitations, including the fact that intercrossing between the O- and C-lineages are very popular among French beekeepers. Although admixed individuals (Q-value < 0.9) were excluded from our analysis, historical introgression events (e.g. 7-10 generations ago) could have remained undetected by Admixture, which could otherwise impact the inferences of the divergence time between Carnica and Ligustica. This hypothesis could be consistent with the phylogeny, since we observed Carnica to be intermediate between Caucasia and Ligustica, with paraphyly in the Carnica group and almost no bootstrap support at the main nodes of the C-lineage (Figs. 1D and S2).

### Secondary gene flow in honey bees predates human disturbances

Modern beekeeping is associated with massive importation of non-local bees, as a consequence, recent admixture among the honey bees, especially between the C- and M-lineages have been widely reported (*e.g.* Harpur et al. 2012; Wallberg et al. 2014; Henriques et al. 2020; Wragg et al. 2022). Using an approach based on the demographic reconstruction of different subspecies pairs, we found that among-lineage gene flow likely precedes the beekeeping-associated transfers of honey bees and is more consistent with naturally occurring secondary contacts. Of course, there remain significant uncertainties inherent to demographic modeling approaches regarding these datings; nonetheless, most estimates are consistent with secondary contact occurring between 1,500 and 25,000 generations ago. While it is generally assumed that queens can live up to five years under controlled conditions in modern beekeeping (Prado et al. 2020), which includes specific management practices like swarm control, feeding, and providing and maintaining ample space for brood nests, there is considerable uncertainty about the generation time of honey bees in natural conditions. Considering both a generation per year as assumed by Wallberg and collaborators (2014) and the fragmented geography of Europe, which favors periods of allopatric isolation in ice age refugia, we consider these estimates to be consistent with post-glacial secondary contacts occurring after the last glacial maximum. Similarly to honey bees, robust statistical evidence for secondary contacts following the last glacial maximum has emerged from demographic modeling studies over the past decade. Notable examples include European white oaks (Leroy et al., 2017; 2020) and European sea bass (Tine et al., 2014; Duranton et al., 2018). These findings are in line with the expected generic nature of postglacial secondary contacts for European biodiversity, given the continent’s unique geography. During the ice ages, lineages diverged in allopatry in different Southern refugia and came into secondary contact after postglacial recolonization (Hewitt, 1999; 2004; Bierne et al. 2011).

### Compelling evidence of modern beekeeping’s influence on present-day diversity

The impact of modern beekeeping on the evolution of honey bee genetic diversity remains a topic of significant debate among evolutionary biologists (Harpur et al. 2012; 2013; De la Rua, 2013; Mikheyev et al. 2015; Espregueira Themudo; 2020; Parejo et al. 2020). Resolving this issue is however crucial for the long-term preservation of honey bees and their vital role in biodiversity and agricultural pollination. To accurately assess changes in genetic diversity, we first excluded admixed individuals from the dataset (Fig. 1), as their temporarily elevated diversity could obscure the long-term trends in genetic variation (De la Rua, 2013). By generating unbiased estimates of nucleotide diversity, Tajima’s D, as well as performing inferences of recent changes in effective population sizes, we found strong evidence for widespread declines of both *N_e_* and nucleotide diversity in honey bees, such that we observe remarkable correlation between the intensity of recent bottlenecks associated with modern beekeeping with present-day honey bee diversity (Fig. 3E).

Focusing on the most managed populations, including *A. m. ligustica*, *A. m. carnica*, *A. m. caucasia*, as well as the ‘Buckfast’ and ‘RoyalJelly’ breeds, we inferred temporal declines of *N_e_* and positive Tajima’s D. Upon identifying within-population structure among our Carnica samples (Fig. S1), which are near-perfectly consistent with samples collected in Germany and outside Germany, we also observed a marked lower diversity among German samples, a pattern that aligns with evidence of a recent bottleneck unique to German Carnica (Fig. 3). German beekeepers have implemented highly effective breeding programs, leveraging controlled queen fertilization through isolated mating stations and artificial insemination (Hoppe et al. 2020), which seem to directly translate into higher median kinship values (Fig. 1C). These methods have however proved their effectiveness for beekeepers with reported genetic gain in some specific traits (Hoppe et al. 2020). Finally, and counterintuitively, the ‘Buckfast’ and ‘RoyalJelly’ breeds are the two genetic clusters exhibiting the highest diversity (Fig. 3A), especially in ‘Buckfast’, a group of hybrid origin (Adam, 1986). This could appear consistent with the conclusions of Harpur and collaborators (2012) suggesting that bee management by humans have contributed to increase genetic diversity of honey bees. It should however be noted that this higher diversity is likely transient. Our result indeed supports the hypothesis of a genetic shrinkage through relatively high median kinship (Fig. 1C), slight to moderate decline in *N_e_* in both lineages, as well as positive tajima’s D values (Fig. 3).

Conservation programs for black honey bees are crucial to preserve the specific genetic makeup of the M-lineage. Indeed, large census population sizes of C-lineage associated with modern beekeeping induce a main risk of massive introgression from the C-to M-lineage, threatening the long-term genetic integrity of the black honey bee genetic background. Recently, Wragg et al. (2022) reported on the effectiveness of conservatories, especially those located on isolated islands, such as Ouessant, Porquerolles and Colonsay, in maintaining pure genetic backgrounds. Our analysis however highlights a fundamental issue with such island reserves. Small islands can only host a limited number of hives (e.g. ∼100 colonies in Ouessant, France), which therefore induce a strong bottleneck at the setting up of the conservatory, followed by snowballing of inbreeding depression. We found considerable support for both the initial bottleneck, which was inferred to be more ancient in Colonsay, Scotland than in Ouessant, France, in line with the difference in the establishment date of the two conservatories. This empirical evidence highlights the high resolutive power of the LD-based approaches to infer recent changes in *N_e_*, such as the one implemented in GONE (Santiago et al. 2020). Given the particularly poor situation across all our indicators, from the high level of relatedness to the evolution of diversity, which is the lowest among all clusters, new initiatives must be undertaken. It is very clear that the current situation does not allow for the long-term preservation of the genetic diversity of black bees in the conservatories. To improve the situation, it is essential to establish European queen exchange programs between conservatories to reintroduce genetic diversity while preserving genetic purity. However, significant effort is also needed to better understand the demographic history of black bees, which appears more complex than previously described, such that these different sub-lineages could be then considered for exchange programs in order to better preserve the black bee diversity as a whole.

### Recombination stability induces long-term conservation of genomic landscapes of diversity

Recombination rates are well characterised as exceptionally high in honey bees, with estimates per generation ranging from 19 to 37 cM/Mb, *i.e.* around 20 times higher than in humans (*e.g.* Beye et al. 2006; Kent et al. 2012; Liu et al. 2015; Wallberg *et al*. 2015; Conlon et al. 2023). In addition to being elevated on average, the genetic landscape is also remarkably heterogeneous along the chromosomes. For instance, long non-recombining regions can be identified from direct evidence such as empirical genetic maps, or by using proxies of recombination rates such as tracks of introgression in hybrids (Wragg et al. 2022) or GC content (Fig. 5; Wallberg et al. 2015). By generating unbiased estimates of nucleotide diversity and GC content within the same window boundaries, we investigated the correlations between recombination and diversity landscapes. Our analysis revealed exceptionally high correlation coefficients, indicating that recombination plays a pivotal role in shaping the diverse genomic landscapes. Specifically, variable recombination rates contribute to the heterogeneous patterns of genomic *N_e_* through varying levels of linked selection, primarily linked to background selection but possibly also influenced by additional selective sweeps. This process collectively shapes the global nucleotide diversity landscape. Reports of correlated landscapes of recombination and nucleotide diversity have accumulated over the last decade, in a large range of species including birds (Burri et al. 2015; Vijay et al. 2016; Van Doren et al. 2017) and trees (Apuli et al. 2020; Shang et al. 2023).

One current trend in evolutionary biology is to track the progression of genomic landscapes over time through comparative studies of population or species pairs along a divergence gradient (Stankowski et al. 2019; Shang et al. 2023). Although this approach was not directly applied here due to the limited availability of population- and species-level genomic resources, we laid the groundwork by examining whether genomic landscapes of diversity are correlated among four *Apis* species. Remarkably, we found high correlation coefficients among all the different landscapes, despite a divergence spanning 15 million years within this genus. Furthermore, we found that GC content in *A. mellifera* was a strong predictor of the diversity landscapes in the other three species (Fig. 4). To explain the long-term conservation of these diverse landscapes, recombination rates must be conserved both at large and fine genomic scales over this long evolutionary timespan. In honey bees, both levels of conservation are likely. First, at large scale, we observed perfect or near-perfect collinearity of the *A. cerana* and *A. mellifera* reference genomes (Fig. S8, see also Wang et al. 2020). Lacking high-quality reference genomes for *A. laboriosa* and *A. dorsata*, we explored whether high collinearity might be a general feature of the clade by comparing the synteny between the two best-assembled genomes within the Apinae subfamily: *Bombus terrestris* (the buff-tailed bumblebee) and *Apis mellifera*. This comparison spans approximately 82 million years of divergence and revealed near-perfect chromosome-by-chromosome correspondence. Although recombination rates vary within Apinae (Kawakami et al. 2019), our findings support the hypothesis that large-scale recombination landscapes are maintained through the preservation of large syntenic blocks. Second, at a finer scale, the stability of genomic landscapes is influenced by the evolution of recombination hotspots. In mammals and many other vertebrates, the PRDM9 gene has been shown to play a crucial role in the rapid turnover of fine-scale recombination maps (Smagulova et al. 2016). Contrary to these species, Honey bees lack a functional PRDM9 gene (Wallberg et al. 2015). In some species lacking PRDM9, including finches (Singhal et al. 2015) and yeast (Lam and Keeney 2015), the absence of PRDM9-dependent recombination has been demonstrated to contribute to the conservation of fine-scale recombination landscapes over long evolutionary time scales. In summary, long-term conservation of heterogeneous recombination landscapes is likely in honey bees and would have contributed to shape their conserved heterogeneous landscapes of nucleotide diversity.

### Limitations of our study

The main limitation of our study lies in the modest overall sample size. Despite analyzing over 300 whole-genome sequences, the number of individuals sampled remains limited, which may affect our ability to provide a comprehensive overview of genetic diversity of honey bees at the continental scale. Collecting bees is a time-consuming task, and despite efforts to gather diverse samples from each genetic cluster, our study relied on a limited number of bee providers, including professional beekeepers and queen importers. This limitation should be considered when interpreting our findings. For instance, all *A. m. caucasia* samples were collected in France and likely represent recently imported genetics from a single honey bee importer from the Caucasus. As a result, the low genetic diversity and bottleneck signals observed in our Caucasia genetic clusters may not fully reflect those found in a larger sampling of *A. m. caucasia* from its native range. Results should be interpreted cautiously since similar considerations may apply to some other genetic groups in our study.

Similarly, demographic modeling is known to be an especially challenging task, for which many confounding factors can induce bias and therefore misleading results. In the present study, we have tried to control this risk as much as possible by using DILS, an ABC approach that allows modeling heterogeneity in effective migration rates and population sizes along the genome in order to account for the presence of species barriers and linked selection, two known sources of incorrect inferences (Roux et al. 2014; Ewing & Jensen, 2016; Schrider *et al*. 2016; Pouyet et al. 2018; Johri *et al*. 2020). DILS can also model non-constant evolution of *N_e_* during the divergence of each population, which represents another source of bias (e.g. Momigliano *et al*. 2021). Although we made considerable efforts to account for most impactful biases, our approach still has limitations. In particular, given the complexity of each model, the actual demographic history is likely far more complex than what we have modeled. Specifically, DILS does not allow demographic inferences for more than two populations at a time, and it is entirely possible that gene flow from third-party species could disrupt inferences for certain pairs. As a result, the overall scenario we produced by comparing results across different pairs (Fig. 2B) should be interpreted with caution. Despite the fact that the main findings have proven to be remarkably consistent among most of the pairs, this strategy should not be considered as favorable as a demographic analysis conducted jointly on all populations. We hope that machine learning approaches will contribute to novel tools that would be as equally robust as DILS, but allowing a larger number of populations at once. Such analyses could then allow for better inference of the respective divergence times and modeling of gene flow across populations.

### Conclusion

In conclusion, by integrating demographic inferences with unbiased estimates of nucleotide diversity, we have gained new insights into the evolution of nucleotide diversity in honey bees, including Western honey bees, the world’s most important pollinator. Initially, coalescent-based demographic models provided a clearer understanding of the evolutionary history of Western honey bees in Europe, including the timing of subspecies diversification and gene flow. Our analysis is consistent with widespread postglacial secondary contacts among lineages, which predates modern beekeeping. Through a combination of methods, including detailed LD-based demographic modeling, we also assessed the impacts of modern beekeeping on the genetic diversity of honey bees. Our study provides substantial evidence of a rapid erosion, raising significant concerns when considering the pivotal role of this species for wild plant and crop pollination. Notably, we found evidence suggesting that current conservation programs, especially those on islands, are ineffective in preserving the genetic diversity of black honey bees in Western Europe for the long-term, although potential solutions could be explored. Finally, our findings highlight correlated heterogeneous nucleotide diversity along the genome among the different species, which is consistent with a conservation of recombination landscapes over more than 15 million years of divergence.

## Materials and Methods

### Datasets

Publicly available sequencing data from three datasets, corresponding to samples from four *Apis* species, were reanalyzed. The first dataset is composed of 629 whole-genome sequenced drones from *A. mellifera* and were published by Wragg et al. (2022). In addition, we gathered sequencing data from workers of three other *Apis* species, that were obtained as part of two independent studies, one on *A. cerana* (Chen et al. 2018) and one on *A. dorsata* and *A. laboriosa* (Cao et al. 2023). We randomly selected two individuals per population described in these two studies, after excluding differentiated populations of *A. dorsata* from the Hainan island identified by Cao et al. 2023. In total, we used whole-genome sequence data from 54 individuals, 18 from Cao et al. 2023 and 36 from Chen et al. (2018), see Table S1 for details.

### Reference genome and recombination

Although at least another high-quality reference genome is available for *Apis mellifera* (Eynard et al. 2024), all subsequent analyses are based on the Amel_HAv3.1 reference (Wallberg et al. 2019). This choice is primarily due to the availability of the associated gene catalog (e.g. used in Fig. 5). Regarding recombination, we used a refined version of the recombination map from Wragg et al. (2022), which is available at https://github.com/avignal5/SeqApiPop/blob/master/Scripts_Genetic_map/. This map, derived from the sequence data of Liu et al. (2015), is based on 55 individuals from three colonies. Despite the notably high recombination rates in honey bee (19 to 37 cM/Mb, *i.e.* around 20 times higher than in humans (*e.g.* Beye et al. 2006; Conlon et al. 2023; Kent et al. 2012; Liu et al. 2015; Wallberg et al. 2015)), such a map can only provide a rough view on the levels of local recombination (e.g. internal track in Fig. 5). Given the limited resolution of the available genetic maps, they do not scale with the 100-kbp windows used for the rest of our analyses. To generate recombination-related information at 100-kbp resolution, we used different proxies to identify candidate regions for low and high recombination, namely the repeated and GC content in the HAv3.1 reference genome (Fig. S9). GC was used since it is an excellent proxy of local recombination rate, because GC-biased gene conversion (gBGC) is a recombination-associated process (*e.g.* Duret & Galtier, 2009). But, we also used the proportion of softmasked positions in the reference genome, since repetitive sequences including TE accumulate in low recombination regions of the genome (*e.g.* Dolgin & Charlesworth, 2008). Based on the distributions of GC, repeated and GC in repeated regions (Fig. S9), we discriminate regions based on empirically determined thresholds, consistent with shoulders in the observed distributions (Fig. S9). We consider as low recombination regions those satisfying the three criteria: low GC, low GC at softmasked positions only and high TE content (in red in the external track Fig. 5). Reciprocally, the high recombination regions were defined as those with higher GC contents and lower TE (in green, Fig. 5). Windows satisfying some but not all criteria were considered as intermediate (in yellow, Fig. 5). Note that this strategy was preferred to a more population-oriented strategy, such as the computation of population-scale recombination rates, in order to ensure the independence of the detection with regards to the different population genetic statistics used in the manuscript. For instance, our detection here does not rely at all on *N_e_*.

### Mapping and SNP calling

For the *A. mellifera* samples, our SNP calling was based on the g.vcf generated by Wragg et al. 2022. In brief, Wragg and collaborators previously used BWA mem (Li et al. 2013) to map reads against the Amel_HAv3.1 reference (Wallberg et al., 2019) and then used local realignment and base quality score recalibration (BQSR) were performed using GATK (McKenna et al. 2010), using SNPs called with GATK haplotypecaller as covariates for BQSR, prior to genotype each drone independently and generate the g.vcf files (for details, see Wragg et al. 2022). For the other *Apis* species, we also mapped them on the same *A. mellifera* genome in order to allow a direct comparison and determine ancestral *vs.* derived alleles (*i.e.* same genomic coordinates). We downloaded the raw data (Table S1) and then performed all steps following Wragg et al. 2022 in order to be consistent.

### Population structure

Prior to performing sequence reconstructions, we checked for the absence of within-population structuration, that could correspond to a form of hierarchical structure not previously identified by Wragg et al. 2022. Despite the initial filtering of the *A. mellifera* individuals exhibiting high Q-values assuming the 9 genetic clusters from Wragg et al. (2022) based on their Supplementary Table 1, additional subtle subgroups remain indeed possible. We performed PCA for each cluster as implemented in the snpgdsPCA function in the R package SNPrelate in order to check for within-group panmixia. Assuming panmixia, individuals are expected to be equally distant in the analysis, with a low proportion of variance explained by the first axis (roughly similar to 1 / #individuals). Based on this expectation, we used a simple rule to exclude outliers, by considering acceptable contributions to the first PCs not exceeding by 50% this expectation (*i.e.* an enrichment factor of 1.5). Above these values, individuals were discarded (see Table S2).

### Kinship

To avoid inaccurate demographic inferences or diversity estimates, we aimed to discard pairs of first-degree relatives. As population structure may lead to underestimation of kinship, we analyzed each genetic cluster separately, estimating pairwise kinship only among samples within each group. In addition, we divided the Carnica group into German and non-German samples, as these two subpopulations were found to be clearly differentiated along PC1 (Fig. S1). For each population, kinship was estimated using a reduced set of variants that met the following criteria: (i) single-nucleotide, (ii) biallelic, (iii) autosomal, (iv) QUAL>200, (v) call rate >90%, (vi) minor allele count >= 2, and (vii) LD with other variants < 60%. We then estimated kinship either with hmmIBD (Schaffner et al. 2018) for the haploid individuals (*A. mellifera*), or KING (version 2.2.5; Manichaikul et al. 2010) for the diploid individuals of the three other species (*A. cerana*, *A. dorsata* and *A. laboriosa*). We ran hmmIBD with parameters nchrom set to 16 and rec_rate set to 9.04 x 10^-7^, while KING was run with default parameters. For pairs of samples with kinship above 0.3536, a value that corresponds to the boundary between the first and second degree of kinship (1/2^1.5^), we discarded one individual per pair. The rules used to select the individual for exclusion were as follows: first, we preferentially excluded individuals involved in multiple pairs. In cases where individuals were involved in only a single pair, we chose to exclude the one with the lowest call rate.

### Sequence data reconstruction

To generate unbiased estimates of nucleotide diversity we reconstructed fasta sequences from the called genotypes and non-variant positions using the approach described in Leroy et al. (2021a, 2021b, 2023); see also Korunes & Samuk, 2021 for the importance of considering non-variant positions to generate unbiased estimates of diversity). The approach was adapted to reconstruct haploid sequences for all the *A. mellifera* individuals in our dataset. Briefly, the pipeline reconstructs the sequence by considering the coverage at each genomic position. All positions that are not in between the 5th and 95th centiles of the individual coverage, as well as those with coverage lower than 3 were hard-masked in the reconstructed sequences. For positions satisfying the coverage criteria, the reference allele is used in the reconstructed sequences with the exception of variant positions annotated as ‘PASS’ in the filter field - for which the allele associated with the genotype call is used instead. Only biallelic SNPs are considered; multiallelic variants and INDELs are discarded. As a consequence, nucleotide diversity estimates reported in this work are based on the diversity at biallelic SNPs only.

### Phylogenetic reconstruction

The evolutionary history of the different subspecies and species was inferred by using the phylogenetic reconstruction method implemented in PanTools (v. 4.2.2; Jonkheer et al. 2022), a pangenomic tool which is computationally adapted to the joint analysis of a large number of genomes. For this analysis, we used all individuals, except genetic clusters from *A. mellifera* corresponding to recent breeds, namely the Buckfast and Royal Jelly. From the whole-genome reconstructed sequences, 50 windows of 100-kbp were randomly sampled. PanTools was then used to construct a pangenome with the “*build_pangenome*” function. We let PanTools select the most optimal k-mer size automatically, as recommended. Subsequently, the “*kmer_classification*” function was used on the built K-mer database to generate a K-mer distance matrix. Finally, a neighbor-joining tree estimation was generated under the R package *ape* (Paradis & Schliep, 2019) based on the mash distance calculated by PanTools. In total, the analysis was replicated 200 times, considering 200 distance matrices generated by independent samplings of 50 windows. The 200 generated trees were concatenated into a single median tree with the *compute.brlen* function from *ape*. The circular visualization of the phylogenetic tree (Fig. 1D) was generated with the “*circlize_dendrogram*” function from the R package dendextend (Galili, 2015), while the phylogram (Fig. S2) was generated with the “*plotBS*” function from the R package phangorn (Schliep, 2011) with p=0 to report all bootstrap values.

### ABC-based demographic inferences with DILS

The software DILS (Fraïsse et al. 2021), which stands for “Demographic Inference using Linked Selection”, utilizes Approximate Bayesian Computation (ABC) to infer the best demographic scenarios and their underlying parameters from sequence data. DILS has several advantages as compared to other available methods, including (i) suitability for relatively complex demographic scenarios (no likelihood calculations), *e.g.* including a temporal change in *N_e_* during divergence (Var. *N_e_*, in Fig. 2B), (ii) greater computationally efficiency than traditional ABC methods due to machine learning, and (iii) consideration of linked selection, an evolutionary process known to reduce the accuracy of more naive demographic methods (Ewing & Jensen, 2016; Schrider et al. 2016; Johri et al. 2020).

Similarly to traditional ABC approaches, the rationale is to simulate genetic data under various demographic models and compare these simulations to observed data on the basis of a set of summary statistics. Specifically, DILS hierarchically considers four model choices for the two-population mode: i) current isolation vs. ongoing migration, ii) scenarios of speciation (*i.e.* Strict Isolation vs. Ancient Migration assuming current isolation or Isolation with Migration vs. Secondary contacts assuming ongoing migration), iii) homogeneous or heterogeneous *N_e_*, and (iv) homogeneous or heterogeneous migration rates (*N_e_*.m). We used the default set of parameters, optionally including the 2D-Site Frequency Spectrum (2D-SFS) as summary statistics (e.g. “SFS used”, Fig. 2B). We evaluated the performance of the ABC inferences by (i) visualizing PCA results based on the summary statistics of both simulations and observed data and ii) analyzing goodness-of-fit tests based on the summary statistics to assess how well simulations match the summary statistics of the observed data. Parameters were then estimated based on the optimized posterior of the best-supported model using both a rejection/regression approach (ABC neural network) and a random forest, as implemented in DILS (Fraïsse et al. 2021; see also the DILS online manual).

Since coalescent-based methods are informative about medium to long-term evolutionary history, demographic inferences were performed on all *A. mellifera* subspecies, but were not performed on genetic clusters corresponding to recent breeds such as ‘Buckfast’ and ‘RoyalJelly’. Samples from *A. laboriosa* were used as an outgroup to orient mutations. All information regarding prior parameter values is available in the GitHub repository (https://github.com/ThibaultLeroyFr/SeqApiPop_WGShoneybeeDataReanalysis/ blob/main/DILS_analysis). In brief, the by-default prior values were used for population sizes for each genetic cluster, including an *N_e_* assumed to be between 0 and 500,000 individuals and divergence time (T_SPLIT_) that can be up to 2,000,000 generations ago. Demographic changes in migration rates (*T*_AM_ and *T*_SC_ for the AM and SC scenarios), as well as for a single temporal change in *N_e_* (expansions or bottlenecks), if any (see “Var. *N_e_*“ in Fig. 2B), are sampled from a uniform distribution between 0 and T_SPLIT._ The mutation rate was set to 3.4×10^-9^, as estimated for single-nucleotide mutations in honey bees by Yang et al. (2015). We reported values for a series of key parameters: *N_e_*, T_SPLIT_, T_AM_ or T_SC_ in cases of support for AM or SC scenarios, as well as potential changes in *N_e_* during divergence (“Var. *N_e_*”, Fig. 2B). Given that DILS provides posteriors assuming diploid individuals, we rescaled the posteriors of *N_e_* and inferred times to units of 1.5*N*_e_ individuals and generations to account for haplodiploidy in honeybees (*i.e.* 30 gene copies correspond to 20 haplodiploid individuals rather than 15 diploid individuals). We decided not to report the posterior of the effective migration rates, as this parameter is poorly estimated under DILS (Fraïsse et al. 2021) and, even if it were accurately estimated, would still require additional caution in interpretation (Burban et al. 2024).

### Inference of recent changes in effective population sizes using GONE

In order to investigate the recent changes in *N_e_* in each species or genetic cluster, we used the approach implemented in GONE (Santiago et al. 2020). For each of the genetic clusters analyzed and ran, we randomly sampled 500,000 high-quality biallelic SNPs among the list of SNPs with a call rate exceeding 90%. Given the specificity of the haplodiploidy in the model, we used different strategies for *A. mellifera* vs. the other species (*A. cerana*, *A. dorsata* and *A. laboriosa*) since haploid drones and workers were sequenced, respectively. For inferences based on workers, GONE was run in “unknown phase, diploid” mode. However, for drones, we considered the “diploid with known phase” mode, since GONE does not include a haploid mode. We generated *n* /2 fully independent hypothetical queens with known phases by pairing the *n* haploid drones of each panmictic cluster. For genetic clusters with an odd number of individuals, one individual was randomly excluded when forming pairs. Also, we considered the genetic map from Wragg et al. 2022 available at (https://github.com/avignal5/SeqApiPop/blob/master/Scripts_Genetic_map/ GenetMap_march_2023_AV.txt) and provided these positions to GONE. All other input parameters of GONE were kept as default, which include a number of internal replicates (REPS) of 40, which means that GONE automatically performs 40 prediction replicates for each run. In addition to the internal replicates, we decided to independently perform external replicates of GONE runs on independent samplings of 500k SNPs. This strategy was put in place in order to check the congruence of the results (see Supplementary Note 2).

### Nucleotide diversity

Nucleotide diversity and Tajima’s D were computed in non-overlapping 100-kbp sliding windows for each genetic cluster identified in *A. mellifera* (Wragg et al. 2022) or from the other *Apis* species (*A. cerana*, *A. dorsata*, and *A. laboriosa*), following the methodology described in Leroy et al. (2021a). Our strategy is based on the reconstructed sequences (see above) in order to yield reliable estimates of nucleotide diversity (Korunes & Samuk, 2021). All scripts are available in the GitHub repository. Variance in nucleotide diversity and Tajima’s D distributions per genetic cluster was visualized using the *geom_violin* function from ggplot2 (Wickham et al. 2016), correlation matrix were plotted with the R package corrplot (Wei & Simko, 2021) and circular plots as implemented in the R package circlize (Gu et al. 2014). For circular visualization, the *circos.trackLines* function was used to plot nucleotide diversity variation along the chromosomes, with the baseline set to the median of the per-group distribution of nucleotide diversity. This baseline value was used to highlight regions with extremely low (bottom 10%, red) and high diversity (top 10%, blue) across the genome.

### Evaluation of large scale synteny

Two different approaches were used to evaluate the large-scale synteny among different genomes. First, we used GeneSpace (Lovell et al. 2022), an approach that combines OrthoFinder (Emms & Kelly, 2019) and MCScanX (Wang et al. 2012) to investigate synteny on the basis of the comparisons of orthologous gene orders between genomes. The full GeneSpace pipeline, including the companion R scripts were used for the comparisons between *A. mellifera* (HAv3.1, Wallberg et al. 2019) and *B. terrestris* (v1.2, Crowley et al. 2023). In the case of the comparison between the reference genomes of *A. cerana* (ACSNU2.0, Park et al. 2015) and *A. mellifera* (HAv3.1, Wallberg et al. 2019), we only considered the results of OrthoFinder in order to be able to plot all 1:1 orthologous relationships with the R package ‘Circlize’ along chromosomes, along the variation of the GC content (Fig. S8). Second, in the case of *A. mellifera* and *A. cerana*, we also performed genome-wide dot plots using D-genies (Cabanettes & Klopp, 2018) to identify conserved and disrupted syntenic regions between the two reference genomes that are whole-genome aligned with minimap2 (Li, 2018).

## Supporting information

SupFile1

SupFile2

## Acknowledgments

We are grateful to the genotoul bioinformatics platform Toulouse Occitanie (Bioinfo Genotoul, https://doi.org/10.15454/1.5572369328961167E12) for providing support, especially regarding computing and storage resources. The *Apis mellifera* data referenced in our previous publication (Wragg et al. 2022) was generated at the GeT platform in Toulouse, France, and was funded by the Animal Genetics division of INRAE and the SeqApiPop programme (FranceAgriMer grant 14-21-AT). We deeply appreciate the continued collaboration and discussions with the co-authors of the earlier publication. TL also thanks Camille Roux and Christelle Fraïsse for their insightful feedback on demographic inferences under DILS, as well as broader discussions on future developments as part of our ANR RadiaSpe project (ANR-23-CE02-0032). TL also acknowledges Mathieu Gautier and Arnaud Estoup for their valuable comments, especially regarding the inference of recent changes in effective population sizes. Additionally, TL is grateful to Matthew Webster and Andreas Wallberg for their discussions on the genetic diversity and adaptation of honeybees as part of a one-month research stay in Uppsala, Sweden, funded by the SFVE-A program of the French Institute in Sweden and the International Relations Department (DRI) of INRAE.

## Data and script availability

All sequencing data utilized in this study are publicly accessible, with corresponding SRA accession numbers provided in Table S1 and File S1. Additionally, all scripts used in the analysis are available on a dedicated github repository: https://github.com/ThibaultLeroyFr/SeqApiPop_WGShoneybeeDataReanalysis.

