## Supporting Information for "Inferring long-term and short-term determinants of genetic diversity in honey bees: Beekeeping impact and conservation strategies"


| | |
|---|---|
| **Supporting Note S1** (discussion regarding inbreeding depression) | **p. 2** |
| **Supporting Note S2** (among-replicate congruence of GONE results) | **p. 4** |
| **Figure S1** (PCAs) | **p. 6** |
| **Figure S2** (Phylogenetic tree) | **p. 8** |
| **Figure S3** (Distributions of diversity depending on recombination regions) | **p.9** |
| **Figure S4** (Correlations of diversity landscapes depending on recomb. regions) | **p. 10** |
| **Figure S5** (Distributions of Tajima's D depending on recombination regions) | **p.11** |
| **Figure S6** (Circular visualization of correlated landscapes of recomb. and Taj. D) | **p.12** |
| **Figure S7** (Correlations of Tajima's D landscapes depending on recomb. regions) | **p.13** |
| **Figure S8 (**Synteny) | **p.14** |
| **Figure S9 (**Distributions of GC and TE contents in the reference genome) | **p. 15** |
| **Table S1** (complete list of individuals included in the study) | **p. 16** |
| **Table S2** (samples excluded - kinship & within-cluster structure) | **p. 34** |
| **Table S3** (restricted list of 267 *A. mellifera* samples) | **p. 35** |

**Supplementary Note S1 : discussion regarding inbreeding depression**

Although our initial objective was to exclude close family relationships in our dataset (*i.e.* outliers in Fig. 1B), the analysis of the whole distribution of $P_{IBD}$ provided important insights regarding the levels of inbreeding in each genetic group (Fig. 1B, Sup. Note 1 Table S1). High kinship values were reported for the two atlantic island conservatories, with respectively the highest and third highest median values for Colonsay, Scotland (median=0.110, interquartile range (IQR): 0.094-0.139) and Ouessant, France (0.083, IQR: 0.071-0.098), respectively. Similarly high median values were reported for breeds ('RoyalJelly': ranked #2, median=0.103, IQR: 0.081-0.141; 'Buckfast': ranked #5, median=0.062, IQR: 0.045-0.082). German Carnica, which could be similarly considered as recent breeds given their intense management (Hoppe *et al.* 2020) despite being part of the C-lineage, also exhibit very high $P_{IBD}$ values (ranked #4, median=0.067, IQR:0.049-0.145). All together these results provide two major insights.

First, conservatories on small isolated islands have already induced relatively high inbreeding values even after a limited number of generations (Fig. 3), as a result of a reduced island surface available to host hives. Indeed, the reported levels of kinship strongly depart from the one observed from continental Spain samples (Iberiensis, median: 0.002; IQR: 0-0.006), the latter exhibiting the lowest kinship among all populations investigated. Interestingly, the Mellifera population, which corresponds to an intermediate group, composed both of samples from the Porquerolles island and the nearby continental conservatory of Solliès, exhibits an intermediate situation (median: 0.039, IQR: 0.014-0.079), suggesting that the inbreeding depression on small islands can be controlled by regularly importing queens from continents.

Second, intensification of beekeeping, including the rearing of new queens, the control of their fertilization, as well the breeding programs, have contributed to a considerable increase in the inbreeding of the population. This is the case especially for 'RoyalJelly' and German Carnica, for which samples are associated with breeding programs that are at least partly based on artificial insemination. For 'RoyalJelly', the high observed values (median=0.103, IQR: 0.081-0.141) contrast with the low values observed with Ligustica (median=0.008, IQR=0.006-0.11), for which the genetics is assumed to originate (see also Fig. 3). Similarly, German Carnica (median=0.067, IQR:0.049-0.145) exhibit far higher values than non-German values (median=0.025, IQR:0.019-0.033). The discrepancy is however lower because non-German Carnica are also impacted by breeding, albeit less intensively.

*Sup. Note 1 Table S1: Summary statistics of the distributions of $P_{IBD}$ for the genetic clusters. Values for quantiles 0.05, 0.25 (1st quartile), 0.5 (median), 0.75 (3rd quartile) and 0.95.*

| Population | Lineage | Quant (0.05) | Quant (0.25) | median | Quant (0.75) | Quant (0.95) |
|---|---|---|---|---|---|---|
| Mellifera | M | 0.0033 | 0.014 | 0.039 | 0.0789 | 0.177 |
| Iberiensis | M | 0 | 0.000412 | 0.00172 | 0.00416 | 0.00717 |
| Colonsay | M | 0.0742 | 0.0942 | 0.11 | 0.139 | 0.209 |
| Ouessant | M | 0.0565 | 0.0709 | 0.0831 | 0.0978 | 0.131 |
| Caucasia | O | 0.00114 | 0.00206 | 0.00302 | 0.00608 | 0.0209 |
| Carnica | C | 0.0145 | 0.0196 | 0.0256 | 0.0346 | 0.0654 |
| *Carnica (German)* | *C* | *0.0309* | *0.0491* | *0.0669* | *0.145* | *0.22* |
| *Carnica (non-German)* | *C* | *0.0144* | *0.0194* | *0.0249* | *0.0333* | *0.0579* |
| Ligustica | C | 0.00348 | 0.00559 | 0.00766 | 0.0115 | 0.0184 |
| RoyalJelly | Breeds | 0.0592 | 0.0815 | 0.103 | 0.141 | 0.268 |
| Buckfast | Breeds | 0.0276 | 0.0447 | 0.0616 | 0.0825 | 0.245 |

### Supplementary Note S2 : among-replicate congruence of GONE results

To investigate the recent changes in $N_e$ in each species or genetic cluster, we used the approach implemented in GONE (Santiago *et al.* 2020). By default, GONE automatically performs 40 internal prediction replicates for each run (REPS parameter). However, we have decided to also perform external replicates by completing 40 different sampling of 500k SNPs and GONE runs for each genetic group. This strategy was put in place in order to check the congruence of the results. In practice, we especially assessed whether the mean of all geometric mean or median provide more accurate inferences at a given timestep, we compared both metrics to those obtained in (i) a replication of the analysis over the set of selected variants and in (ii) a replication of the analysis involving the random drawing of another set of variants. Considering correlations between the various estimates, we computed the median and the mean of the 40 replicate geographic means at each generation. Comparing replicates of analysis, we found that the median among the 40 independent inferences of $N_e$ to provide slightly more consistent results than the average (e.g. Pearson's correlation coefficients of 0.945 and 0.933, respectively in Sup. Note 2 Fig. 1). For this reason, we consider the median as the most appropriate aggregation method.

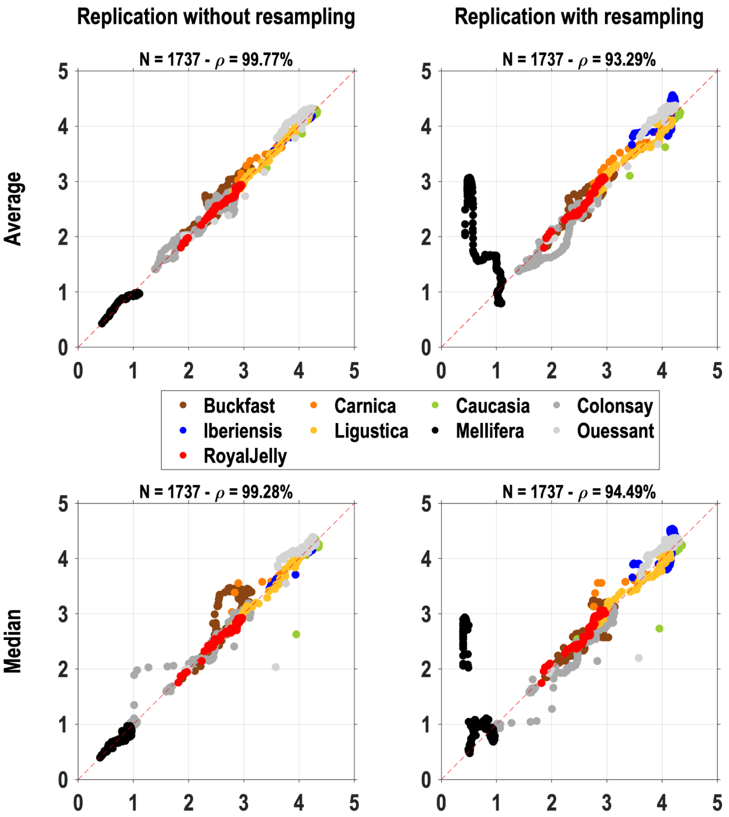


*Sup. Note 2 Figure 1. Consistency of GONE-inferred $N_e$ across different runs and replicates. The $N_e$ estimates from an original run (x-axis in any sub-panel) were compared to the $N_e$ estimates of a replicate run (y-axis in any sub-panel), either based on the same sampling of 500k SNP (left column) or based on different samplings (external replication, right column). Estimates were aggregated over 40 replicates either by their mean (top row) or their median (bottom row).*

Before considering the median of the 40 *GONE-inferred* $N_e$ as the final summary statistic (Fig. 3), we also visually inspected the among-replicate variances (Sup. Note 2 Fig.

2). With the notable exception of Mellifera (top left in Sup. Note 2 Fig. 2), we reported extremely reduced variability among the replicates based on the observed (IQR).

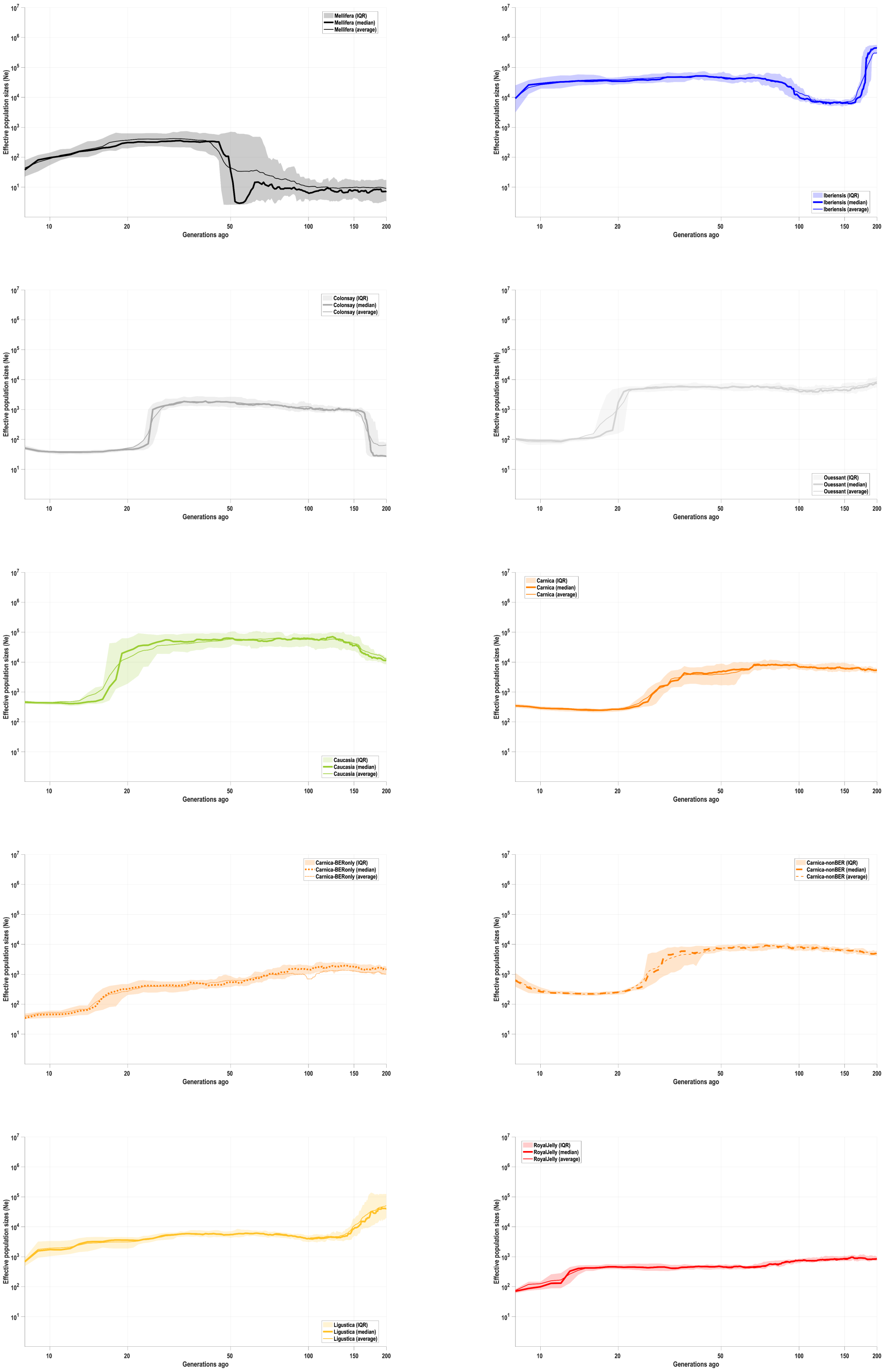

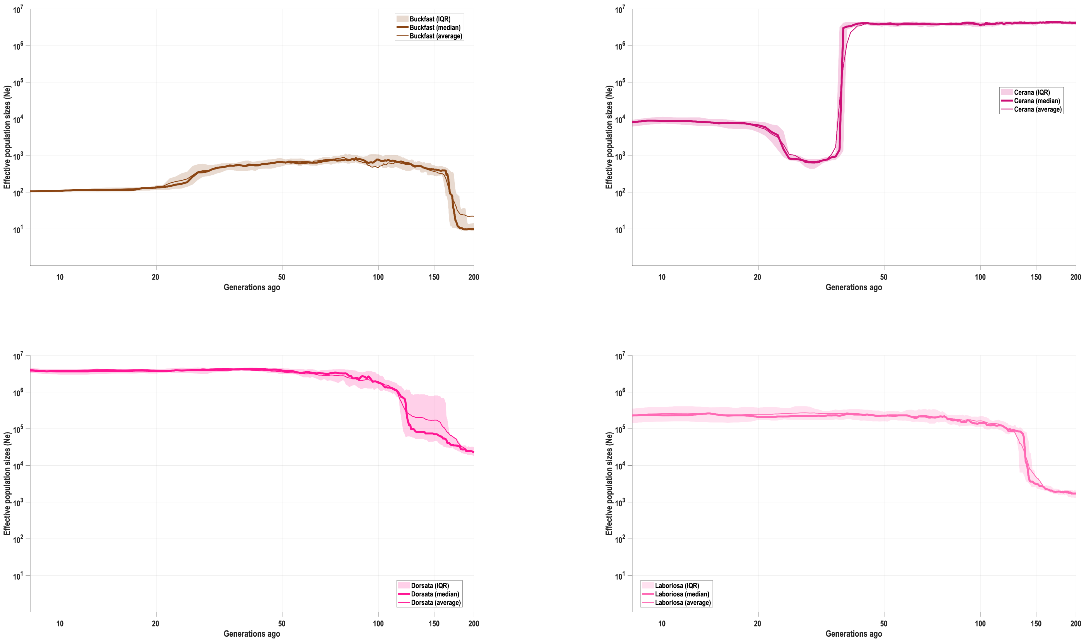


*Sup Note 2 Figure 2: Cluster-per-cluster observed among-replicate variability. From top to bottom: Mellifera (black), Iberiensis (blue), Colonsay (dark grey) Ouessant (light grey), Caucasia (green), Carnica (orange), German Carnica (=BER, dotted orange line), non-German Carnica (=non-BER, dashed orange line), Ligustica (yellow), RoyalJelly (red), Buckfast (brown), A. cerana (dark pink), A. dorsata (pink), and A. ligustica (light pink).*

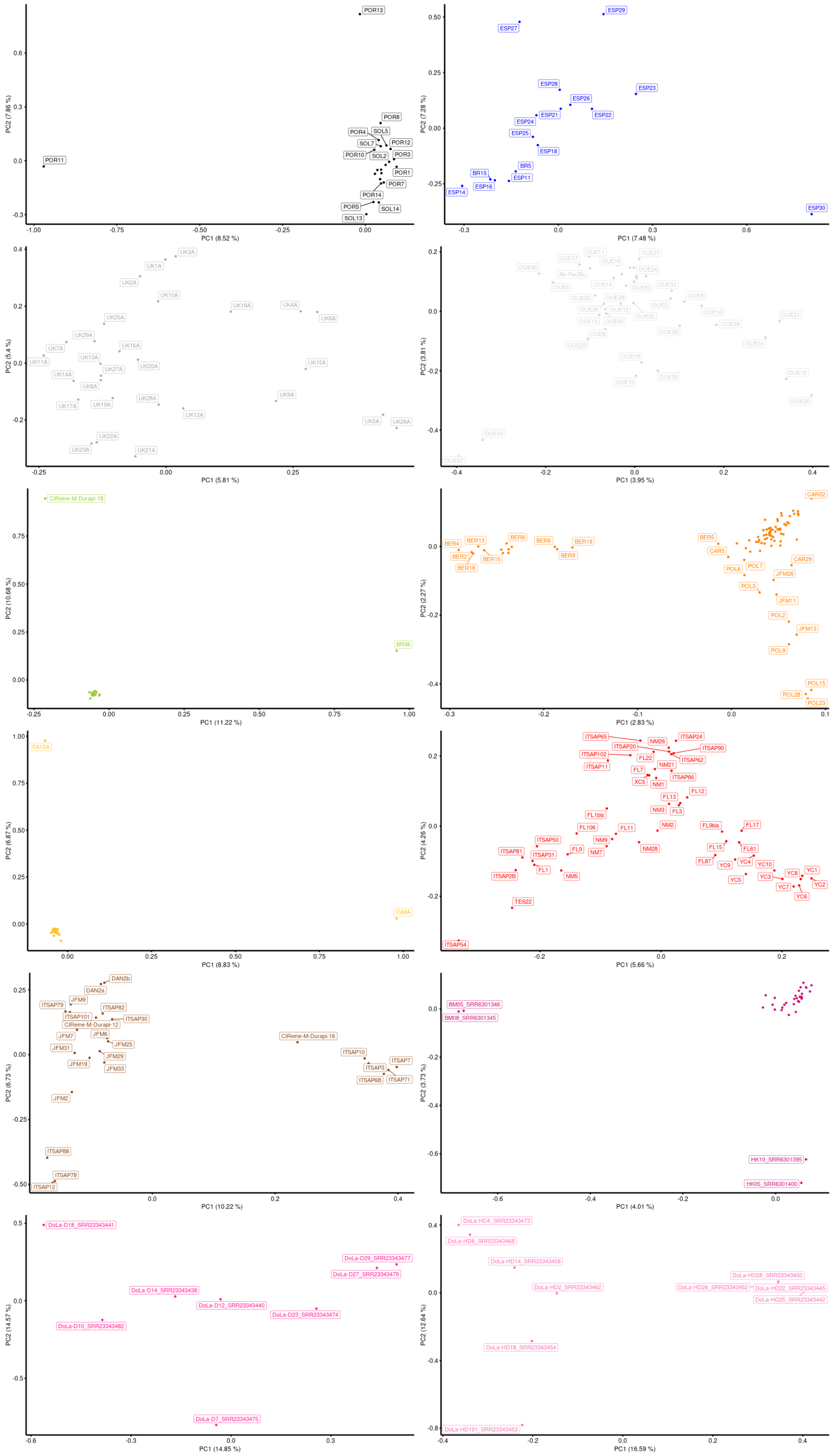


*Figure S1: Axis 1 and 2 of the within-group PCAs for Mellifera (black), Iberiensis (blue), Colonsay (dark grey), Ouessant (light grey), Caucasia (green), Carnica (orange), Ligustica (yellow), RoyalJelly (red), Buckfast (brown), A. cerana (dark pink), A. dorsata (pink), and A. ligustica (light pink).*

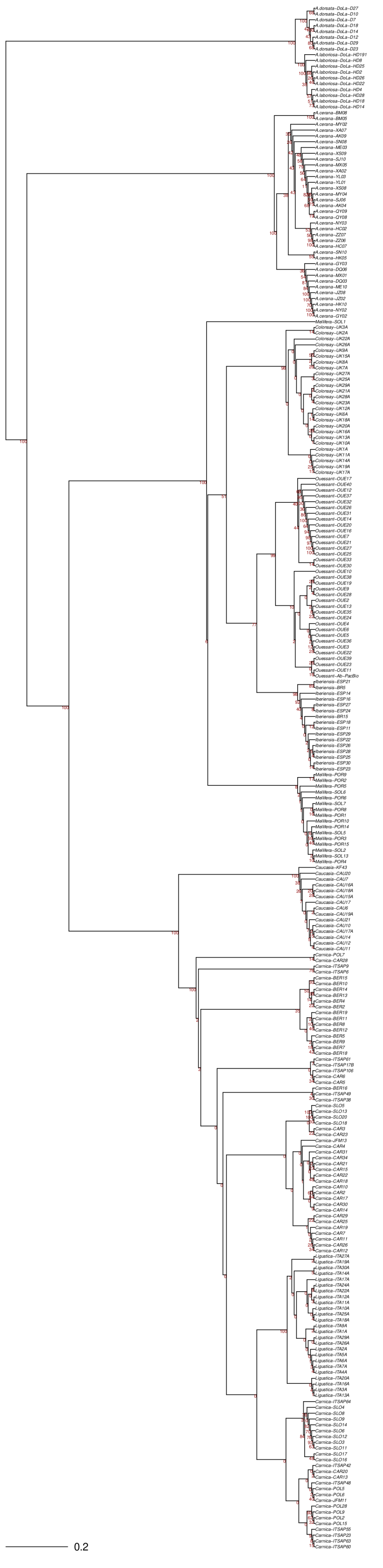

*Figure S2: Detailed phylogenetic tree of Fig. 1D showing all bootstrap values.*

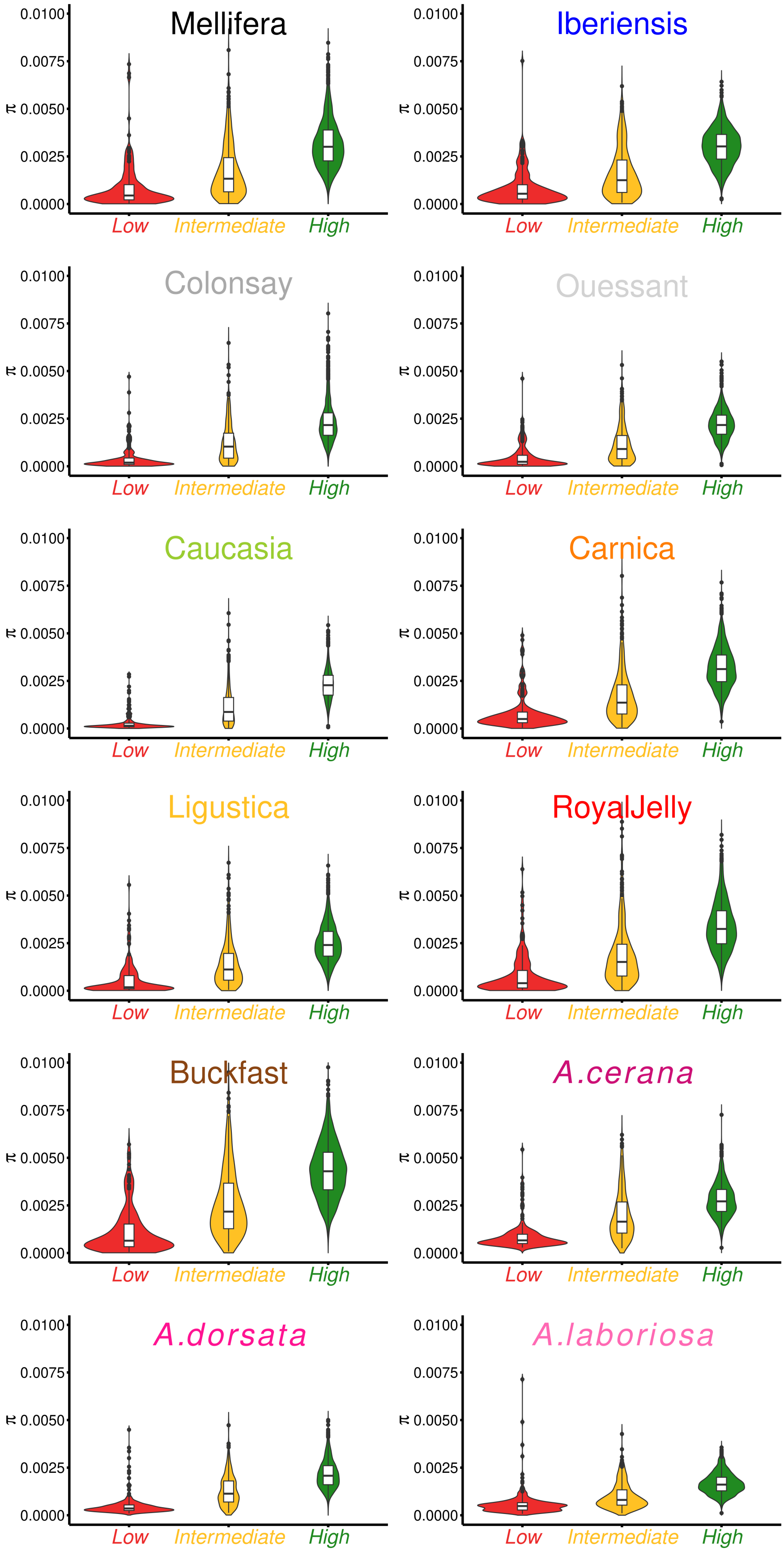


*Figure S3: Distributions of the observed levels of nucleotide diversity in regions consistent with low (red), intermediate (yellow) or high (green) recombination rates. For details regarding the detection of these regions, see Materials and Methods.*

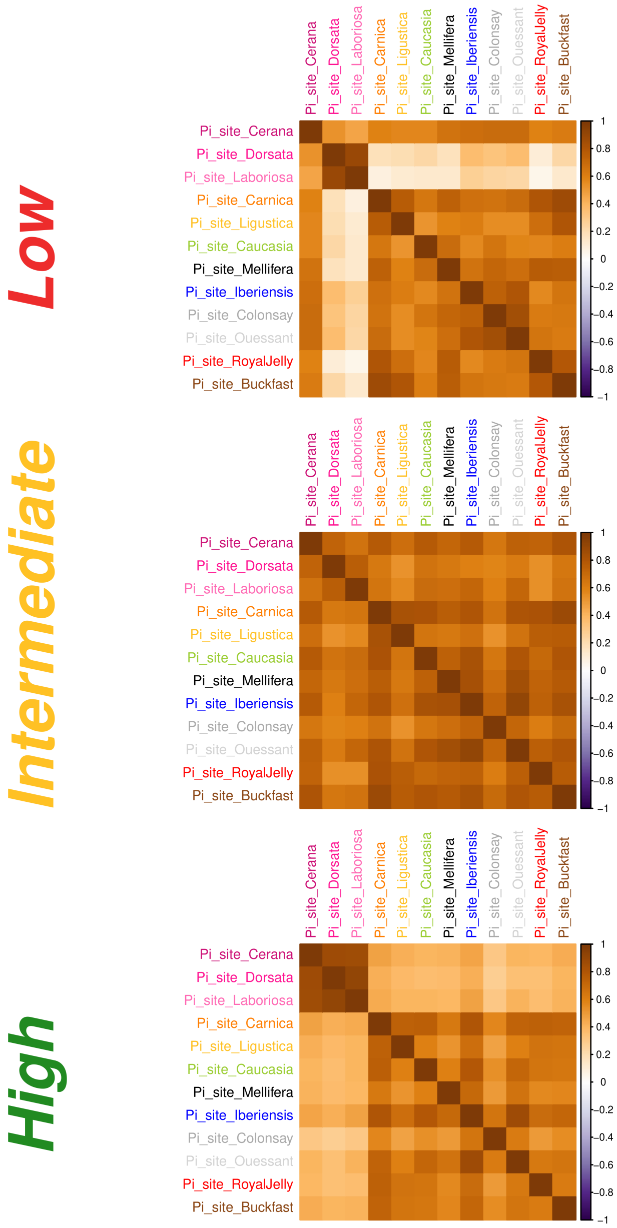


*Figure S4: Correlation plots of the landscapes of nucleotide diversity in regions consistent with low (top), intermediate (middle) or high (bottom) recombination rates. For details regarding the detection of these regions, see Materials and Methods.*

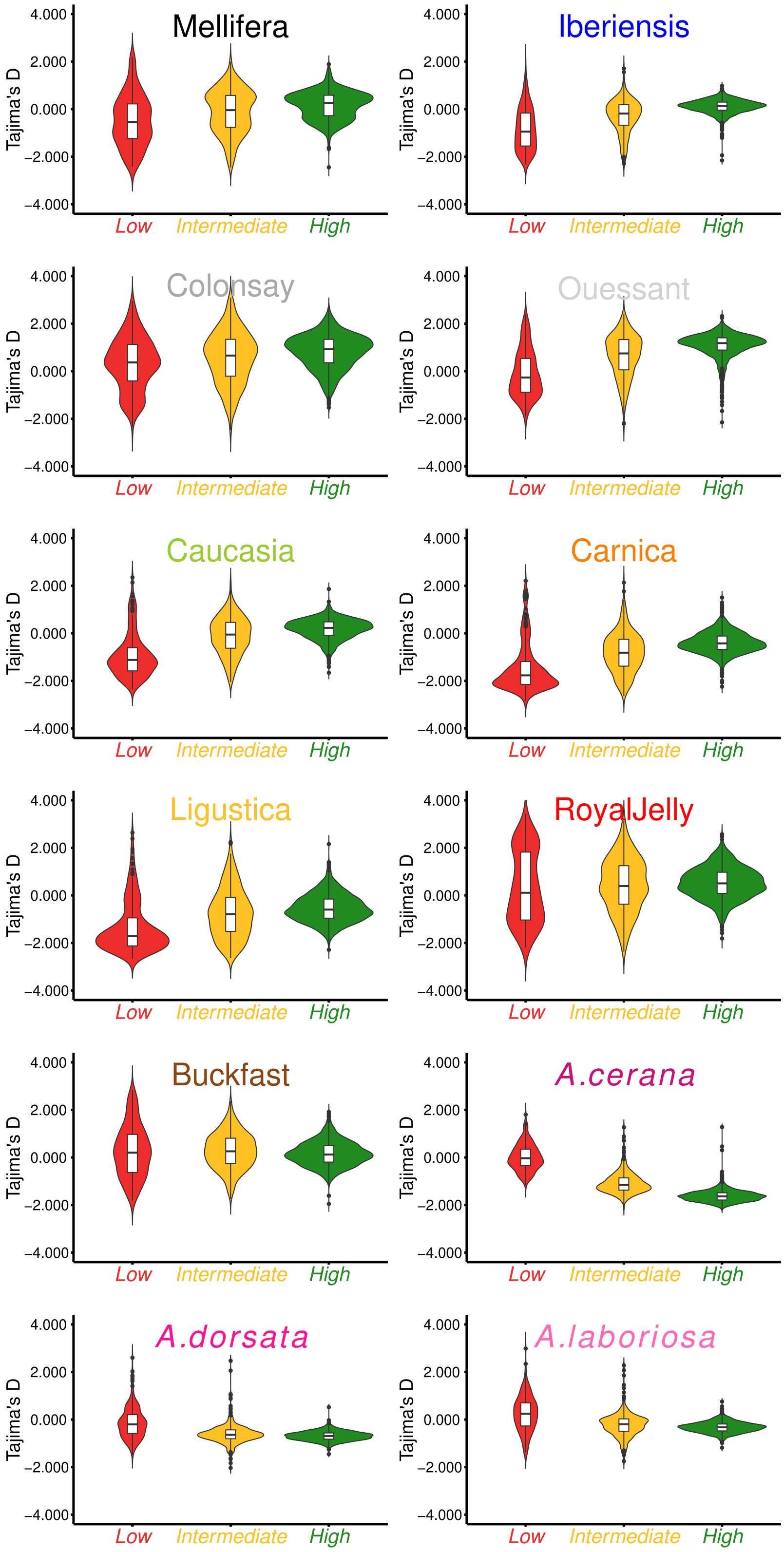


*Fig. S5: Distributions of the Tajima's D in regions consistent with low (red), intermediate (yellow) or high (green) recombination rates. For details regarding the detection of these regions, see Materials and Methods.*

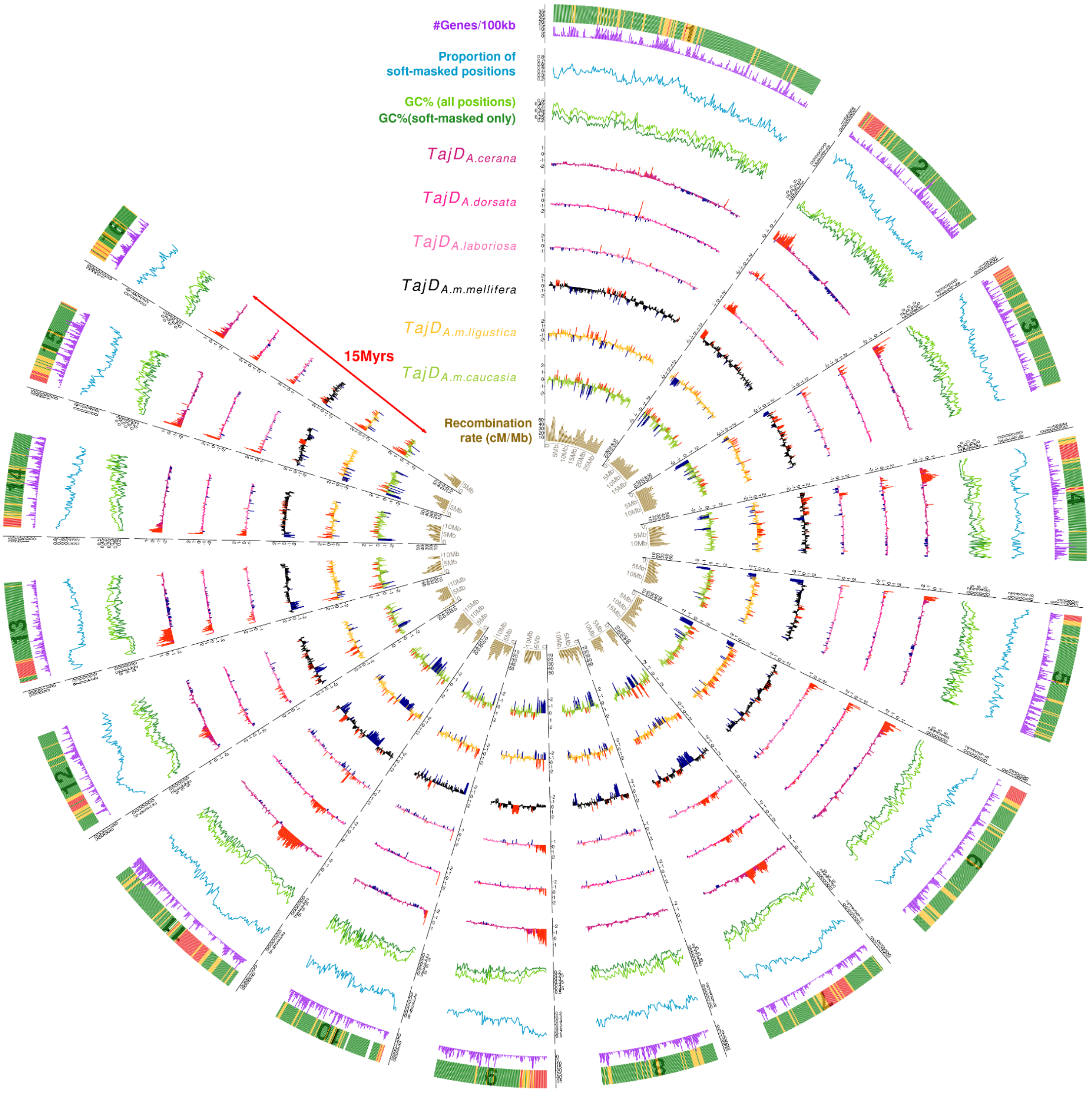


*Figure S6: Circular visualization of the correlated genomic landscapes of recombination and Tajima's D in* Apis, covering more than 15 million years of divergence*. From external to internal: gene density, proportion of soft-masked position on the reference genome, GC-content and GC-content (dark green) at soft-masked (green) positions only, Tajima's D for A. cerana, A. dorsata, A. laboriosa, Mellifera, Ligustica and Caucasia and recombination rate (cM/Mb). The most external track highlights the genomic windows that are consistent with high-recombination (green), intermediate (orange) or low-recombination (red). The chromosome number is also indicated, from 1 to 16.*

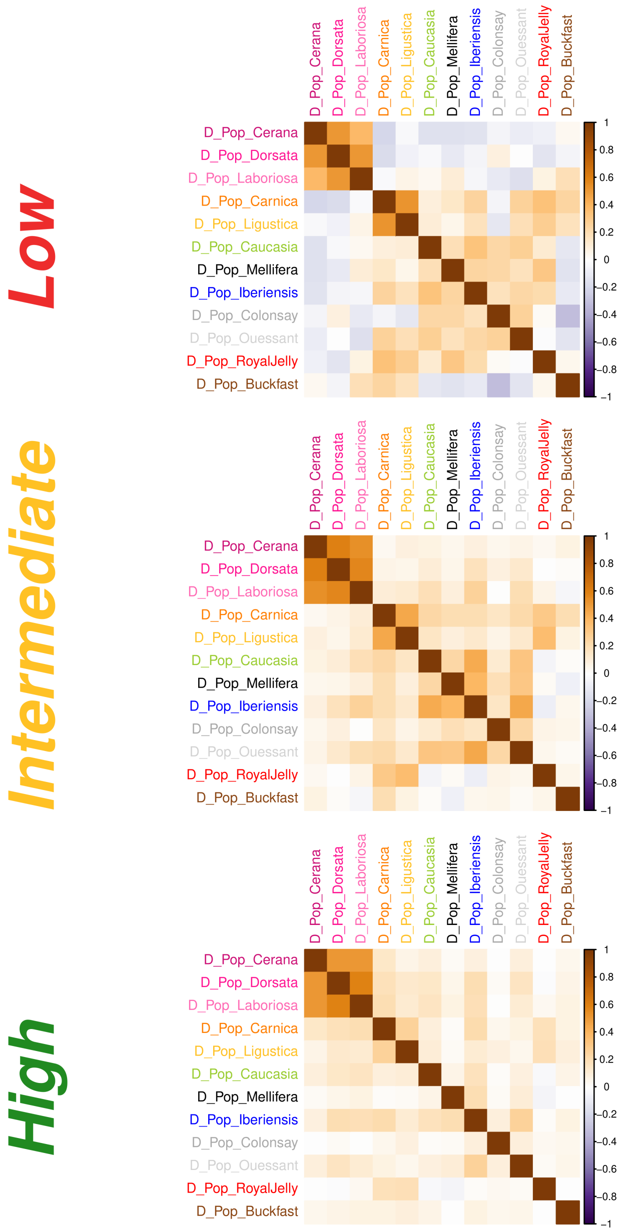


*Figure S7: Correlation plots of the Tajima's D landscapes in regions consistent with low (top), intermediate (middle) or high (bottom) recombination rates. For details regarding the detection of these regions, see Materials and Methods.*

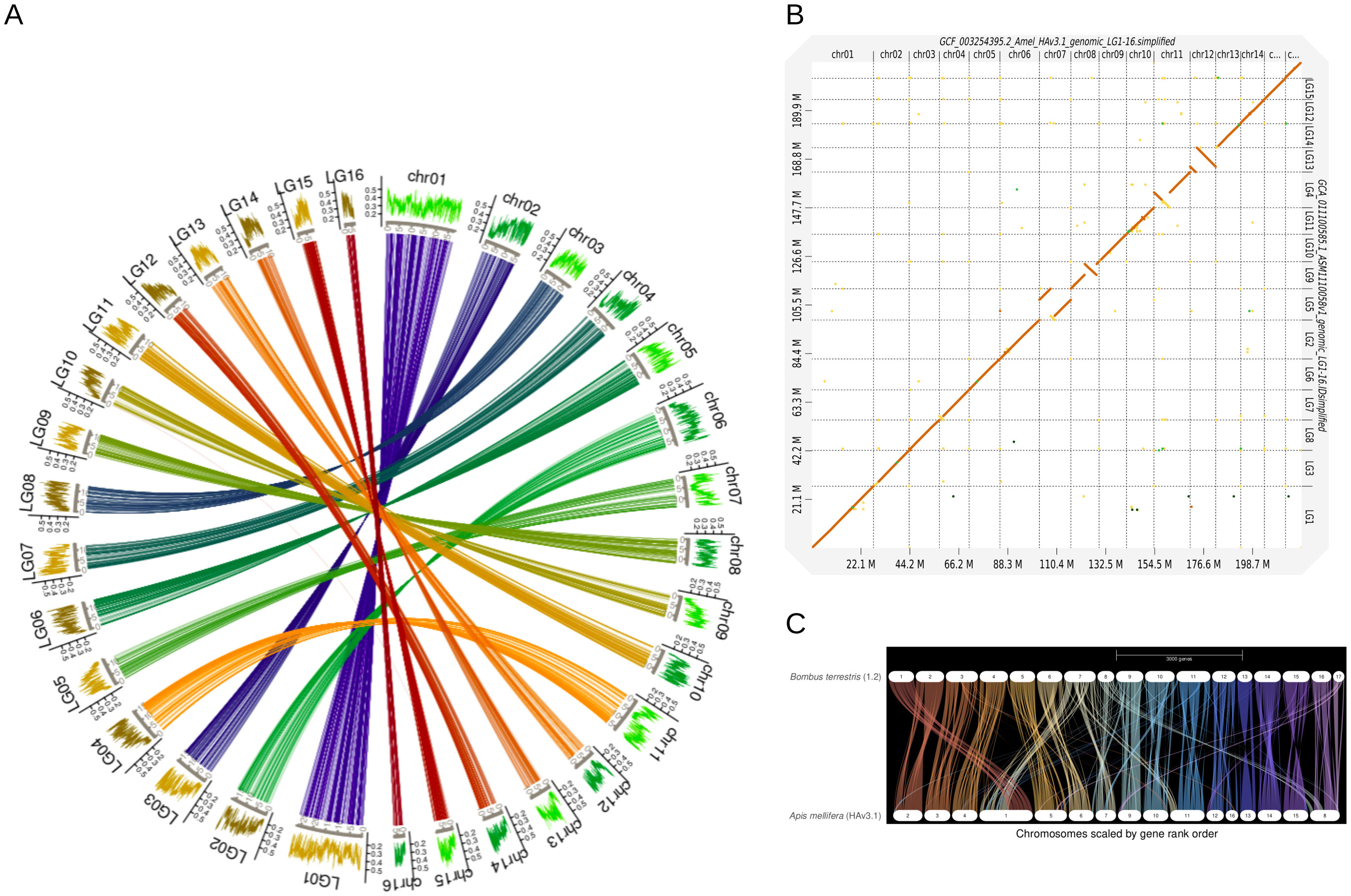


*Figure S8: High conservation of the synteny between the* Apis mellifera *reference genome (HAv3.1) and those of* Apis cerana *(3.0, A & B) and* Bombus terrestris *(1.2, C). Circlize plot showing the GC content (external) and 5.5k 1:1 orthologs inferred by Orthofinder v.2.5.4. B. Dotplot between* A. mellifera *and* A. cerana *as inferred with Dgenies using minimap2. C. Riparian plot drawn from Genespace using the 1:1 orthologs between* A. mellifera *and* B. terrestris *as inferred by Orthofinder v.2.5.4.*

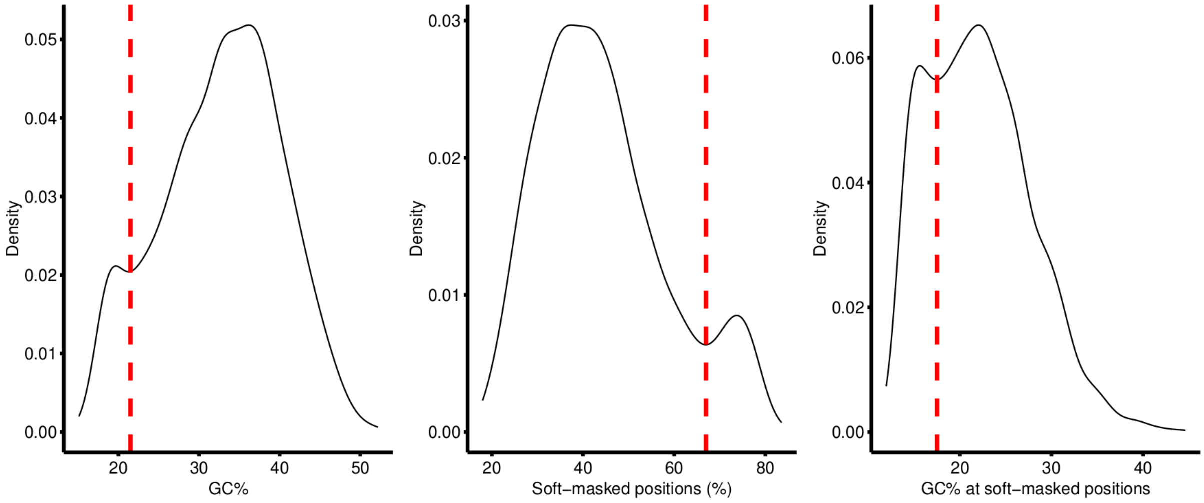


*Figure S9: Distributions of the contents in GC (left), soft-masked positions and GC at soft-masked positions in the HAv3.1 reference genome of* Apis mellifera *(Wallberg et al. 2019). Soft-masked positions are used as a proxy of the repeated content, including transposable elements. The dotted line corresponds to the thresholds used to identify potential low and high recombining regions (see Materials and Methods).*

*Table S1: List of all the samples included in the study. SRA and BioSamples accession numbers to download the data are provided. For more details regarding, including Q-values for* A. mellifera *samples used to filter individuals, see Sup File 1.*

| **Original_s** | **AccessionRunSRA** | **AccessionBioSample** | **Species** | **Gen.Origin** | **ploidy** | **sampleID** | **Label (initial publi.)** | **Geographic.Origin** | **Seq.Machine** |
|---|---|---|---|---|---|---|---|---|---|
| Wragg_et_al_2022_MER | SRR15173334 | SAMN20257031 | *A. mellifera* | Unknown | 1 | BR25 | Ariege Breeder | Ariege, France | HiSeq |
| Wragg_et_al_2022_MER | SRR15173323 | SAMN20257032 | *A. mellifera* | Unknown | 1 | BR26 | Ariege Breeder | Ariege, France | HiSeq |
| Wragg_et_al_2022_MER | SRR15173864 | SAMN20257033 | *A. mellifera* | Unknown | 1 | BR27 | Ariege Breeder | Ariege, France | HiSeq |
| Wragg_et_al_2022_MER | SRR15173850 | SAMN20257034 | *A. mellifera* | Unknown | 1 | BR28 | Ariege Breeder | Ariege, France | HiSeq |
| Wragg_et_al_2022_MER | SRR15173839 | SAMN20257035 | *A. mellifera* | Unknown | 1 | BR29 | Ariege Breeder | Ariege, France | HiSeq |
| Wragg_et_al_2022_MER | SRR15173817 | SAMN20257037 | *A. mellifera* | Unknown | 1 | BR30 | Ariege Breeder | Ariege, France | HiSeq |
| Wragg_et_al_2022_MER | SRR15173813 | SAMN20257038 | *A. mellifera* | Unknown | 1 | BR31 | Ariege Breeder | Ariege, France | HiSeq |
| Wragg_et_al_2022_MER | SRR15173812 | SAMN20257039 | *A. mellifera* | Unknown | 1 | BR32 | Ariege Breeder | Ariege, France | HiSeq |
| Wragg_et_al_2022_MER | SRR15173811 | SAMN20257040 | *A. mellifera* | Unknown | 1 | BR33 | Ariege Breeder | Ariege, France | HiSeq |
| Wragg_et_al_2022_MER | SRR15173810 | SAMN20257041 | *A. mellifera* | Unknown | 1 | BR34 | Ariege Breeder | Ariege, France | HiSeq |
| Wragg_et_al_2022_MER | SRR15173905 | SAMN20257042 | *A. mellifera* | Unknown | 1 | BR35 | Ariege Breeder | Ariege, France | HiSeq |
| Wragg_et_al_2022_MER | SRR15173903 | SAMN20257043 | *A. mellifera* | Unknown | 1 | BR36 | Ariege Breeder | Ariege, France | HiSeq |
| Wragg_et_al_2022_MER | SRR15173511 | SAMN20257015 | *A. mellifera* | *A.m.mellifera* | 1 | BR10 | Ariege Conservatory | Ariege, France | HiSeq |
| Wragg_et_al_2022_MER | SRR15173467 | SAMN20257019 | *A. mellifera* | *A.m.mellifera* | 1 | BR15 | Ariege Conservatory | Ariege, France | HiSeq |
| Wragg_et_al_2022_MER | SRR15173378 | SAMN20257027 | *A. mellifera* | *A.m.mellifera* | 1 | BR21 | Ariege Conservatory | Ariege, France | HiSeq |
| Wragg_et_al_2022_MER | SRR15173902 | SAMN20257044 | *A. mellifera* | *A.m.mellifera* | 1 | BR38 | Ariege Conservatory | Ariege, France | HiSeq |
| Wragg_et_al_2022_MER | SRR15266709 | SAMN20432941 | *A. mellifera* | *A.m.mellifera* | 1 | BR45-PE | Ariege Conservatory | Ariege, France | HiSeq |
| Wragg_et_al_2022_MER | SRR15173892 | SAMN20257053 | *A. mellifera* | *A.m.mellifera* | 1 | BR49 | Ariege Conservatory | Ariege, France | HiSeq |
| Wragg_et_al_2022_MER | SRR15173891 | SAMN20257054 | *A. mellifera* | *A.m.mellifera* | 1 | BR5 | Ariege Conservatory | Ariege, France | HiSeq |
| Wragg_et_al_2022_MER | SRR15173887 | SAMN20257058 | *A. mellifera* | *A.m.mellifera* | 1 | BR54 | Ariege Conservatory | Ariege, France | HiSeq |
| Wragg_et_al_2022_MER | SRR15173881 | SAMN20257063 | *A. mellifera* | Unknown | 1 | BRT1 | Brittany Breeder | Brittany, France | NovaSeq |
| Wragg_et_al_2022_MER | SRR15173880 | SAMN20257064 | *A. mellifera* | Unknown | 1 | BRT2 | Brittany Breeder | Brittany, France | NovaSeq |
| Wragg_et_al_2022_MER | SRR15173878 | SAMN20257065 | *A. mellifera* | Unknown | 1 | BRT3 | Brittany Breeder | Brittany, France | NovaSeq |
| Wragg_et_al_2022_MER | SRR15173624 | SAMN20257241 | *A. mellifera* | *A.m.mellifera* | 1 | ITSAP85 | Brittany Conservatory | Brittany, France | HiSeq |
| Wragg_et_al_2022_MER | SRR15173616 | SAMN20257248 | *A. mellifera* | *A.m.mellifera* | 1 | ITSAP91 | Brittany Conservatory | Brittany, France | HiSeq |
| Wragg_et_al_2022_MER | SRR15173613 | SAMN20257251 | *A. mellifera* | *A.m.mellifera* | 1 | ITSAP94 | Brittany Conservatory | Brittany, France | HiSeq |
| Wragg_et_al_2022_MER | SRR15173685 | SAMN20257185 | *A. mellifera* | *A.m.mellifera* | 1 | ITSAP21 | Brittany Conservatory | Brittany, France | HiSeq |
| Wragg_et_al_2022_MER | SRR15173696 | SAMN20257175 | *A. mellifera* | Buckfast | 1 | ITSAP13B | Buckfast France | Haut-Rhin, France | HiSeq |
| Wragg_et_al_2022_MER | SRR15173682 | SAMN20257188 | *A. mellifera* | Buckfast | 1 | ITSAP25 | Buckfast France | Haut-Rhin, France | HiSeq |
| Wragg_et_al_2022_MER | SRR15173681 | SAMN20257189 | *A. mellifera* | Buckfast | 1 | ITSAP28 | Buckfast France | Haut-Rhin, France | HiSeq |
| Wragg_et_al_2022_MER | SRR15173676 | SAMN20257193 | *A. mellifera* | Buckfast | 1 | ITSAP30 | Buckfast France | Haut-Rhin, France | HiSeq |
| Wragg_et_al_2022_MER | SRR15173659 | SAMN20257209 | *A. mellifera* | Buckfast | 1 | ITSAP5 | Buckfast France | Haut-Rhin, France | HiSeq |
| Wragg_et_al_2022_MER | SRR15173627 | SAMN20257238 | *A. mellifera* | Buckfast | 1 | ITSAP82 | Buckfast France | Haut-Rhin, France | HiSeq |
| Wragg_et_al_2022_MER | ERR1700887 | SAMEA4521109 | *A. mellifera* | Buckfast | 1 | Buck1 | Buckfast Switzerland | Switzerland | HiSeq |
| Wragg_et_al_2022_MER | ERR1700886 | SAMEA4521110 | *A. mellifera* | Buckfast | 1 | Buck10 | Buckfast Switzerland | Switzerland | HiSeq |
| Wragg_et_al_2022_MER | ERR1700885 | SAMEA4521111 | *A. mellifera* | Buckfast | 1 | Buck11 | Buckfast Switzerland | Switzerland | HiSeq |
| Wragg_et_al_2022_MER | ERR1700884 | SAMEA4521112 | *A. mellifera* | Buckfast | 1 | Buck2 | Buckfast Switzerland | Switzerland | HiSeq |
| Wragg_et_al_2022_MER | ERR1700883 | SAMEA4521113 | *A. mellifera* | Buckfast | 1 | Buck3 | Buckfast Switzerland | Switzerland | HiSeq |
| Wragg_et_al_2022_MER | ERR1700882 | SAMEA4521114 | *A. mellifera* | Buckfast | 1 | Buck4 | Buckfast Switzerland | Switzerland | HiSeq |

| | | | | | | | | | |
|---|---|---|---|---|---|---|---|---|---|
| Wragg_et_al_2022_MER | ERR1700881 | SAMEA4521115 | *A. mellifera* | Buckfast | 1 | Buck5 | Buckfast Switzerland | Switzerland | HiSeq |
| Wragg_et_al_2022_MER | ERR1700880 | SAMEA4521116 | *A. mellifera* | Buckfast | 1 | Buck6 | Buckfast Switzerland | Switzerland | HiSeq |
| Wragg_et_al_2022_MER | ERR1700879 | SAMEA4521117 | *A. mellifera* | Buckfast | 1 | Buck7 | Buckfast Switzerland | Switzerland | HiSeq |
| Wragg_et_al_2022_MER | ERR1700878 | SAMEA4521118 | *A. mellifera* | Buckfast | 1 | Buck8 | Buckfast Switzerland | Switzerland | HiSeq |
| Wragg_et_al_2022_MER | ERR1700877 | SAMEA4521119 | *A. mellifera* | Buckfast | 1 | Buck9 | Buckfast Switzerland | Switzerland | HiSeq |
| Wragg_et_al_2022_MER | ERR1700773 | SAMEA4521223 | *A. mellifera* | Buckfast | 1 | Ticino1 | Buckfast Switzerland | Switzerland | HiSeq |
| Wragg_et_al_2022_MER | ERR1700772 | SAMEA4521224 | *A. mellifera* | Buckfast | 1 | Ticino2 | Buckfast Switzerland | Switzerland | HiSeq |
| Wragg_et_al_2022_MER | ERR1700771 | SAMEA4521225 | *A. mellifera* | Buckfast | 1 | Ticino3 | Buckfast Switzerland | Switzerland | HiSeq |
| Wragg_et_al_2022_MER | ERR1700770 | SAMEA4521226 | *A. mellifera* | Buckfast | 1 | Ticino4 | Buckfast Switzerland | Switzerland | HiSeq |
| Wragg_et_al_2022_MER | ERR1700769 | SAMEA4521227 | *A. mellifera* | Buckfast | 1 | Ticino6 | Buckfast Switzerland | Switzerland | HiSeq |
| Wragg_et_al_2022_MER | ERR1700768 | SAMEA4521228 | *A. mellifera* | Buckfast | 1 | Ticino8 | Buckfast Switzerland | Switzerland | HiSeq |
| Wragg_et_al_2022_MER | SRR15173693 | SAMN20257178 | *A. mellifera* | *A.m. carnica* | 1 | ITSAP16 | Carnica France | France | HiSeq |
| Wragg_et_al_2022_MER | SRR15173691 | SAMN20257180 | *A. mellifera* | *A.m. carnica* | 1 | ITSAP17B | Carnica France | France | HiSeq |
| Wragg_et_al_2022_MER | SRR15173684 | SAMN20257186 | *A. mellifera* | *A.m. carnica* | 1 | ITSAP23 | Carnica France | France | HiSeq |
| Wragg_et_al_2022_MER | SRR15173670 | SAMN20257199 | *A. mellifera* | *A.m. carnica* | 1 | ITSAP38 | Carnica France | France | HiSeq |
| Wragg_et_al_2022_MER | SRR15173667 | SAMN20257202 | *A. mellifera* | *A.m. carnica* | 1 | ITSAP42 | Carnica France | France | HiSeq |
| Wragg_et_al_2022_MER | SRR15173661 | SAMN20257207 | *A. mellifera* | *A.m. carnica* | 1 | ITSAP48 | Carnica France | France | HiSeq |
| Wragg_et_al_2022_MER | SRR15173660 | SAMN20257208 | *A. mellifera* | *A.m. carnica* | 1 | ITSAP49 | Carnica France | France | HiSeq |
| Wragg_et_al_2022_MER | SRR15173652 | SAMN20257215 | *A. mellifera* | *A.m. carnica* | 1 | ITSAP55 | Carnica France | France | HiSeq |
| Wragg_et_al_2022_MER | SRR15173650 | SAMN20257217 | *A. mellifera* | *A.m. carnica* | 1 | ITSAP60 | Carnica France | France | HiSeq |
| Wragg_et_al_2022_MER | SRR15173649 | SAMN20257218 | *A. mellifera* | *A.m. carnica* | 1 | ITSAP61 | Carnica France | France | HiSeq |
| Wragg_et_al_2022_MER | SRR15173647 | SAMN20257220 | *A. mellifera* | *A.m. carnica* | 1 | ITSAP63 | Carnica France | France | HiSeq |
| Wragg_et_al_2022_MER | SRR15173646 | SAMN20257221 | *A. mellifera* | *A.m. carnica* | 1 | ITSAP64 | Carnica France | France | HiSeq |
| Wragg_et_al_2022_MER | SRR15173618 | SAMN20257246 | *A. mellifera* | *A.m. carnica* | 1 | ITSAP9 | Carnica France | France | HiSeq |
| Wragg_et_al_2022_MER | SRR3157145 | SAMN04481240 | *A. mellifera* | *A.m. carnica* | 1 | BER10 | Carnica Germany | Germany | HiSeq |
| Wragg_et_al_2022_MER | SRR3157146 | SAMN04481241 | *A. mellifera* | *A.m. carnica* | 1 | BER11 | Carnica Germany | Germany | HiSeq |
| Wragg_et_al_2022_MER | SRR3157147 | SAMN04481242 | *A. mellifera* | *A.m. carnica* | 1 | BER12 | Carnica Germany | Germany | HiSeq |
| Wragg_et_al_2022_MER | SRR3157148 | SAMN04481243 | *A. mellifera* | *A.m. carnica* | 1 | BER13 | Carnica Germany | Germany | HiSeq |
| Wragg_et_al_2022_MER | SRR3157149 | SAMN04481244 | *A. mellifera* | *A.m. carnica* | 1 | BER14 | Carnica Germany | Germany | HiSeq |
| Wragg_et_al_2022_MER | SRR6236936 | SAMN07956261 | *A. mellifera* | *A.m. carnica* | 1 | BER15 | Carnica Germany | Germany | HiSeq |
| Wragg_et_al_2022_MER | SRR15173556 | SAMN20257011 | *A. mellifera* | *A.m. carnica* | 1 | BER16 | Carnica Germany | Germany | HiSeq |
| Wragg_et_al_2022_MER | SRR6236935 | SAMN07956262 | *A. mellifera* | *A.m. carnica* | 1 | BER17 | Carnica Germany | Germany | HiSeq |
| Wragg_et_al_2022_MER | SRR3157150 | SAMN04481245 | *A. mellifera* | *A.m. carnica* | 1 | BER18 | Carnica Germany | Germany | HiSeq |
| Wragg_et_al_2022_MER | SRR15173545 | SAMN20257012 | *A. mellifera* | *A.m. carnica* | 1 | BER19 | Carnica Germany | Germany | HiSeq |
| Wragg_et_al_2022_MER | SRR3157151 | SAMN04481246 | *A. mellifera* | *A.m. carnica* | 1 | BER2 | Carnica Germany | Germany | HiSeq |
| Wragg_et_al_2022_MER | SRR15173533 | SAMN20257013 | *A. mellifera* | *A.m. carnica* | 1 | BER21 | Carnica Germany | Germany | HiSeq |
| Wragg_et_al_2022_MER | SRR3157152 | SAMN04481247 | *A. mellifera* | *A.m. carnica* | 1 | BER4 | Carnica Germany | Germany | HiSeq |
| Wragg_et_al_2022_MER | SRR3157153 | SAMN04481248 | *A. mellifera* | *A.m. carnica* | 1 | BER5 | Carnica Germany | Germany | HiSeq |
| Wragg_et_al_2022_MER | SRR3157154 | SAMN04481249 | *A. mellifera* | *A.m. carnica* | 1 | BER6 | Carnica Germany | Germany | HiSeq |
| Wragg_et_al_2022_MER | SRR15173522 | SAMN20257014 | *A. mellifera* | *A.m. carnica* | 1 | BER7 | Carnica Germany | Germany | HiSeq |
| Wragg_et_al_2022_MER | SRR3157155 | SAMN04481250 | *A. mellifera* | *A.m. carnica* | 1 | BER8 | Carnica Germany | Germany | HiSeq |
| Wragg_et_al_2022_MER | SRR3157156 | SAMN04481251 | *A. mellifera* | *A.m. carnica* | 1 | BER9 | Carnica Germany | Germany | HiSeq |
| Wragg_et_al_2022_MER | SRR15173915 | SAMN20257314 | *A. mellifera* | *A.m. carnica* | 1 | POL11 | Carnica Poland | Poland | HiSeq |

| | | | | | | | | | |
|---|---|---|---|---|---|---|---|---|---|
| Wragg_et_al_2022_MER | SRR15173914 | SAMN20257315 | *A. mellifera* | *A.m. carnica* | 1 | POL12 | Carnica Poland | Poland | HiSeq |
| Wragg_et_al_2022_MER | SRR15173913 | SAMN20257316 | *A. mellifera* | *A.m. carnica* | 1 | POL13 | Carnica Poland | Poland | HiSeq |
| Wragg_et_al_2022_MER | SRR15173912 | SAMN20257317 | *A. mellifera* | *A.m. carnica* | 1 | POL14 | Carnica Poland | Poland | HiSeq |
| Wragg_et_al_2022_MER | SRR15173911 | SAMN20257318 | *A. mellifera* | *A.m. carnica* | 1 | POL15 | Carnica Poland | Poland | HiSeq |
| Wragg_et_al_2022_MER | SRR15173910 | SAMN20257319 | *A. mellifera* | *A.m. carnica* | 1 | POL19 | Carnica Poland | Poland | HiSeq |
| Wragg_et_al_2022_MER | SRR15173909 | SAMN20257320 | *A. mellifera* | *A.m. carnica* | 1 | POL2 | Carnica Poland | Poland | HiSeq |
| Wragg_et_al_2022_MER | SRR15173908 | SAMN20257321 | *A. mellifera* | *A.m. carnica* | 1 | POL21 | Carnica Poland | Poland | HiSeq |
| Wragg_et_al_2022_MER | SRR15173907 | SAMN20257322 | *A. mellifera* | *A.m. carnica* | 1 | POL23 | Carnica Poland | Poland | HiSeq |
| Wragg_et_al_2022_MER | SRR15173319 | SAMN20257323 | *A. mellifera* | *A.m. carnica* | 1 | POL25 | Carnica Poland | Poland | HiSeq |
| Wragg_et_al_2022_MER | SRR15173587 | SAMN20257324 | *A. mellifera* | *A.m. carnica* | 1 | POL26 | Carnica Poland | Poland | HiSeq |
| Wragg_et_al_2022_MER | SRR15173586 | SAMN20257325 | *A. mellifera* | *A.m. carnica* | 1 | POL27 | Carnica Poland | Poland | HiSeq |
| Wragg_et_al_2022_MER | SRR15173585 | SAMN20257326 | *A. mellifera* | *A.m. carnica* | 1 | POL28 | Carnica Poland | Poland | HiSeq |
| Wragg_et_al_2022_MER | SRR15173584 | SAMN20257327 | *A. mellifera* | *A.m. carnica* | 1 | POL3 | Carnica Poland | Poland | HiSeq |
| Wragg_et_al_2022_MER | SRR15173583 | SAMN20257328 | *A. mellifera* | *A.m. carnica* | 1 | POL30 | Carnica Poland | Poland | HiSeq |
| Wragg_et_al_2022_MER | SRR15173582 | SAMN20257329 | *A. mellifera* | *A.m. carnica* | 1 | POL5 | Carnica Poland | Poland | HiSeq |
| Wragg_et_al_2022_MER | SRR15173581 | SAMN20257330 | *A. mellifera* | *A.m. carnica* | 1 | POL6 | Carnica Poland | Poland | HiSeq |
| Wragg_et_al_2022_MER | SRR15173580 | SAMN20257331 | *A. mellifera* | *A.m. carnica* | 1 | POL7 | Carnica Poland | Poland | HiSeq |
| Wragg_et_al_2022_MER | SRR15173579 | SAMN20257332 | *A. mellifera* | *A.m. carnica* | 1 | POL9 | Carnica Poland | Poland | HiSeq |
| Wragg_et_al_2022_MER | SRR3157167 | SAMN04481262 | *A. mellifera* | *A.m. carnica* | 1 | SLO1 | Carnica Slovenia | Slovenia | HiSeq |
| Wragg_et_al_2022_MER | SRR3157157 | SAMN04481252 | *A. mellifera* | *A.m. carnica* | 1 | SLO10 | Carnica Slovenia | Slovenia | HiSeq |
| Wragg_et_al_2022_MER | SRR3157158 | SAMN04481253 | *A. mellifera* | *A.m. carnica* | 1 | SLO11 | Carnica Slovenia | Slovenia | HiSeq |
| Wragg_et_al_2022_MER | SRR3157159 | SAMN04481254 | *A. mellifera* | *A.m. carnica* | 1 | SLO12 | Carnica Slovenia | Slovenia | HiSeq |
| Wragg_et_al_2022_MER | SRR3157160 | SAMN04481255 | *A. mellifera* | *A.m. carnica* | 1 | SLO13 | Carnica Slovenia | Slovenia | HiSeq |
| Wragg_et_al_2022_MER | SRR3157161 | SAMN04481256 | *A. mellifera* | *A.m. carnica* | 1 | SLO14 | Carnica Slovenia | Slovenia | HiSeq |
| Wragg_et_al_2022_MER | SRR3157162 | SAMN04481257 | *A. mellifera* | *A.m. carnica* | 1 | SLO15 | Carnica Slovenia | Slovenia | HiSeq |
| Wragg_et_al_2022_MER | SRR3157163 | SAMN04481258 | *A. mellifera* | *A.m. carnica* | 1 | SLO16 | Carnica Slovenia | Slovenia | HiSeq |
| Wragg_et_al_2022_MER | SRR3157164 | SAMN04481259 | *A. mellifera* | *A.m. carnica* | 1 | SLO17 | Carnica Slovenia | Slovenia | HiSeq |
| Wragg_et_al_2022_MER | SRR3157165 | SAMN04481260 | *A. mellifera* | *A.m. carnica* | 1 | SLO18 | Carnica Slovenia | Slovenia | HiSeq |
| Wragg_et_al_2022_MER | SRR3157166 | SAMN04481261 | *A. mellifera* | *A.m. carnica* | 1 | SLO19 | Carnica Slovenia | Slovenia | HiSeq |
| Wragg_et_al_2022_MER | SRR3157169 | SAMN04481264 | *A. mellifera* | *A.m. carnica* | 1 | SLO2 | Carnica Slovenia | Slovenia | HiSeq |
| Wragg_et_al_2022_MER | SRR3157168 | SAMN04481263 | *A. mellifera* | *A.m. carnica* | 1 | SLO20 | Carnica Slovenia | Slovenia | HiSeq |
| Wragg_et_al_2022_MER | SRR3157170 | SAMN04481265 | *A. mellifera* | *A.m. carnica* | 1 | SLO3 | Carnica Slovenia | Slovenia | HiSeq |
| Wragg_et_al_2022_MER | SRR3157171 | SAMN04481266 | *A. mellifera* | *A.m. carnica* | 1 | SLO4 | Carnica Slovenia | Slovenia | HiSeq |
| Wragg_et_al_2022_MER | SRR3157172 | SAMN04481267 | *A. mellifera* | *A.m. carnica* | 1 | SLO5 | Carnica Slovenia | Slovenia | HiSeq |
| Wragg_et_al_2022_MER | SRR3157173 | SAMN04481268 | *A. mellifera* | *A.m. carnica* | 1 | SLO6 | Carnica Slovenia | Slovenia | HiSeq |
| Wragg_et_al_2022_MER | SRR3157174 | SAMN04481269 | *A. mellifera* | *A.m. carnica* | 1 | SLO7 | Carnica Slovenia | Slovenia | HiSeq |
| Wragg_et_al_2022_MER | SRR15173544 | SAMN20257363 | *A. mellifera* | *A.m. carnica* | 1 | SLO8 | Carnica Slovenia | Slovenia | HiSeq |
| Wragg_et_al_2022_MER | SRR15173543 | SAMN20257364 | *A. mellifera* | *A.m. carnica* | 1 | SLO9 | Carnica Slovenia | Slovenia | HiSeq |
| Wragg_et_al_2022_MER | ERR1700875 | SAMEA4521121 | *A. mellifera* | *A.m. carnica* | 1 | CAR10 | Carnica Switzerland | Switzerland | HiSeq |
| Wragg_et_al_2022_MER | ERR1700874 | SAMEA4521122 | *A. mellifera* | *A.m. carnica* | 1 | CAR11 | Carnica Switzerland | Switzerland | HiSeq |
| Wragg_et_al_2022_MER | ERR1700873 | SAMEA4521123 | *A. mellifera* | *A.m. carnica* | 1 | CAR12 | Carnica Switzerland | Switzerland | HiSeq |
| Wragg_et_al_2022_MER | ERR1700872 | SAMEA4521124 | *A. mellifera* | *A.m. carnica* | 1 | CAR13 | Carnica Switzerland | Switzerland | HiSeq |
| Wragg_et_al_2022_MER | ERR1700871 | SAMEA4521125 | *A. mellifera* | *A.m. carnica* | 1 | CAR14 | Carnica Switzerland | Switzerland | HiSeq |

| | | | | | | | | | |
|---|---|---|---|---|---|---|---|---|---|
| Wragg_et_al_2022_MER | ERR1700870 | SAMEA4521126 | *A. mellifera* | *A.m. carnica* | 1 | CAR15 | Carnica Switzerland | Switzerland | HiSeq |
| Wragg_et_al_2022_MER | ERR1700869 | SAMEA4521127 | *A. mellifera* | *A.m. carnica* | 1 | CAR16 | Carnica Switzerland | Switzerland | HiSeq |
| Wragg_et_al_2022_MER | ERR1700868 | SAMEA4521128 | *A. mellifera* | *A.m. carnica* | 1 | CAR17 | Carnica Switzerland | Switzerland | HiSeq |
| Wragg_et_al_2022_MER | ERR1700867 | SAMEA4521129 | *A. mellifera* | *A.m. carnica* | 1 | CAR18 | Carnica Switzerland | Switzerland | HiSeq |
| Wragg_et_al_2022_MER | ERR1700866 | SAMEA4521130 | *A. mellifera* | *A.m. carnica* | 1 | CAR19 | Carnica Switzerland | Switzerland | HiSeq |
| Wragg_et_al_2022_MER | ERR1700865 | SAMEA4521131 | *A. mellifera* | *A.m. carnica* | 1 | CAR2 | Carnica Switzerland | Switzerland | HiSeq |
| Wragg_et_al_2022_MER | ERR1700864 | SAMEA4521132 | *A. mellifera* | *A.m. carnica* | 1 | CAR20 | Carnica Switzerland | Switzerland | HiSeq |
| Wragg_et_al_2022_MER | ERR1700863 | SAMEA4521133 | *A. mellifera* | *A.m. carnica* | 1 | CAR21 | Carnica Switzerland | Switzerland | HiSeq |
| Wragg_et_al_2022_MER | ERR1700862 | SAMEA4521134 | *A. mellifera* | *A.m. carnica* | 1 | CAR22 | Carnica Switzerland | Switzerland | HiSeq |
| Wragg_et_al_2022_MER | ERR1700861 | SAMEA4521135 | *A. mellifera* | *A.m. carnica* | 1 | CAR23 | Carnica Switzerland | Switzerland | HiSeq |
| Wragg_et_al_2022_MER | ERR1700860 | SAMEA4521136 | *A. mellifera* | *A.m. carnica* | 1 | CAR25 | Carnica Switzerland | Switzerland | HiSeq |
| Wragg_et_al_2022_MER | ERR1700859 | SAMEA4521137 | *A. mellifera* | *A.m. carnica* | 1 | CAR26 | Carnica Switzerland | Switzerland | HiSeq |
| Wragg_et_al_2022_MER | ERR1700858 | SAMEA4521138 | *A. mellifera* | *A.m. carnica* | 1 | CAR28 | Carnica Switzerland | Switzerland | HiSeq |
| Wragg_et_al_2022_MER | ERR1700857 | SAMEA4521139 | *A. mellifera* | *A.m. carnica* | 1 | CAR29 | Carnica Switzerland | Switzerland | HiSeq |
| Wragg_et_al_2022_MER | ERR1700856 | SAMEA4521140 | *A. mellifera* | *A.m. carnica* | 1 | CAR3 | Carnica Switzerland | Switzerland | HiSeq |
| Wragg_et_al_2022_MER | ERR1700855 | SAMEA4521141 | *A. mellifera* | *A.m. carnica* | 1 | CAR30 | Carnica Switzerland | Switzerland | HiSeq |
| Wragg_et_al_2022_MER | ERR1700854 | SAMEA4521142 | *A. mellifera* | *A.m. carnica* | 1 | CAR31 | Carnica Switzerland | Switzerland | HiSeq |
| Wragg_et_al_2022_MER | ERR1700853 | SAMEA4521143 | *A. mellifera* | *A.m. carnica* | 1 | CAR32 | Carnica Switzerland | Switzerland | HiSeq |
| Wragg_et_al_2022_MER | ERR1700852 | SAMEA4521144 | *A. mellifera* | *A.m. carnica* | 1 | CAR33 | Carnica Switzerland | Switzerland | HiSeq |
| Wragg_et_al_2022_MER | ERR1700851 | SAMEA4521145 | *A. mellifera* | *A.m. carnica* | 1 | CAR34 | Carnica Switzerland | Switzerland | HiSeq |
| Wragg_et_al_2022_MER | ERR1700850 | SAMEA4521146 | *A. mellifera* | *A.m. carnica* | 1 | CAR4 | Carnica Switzerland | Switzerland | HiSeq |
| Wragg_et_al_2022_MER | ERR1700849 | SAMEA4521147 | *A. mellifera* | *A.m. carnica* | 1 | CAR5 | Carnica Switzerland | Switzerland | HiSeq |
| Wragg_et_al_2022_MER | ERR1700848 | SAMEA4521148 | *A. mellifera* | *A.m. carnica* | 1 | CAR6 | Carnica Switzerland | Switzerland | HiSeq |
| Wragg_et_al_2022_MER | ERR1700847 | SAMEA4521149 | *A. mellifera* | *A.m. carnica* | 1 | CAR7 | Carnica Switzerland | Switzerland | HiSeq |
| Wragg_et_al_2022_MER | ERR1700846 | SAMEA4521150 | *A. mellifera* | *A.m. carnica* | 1 | CAR8 | Carnica Switzerland | Switzerland | HiSeq |
| Wragg_et_al_2022_MER | ERR1700845 | SAMEA4521151 | *A. mellifera* | *A.m. carnica* | 1 | CAR9 | Carnica Switzerland | Switzerland | HiSeq |
| Wragg_et_al_2022_MER | SRR6236934 | SAMN07956263 | *A. mellifera* | *A.m. caucasia* | 1 | CAU10 | Caucasia France | France | HiSeq |
| Wragg_et_al_2022_MER | SRR6236933 | SAMN07956264 | *A. mellifera* | *A.m. caucasia* | 1 | CAU11 | Caucasia France | France | HiSeq |
| Wragg_et_al_2022_MER | SRR6236940 | SAMN07956265 | *A. mellifera* | *A.m. caucasia* | 1 | CAU12 | Caucasia France | France | HiSeq |
| Wragg_et_al_2022_MER | SRR6236939 | SAMN07956266 | *A. mellifera* | *A.m. caucasia* | 1 | CAU14 | Caucasia France | France | HiSeq |
| Wragg_et_al_2022_MER | SRR15173877 | SAMN20257066 | *A. mellifera* | *A.m. caucasia* | 1 | CAU15A | Caucasia France | France | HiSeq |
| Wragg_et_al_2022_MER | SRR15173876 | SAMN20257067 | *A. mellifera* | *A.m. caucasia* | 1 | CAU16A | Caucasia France | France | HiSeq |
| Wragg_et_al_2022_MER | SRR6236938 | SAMN07956267 | *A. mellifera* | *A.m. caucasia* | 1 | CAU17 | Caucasia France | France | HiSeq |
| Wragg_et_al_2022_MER | SRR15173875 | SAMN20257068 | *A. mellifera* | *A.m. caucasia* | 1 | CAU17A | Caucasia France | France | HiSeq |
| Wragg_et_al_2022_MER | SRR15173874 | SAMN20257069 | *A. mellifera* | *A.m. caucasia* | 1 | CAU18A | Caucasia France | France | HiSeq |
| Wragg_et_al_2022_MER | SRR15173873 | SAMN20257070 | *A. mellifera* | *A.m. caucasia* | 1 | CAU19A | Caucasia France | France | HiSeq |
| Wragg_et_al_2022_MER | SRR6236944 | SAMN07956268 | *A. mellifera* | *A.m. caucasia* | 1 | CAU20 | Caucasia France | France | HiSeq |
| Wragg_et_al_2022_MER | SRR6236943 | SAMN07956269 | *A. mellifera* | *A.m. caucasia* | 1 | CAU21 | Caucasia France | France | HiSeq |
| Wragg_et_al_2022_MER | SRR6236969 | SAMN07956270 | *A. mellifera* | *A.m. caucasia* | 1 | CAU5 | Caucasia France | France | HiSeq |
| Wragg_et_al_2022_MER | SRR6236946 | SAMN07956271 | *A. mellifera* | *A.m. caucasia* | 1 | CAU6 | Caucasia France | France | HiSeq |
| Wragg_et_al_2022_MER | SRR6236945 | SAMN07956272 | *A. mellifera* | *A.m. caucasia* | 1 | CAU7 | Caucasia France | France | HiSeq |
| Wragg_et_al_2022_MER | SRR15173872 | SAMN20257071 | *A. mellifera* | Unknown | 1 | CHI10A | China | China | HiSeq |
| Wragg_et_al_2022_MER | SRR15173871 | SAMN20257072 | *A. mellifera* | Unknown | 1 | CHI1A | China | China | HiSeq |

| | | | | | | | | | |
|---|---|---|---|---|---|---|---|---|---|
| Wragg_et_al_2022_MER | SRR15173809 | SAMN20257073 | *A. mellifera* | Unknown | 1 | CHI2A | China | China | HiSeq |
| Wragg_et_al_2022_MER | SRR15173808 | SAMN20257074 | *A. mellifera* | Unknown | 1 | CHI3A | China | China | HiSeq |
| Wragg_et_al_2022_MER | SRR15173807 | SAMN20257075 | *A. mellifera* | Unknown | 1 | CHI4A | China | China | HiSeq |
| Wragg_et_al_2022_MER | SRR15173806 | SAMN20257076 | *A. mellifera* | Unknown | 1 | CHI5A | China | China | HiSeq |
| Wragg_et_al_2022_MER | SRR15173805 | SAMN20257077 | *A. mellifera* | Unknown | 1 | CHI6A | China | China | HiSeq |
| Wragg_et_al_2022_MER | SRR15173804 | SAMN20257078 | *A. mellifera* | Unknown | 1 | CHI7A | China | China | HiSeq |
| Wragg_et_al_2022_MER | SRR15173803 | SAMN20257079 | *A. mellifera* | Unknown | 1 | CHI8A | China | China | HiSeq |
| Wragg_et_al_2022_MER | SRR15173802 | SAMN20257080 | *A. mellifera* | Unknown | 1 | CHI9A | China | China | HiSeq |
| Wragg_et_al_2022_MER | SRR15173525 | SAMN20257380 | *A. mellifera* | *A.m.mellifera* | 1 | UK10A | Colonsay Conservatory | Scotland | HiSeq |
| Wragg_et_al_2022_MER | SRR15173524 | SAMN20257381 | *A. mellifera* | *A.m.mellifera* | 1 | UK11A | Colonsay Conservatory | Scotland | HiSeq |
| Wragg_et_al_2022_MER | SRR15173523 | SAMN20257382 | *A. mellifera* | *A.m.mellifera* | 1 | UK12A | Colonsay Conservatory | Scotland | HiSeq |
| Wragg_et_al_2022_MER | SRR15173521 | SAMN20257383 | *A. mellifera* | *A.m.mellifera* | 1 | UK13A | Colonsay Conservatory | Scotland | HiSeq |
| Wragg_et_al_2022_MER | SRR15173520 | SAMN20257384 | *A. mellifera* | *A.m.mellifera* | 1 | UK14A | Colonsay Conservatory | Scotland | HiSeq |
| Wragg_et_al_2022_MER | SRR15173519 | SAMN20257385 | *A. mellifera* | *A.m.mellifera* | 1 | UK15A | Colonsay Conservatory | Scotland | HiSeq |
| Wragg_et_al_2022_MER | SRR15173518 | SAMN20257386 | *A. mellifera* | *A.m.mellifera* | 1 | UK16A | Colonsay Conservatory | Scotland | HiSeq |
| Wragg_et_al_2022_MER | SRR15173517 | SAMN20257387 | *A. mellifera* | *A.m.mellifera* | 1 | UK17A | Colonsay Conservatory | Scotland | HiSeq |
| Wragg_et_al_2022_MER | SRR15173516 | SAMN20257388 | *A. mellifera* | *A.m.mellifera* | 1 | UK18A | Colonsay Conservatory | Scotland | HiSeq |
| Wragg_et_al_2022_MER | SRR15173515 | SAMN20257389 | *A. mellifera* | *A.m.mellifera* | 1 | UK19A | Colonsay Conservatory | Scotland | HiSeq |
| Wragg_et_al_2022_MER | SRR15173514 | SAMN20257390 | *A. mellifera* | *A.m.mellifera* | 1 | UK1A | Colonsay Conservatory | Scotland | HiSeq |
| Wragg_et_al_2022_MER | SRR15173513 | SAMN20257391 | *A. mellifera* | *A.m.mellifera* | 1 | UK20A | Colonsay Conservatory | Scotland | HiSeq |
| Wragg_et_al_2022_MER | SRR15173512 | SAMN20257392 | *A. mellifera* | *A.m.mellifera* | 1 | UK21A | Colonsay Conservatory | Scotland | HiSeq |
| Wragg_et_al_2022_MER | SRR15173510 | SAMN20257393 | *A. mellifera* | *A.m.mellifera* | 1 | UK22A | Colonsay Conservatory | Scotland | HiSeq |
| Wragg_et_al_2022_MER | SRR15173509 | SAMN20257394 | *A. mellifera* | *A.m.mellifera* | 1 | UK23A | Colonsay Conservatory | Scotland | HiSeq |
| Wragg_et_al_2022_MER | SRR15173508 | SAMN20257395 | *A. mellifera* | *A.m.mellifera* | 1 | UK25A | Colonsay Conservatory | Scotland | HiSeq |
| Wragg_et_al_2022_MER | SRR15173507 | SAMN20257396 | *A. mellifera* | *A.m.mellifera* | 1 | UK26A | Colonsay Conservatory | Scotland | HiSeq |
| Wragg_et_al_2022_MER | SRR15173506 | SAMN20257397 | *A. mellifera* | *A.m.mellifera* | 1 | UK27A | Colonsay Conservatory | Scotland | HiSeq |
| Wragg_et_al_2022_MER | SRR15173505 | SAMN20257398 | *A. mellifera* | *A.m.mellifera* | 1 | UK28A | Colonsay Conservatory | Scotland | HiSeq |
| Wragg_et_al_2022_MER | SRR15173504 | SAMN20257399 | *A. mellifera* | *A.m.mellifera* | 1 | UK29A | Colonsay Conservatory | Scotland | HiSeq |
| Wragg_et_al_2022_MER | SRR15173503 | SAMN20257400 | *A. mellifera* | *A.m.mellifera* | 1 | UK2A | Colonsay Conservatory | Scotland | HiSeq |
| Wragg_et_al_2022_MER | SRR15173502 | SAMN20257401 | *A. mellifera* | *A.m.mellifera* | 1 | UK3A | Colonsay Conservatory | Scotland | HiSeq |
| Wragg_et_al_2022_MER | SRR15173501 | SAMN20257402 | *A. mellifera* | *A.m.mellifera* | 1 | UK4A | Colonsay Conservatory | Scotland | HiSeq |
| Wragg_et_al_2022_MER | SRR15173499 | SAMN20257403 | *A. mellifera* | *A.m.mellifera* | 1 | UK5A | Colonsay Conservatory | Scotland | HiSeq |
| Wragg_et_al_2022_MER | SRR15173498 | SAMN20257404 | *A. mellifera* | *A.m.mellifera* | 1 | UK6A | Colonsay Conservatory | Scotland | HiSeq |
| Wragg_et_al_2022_MER | SRR15173497 | SAMN20257405 | *A. mellifera* | *A.m.mellifera* | 1 | UK7A | Colonsay Conservatory | Scotland | HiSeq |
| Wragg_et_al_2022_MER | SRR15173496 | SAMN20257406 | *A. mellifera* | *A.m.mellifera* | 1 | UK8A | Colonsay | Scotland | HiSeq |

| | | | | | | | | | |
|---|---|---|---|---|---|---|---|---|---|
| | | | | | | | Conservatory | | |
| Wragg_et_al_2022_MER | SRR15173495 | SAMN20257407 | *A. mellifera* | *A.m.mellifera* | 1 | UK9A | Colonsay Conservatory | Scotland | HiSeq |
| Wragg_et_al_2022_MER | SRR15173859 | SAMN20256974 | *A. mellifera* | Unknown | 1 | AOC10 | Corsica Breeder | Corsica | HiSeq |
| Wragg_et_al_2022_MER | SRR15173700 | SAMN20256975 | *A. mellifera* | Unknown | 1 | AOC11 | Corsica Breeder | Corsica | HiSeq |
| Wragg_et_al_2022_MER | SRR15173589 | SAMN20256976 | *A. mellifera* | Unknown | 1 | AOC12 | Corsica Breeder | Corsica | HiSeq |
| Wragg_et_al_2022_MER | SRR15173534 | SAMN20256977 | *A. mellifera* | Unknown | 1 | AOC14 | Corsica Breeder | Corsica | HiSeq |
| Wragg_et_al_2022_MER | SRR15173423 | SAMN20256978 | *A. mellifera* | Unknown | 1 | AOC15 | Corsica Breeder | Corsica | HiSeq |
| Wragg_et_al_2022_MER | SRR15173865 | SAMN20256979 | *A. mellifera* | Unknown | 1 | AOC16 | Corsica Breeder | Corsica | HiSeq |
| Wragg_et_al_2022_MER | SRR15173904 | SAMN20256980 | *A. mellifera* | Unknown | 1 | AOC17 | Corsica Breeder | Corsica | HiSeq |
| Wragg_et_al_2022_MER | SRR15173893 | SAMN20256981 | *A. mellifera* | Unknown | 1 | AOC18 | Corsica Breeder | Corsica | HiSeq |
| Wragg_et_al_2022_MER | SRR15173882 | SAMN20256982 | *A. mellifera* | Unknown | 1 | AOC19 | Corsica Breeder | Corsica | HiSeq |
| Wragg_et_al_2022_MER | SRR15173858 | SAMN20256983 | *A. mellifera* | Unknown | 1 | AOC2 | Corsica Breeder | Corsica | HiSeq |
| Wragg_et_al_2022_MER | SRR15173799 | SAMN20256984 | *A. mellifera* | Unknown | 1 | AOC20 | Corsica Breeder | Corsica | HiSeq |
| Wragg_et_al_2022_MER | SRR15173788 | SAMN20256985 | *A. mellifera* | Unknown | 1 | AOC21 | Corsica Breeder | Corsica | HiSeq |
| Wragg_et_al_2022_MER | SRR15173777 | SAMN20256986 | *A. mellifera* | Unknown | 1 | AOC22 | Corsica Breeder | Corsica | HiSeq |
| Wragg_et_al_2022_MER | SRR15173766 | SAMN20256987 | *A. mellifera* | Unknown | 1 | AOC25 | Corsica Breeder | Corsica | HiSeq |
| Wragg_et_al_2022_MER | SRR15173755 | SAMN20256988 | *A. mellifera* | Unknown | 1 | AOC26 | Corsica Breeder | Corsica | HiSeq |
| Wragg_et_al_2022_MER | SRR15173744 | SAMN20256989 | *A. mellifera* | Unknown | 1 | AOC27 | Corsica Breeder | Corsica | HiSeq |
| Wragg_et_al_2022_MER | SRR15173733 | SAMN20256990 | *A. mellifera* | Unknown | 1 | AOC28 | Corsica Breeder | Corsica | HiSeq |
| Wragg_et_al_2022_MER | SRR15173722 | SAMN20256991 | *A. mellifera* | Unknown | 1 | AOC29 | Corsica Breeder | Corsica | HiSeq |
| Wragg_et_al_2022_MER | SRR15173711 | SAMN20256992 | *A. mellifera* | Unknown | 1 | AOC3 | Corsica Breeder | Corsica | HiSeq |
| Wragg_et_al_2022_MER | SRR15173699 | SAMN20256993 | *A. mellifera* | Unknown | 1 | AOC31 | Corsica Breeder | Corsica | HiSeq |
| Wragg_et_al_2022_MER | SRR15173688 | SAMN20256994 | *A. mellifera* | Unknown | 1 | AOC32 | Corsica Breeder | Corsica | HiSeq |
| Wragg_et_al_2022_MER | SRR15173677 | SAMN20256995 | *A. mellifera* | Unknown | 1 | AOC33 | Corsica Breeder | Corsica | HiSeq |
| Wragg_et_al_2022_MER | SRR15173666 | SAMN20256996 | *A. mellifera* | Unknown | 1 | AOC34 | Corsica Breeder | Corsica | HiSeq |
| Wragg_et_al_2022_MER | SRR15173655 | SAMN20256997 | *A. mellifera* | Unknown | 1 | AOC35 | Corsica Breeder | Corsica | HiSeq |
| Wragg_et_al_2022_MER | SRR15173644 | SAMN20256998 | *A. mellifera* | Unknown | 1 | AOC36 | Corsica Breeder | Corsica | HiSeq |
| Wragg_et_al_2022_MER | SRR15173633 | SAMN20256999 | *A. mellifera* | Unknown | 1 | AOC37 | Corsica Breeder | Corsica | HiSeq |
| Wragg_et_al_2022_MER | SRR15173622 | SAMN20257000 | *A. mellifera* | Unknown | 1 | AOC38 | Corsica Breeder | Corsica | HiSeq |
| Wragg_et_al_2022_MER | SRR15173611 | SAMN20257001 | *A. mellifera* | Unknown | 1 | AOC39 | Corsica Breeder | Corsica | HiSeq |
| Wragg_et_al_2022_MER | SRR15173588 | SAMN20257003 | *A. mellifera* | Unknown | 1 | AOC40 | Corsica Breeder | Corsica | HiSeq |
| Wragg_et_al_2022_MER | SRR15173950 | SAMN20257004 | *A. mellifera* | Unknown | 1 | AOC41 | Corsica Breeder | Corsica | HiSeq |
| Wragg_et_al_2022_MER | SRR15173939 | SAMN20257005 | *A. mellifera* | Unknown | 1 | AOC42 | Corsica Breeder | Corsica | HiSeq |
| Wragg_et_al_2022_MER | SRR15173928 | SAMN20257006 | *A. mellifera* | Unknown | 1 | AOC5 | Corsica Breeder | Corsica | HiSeq |
| Wragg_et_al_2022_MER | SRR15173917 | SAMN20257007 | *A. mellifera* | Unknown | 1 | AOC6 | Corsica Breeder | Corsica | HiSeq |
| Wragg_et_al_2022_MER | SRR15173906 | SAMN20257008 | *A. mellifera* | Unknown | 1 | AOC7 | Corsica Breeder | Corsica | HiSeq |
| Wragg_et_al_2022_MER | SRR15173578 | SAMN20257009 | *A. mellifera* | Unknown | 1 | AOC8 | Corsica Breeder | Corsica | HiSeq |
| Wragg_et_al_2022_MER | SRR15173567 | SAMN20257010 | *A. mellifera* | Unknown | 1 | AOC9 | Corsica Breeder | Corsica | HiSeq |
| Wragg_et_al_2022_MER | SRR15173704 | SAMN20257169 | *A. mellifera* | Unknown | 1 | ITSAP104 | Corsica Breeder | Corsica | HiSeq |
| Wragg_et_al_2022_MER | SRR15173671 | SAMN20257198 | *A. mellifera* | Unknown | 1 | ITSAP37 | Corsica Breeder | Corsica | HiSeq |
| Wragg_et_al_2022_MER | SRR15173654 | SAMN20257213 | *A. mellifera* | Unknown | 1 | ITSAP53 | Corsica Breeder | Corsica | HiSeq |
| Wragg_et_al_2022_MER | SRR15173634 | SAMN20257232 | *A. mellifera* | Unknown | 1 | ITSAP73 | Corsica Breeder | Corsica | HiSeq |

| | | | | | | | | | |
|---|---|---|---|---|---|---|---|---|---|
| Wragg_et_al_2022_MER | SRR15173626 | SAMN20257239 | *A. mellifera* | Unknown | 1 | ITSAP83 | Corsica Breeder | Corsica | HiSeq |
| Wragg_et_al_2022_MER | SRR15173625 | SAMN20257240 | *A. mellifera* | Unknown | 1 | ITSAP84 | Corsica Breeder | Corsica | HiSeq |
| Wragg_et_al_2022_MER | SRR15173527 | SAMN20257378 | *A. mellifera* | Unknown | 1 | TES19 | Corsica Breeder | Corsica | HiSeq |
| Wragg_et_al_2022_MER | SRR15173609 | SAMN20257254 | *A. mellifera* | Unknown | 1 | JSC1 | Hautes Pyrenees Breeder | Hautes Pyrenees, France | HiSeq |
| Wragg_et_al_2022_MER | SRR15173608 | SAMN20257255 | *A. mellifera* | Unknown | 1 | JSC10 | Hautes Pyrenees Breeder | Hautes Pyrenees, France | HiSeq |
| Wragg_et_al_2022_MER | SRR15173607 | SAMN20257256 | *A. mellifera* | Unknown | 1 | JSC11 | Hautes Pyrenees Breeder | Hautes Pyrenees, France | HiSeq |
| Wragg_et_al_2022_MER | SRR15173606 | SAMN20257257 | *A. mellifera* | Unknown | 1 | JSC12 | Hautes Pyrenees Breeder | Hautes Pyrenees, France | HiSeq |
| Wragg_et_al_2022_MER | SRR15173605 | SAMN20257258 | *A. mellifera* | Unknown | 1 | JSC13 | Hautes Pyrenees Breeder | Hautes Pyrenees, France | HiSeq |
| Wragg_et_al_2022_MER | SRR15173604 | SAMN20257259 | *A. mellifera* | Unknown | 1 | JSC14 | Hautes Pyrenees Breeder | Hautes Pyrenees, France | HiSeq |
| Wragg_et_al_2022_MER | SRR15173603 | SAMN20257260 | *A. mellifera* | Unknown | 1 | JSC15 | Hautes Pyrenees Breeder | Hautes Pyrenees, France | HiSeq |
| Wragg_et_al_2022_MER | SRR15173602 | SAMN20257261 | *A. mellifera* | Unknown | 1 | JSC16 | Hautes Pyrenees Breeder | Hautes Pyrenees, France | HiSeq |
| Wragg_et_al_2022_MER | SRR15173601 | SAMN20257262 | *A. mellifera* | Unknown | 1 | JSC17 | Hautes Pyrenees Breeder | Hautes Pyrenees, France | HiSeq |
| Wragg_et_al_2022_MER | SRR15173599 | SAMN20257263 | *A. mellifera* | Unknown | 1 | JSC18 | Hautes Pyrenees Breeder | Hautes Pyrenees, France | HiSeq |
| Wragg_et_al_2022_MER | SRR15173598 | SAMN20257264 | *A. mellifera* | Unknown | 1 | JSC19 | Hautes Pyrenees Breeder | Hautes Pyrenees, France | HiSeq |
| Wragg_et_al_2022_MER | SRR15173597 | SAMN20257265 | *A. mellifera* | Unknown | 1 | JSC2 | Hautes Pyrenees Breeder | Hautes Pyrenees, France | HiSeq |
| Wragg_et_al_2022_MER | SRR15173596 | SAMN20257266 | *A. mellifera* | Unknown | 1 | JSC20 | Hautes Pyrenees Breeder | Hautes Pyrenees, France | HiSeq |
| Wragg_et_al_2022_MER | SRR15173595 | SAMN20257267 | *A. mellifera* | Unknown | 1 | JSC3 | Hautes Pyrenees Breeder | Hautes Pyrenees, France | HiSeq |
| Wragg_et_al_2022_MER | SRR15173594 | SAMN20257268 | *A. mellifera* | Unknown | 1 | JSC4 | Hautes Pyrenees Breeder | Hautes Pyrenees, France | HiSeq |
| Wragg_et_al_2022_MER | SRR15173593 | SAMN20257269 | *A. mellifera* | Unknown | 1 | JSC5 | Hautes Pyrenees Breeder | Hautes Pyrenees, France | HiSeq |
| Wragg_et_al_2022_MER | SRR15173592 | SAMN20257270 | *A. mellifera* | Unknown | 1 | JSC7 | Hautes Pyrenees Breeder | Hautes Pyrenees, France | HiSeq |
| Wragg_et_al_2022_MER | SRR15173591 | SAMN20257271 | *A. mellifera* | Unknown | 1 | JSC8 | Hautes Pyrenees Breeder | Hautes Pyrenees, France | HiSeq |
| Wragg_et_al_2022_MER | SRR15173590 | SAMN20257272 | *A. mellifera* | Unknown | 1 | JSC9 | Hautes Pyrenees Breeder | Hautes Pyrenees, France | HiSeq |
| Wragg_et_al_2022_MER | SRR15173709 | SAMN20257164 | *A. mellifera* | Unknown | 1 | ITSAP10 | Herault Breeder | Herault, France | HiSeq |
| Wragg_et_al_2022_MER | SRR15173657 | SAMN20257211 | *A. mellifera* | Unknown | 1 | ITSAP51 | Herault Breeder | Herault, France | HiSeq |
| Wragg_et_al_2022_MER | SRR15173656 | SAMN20257212 | *A. mellifera* | Unknown | 1 | ITSAP52 | Herault Breeder | Herault, France | HiSeq |
| Wragg_et_al_2022_MER | SRR15173639 | SAMN20257227 | *A. mellifera* | Unknown | 1 | ITSAP6B | Herault Breeder | Herault, France | HiSeq |
| Wragg_et_al_2022_MER | SRR15173638 | SAMN20257228 | *A. mellifera* | Unknown | 1 | ITSAP7 | Herault Breeder | Herault, France | HiSeq |
| Wragg_et_al_2022_MER | SRR15173636 | SAMN20257230 | *A. mellifera* | Unknown | 1 | ITSAP71 | Herault Breeder | Herault, France | HiSeq |
| Wragg_et_al_2022_MER | SRR15173623 | SAMN20257242 | *A. mellifera* | Unknown | 1 | ITSAP86 | Herault Breeder | Herault, France | HiSeq |
| Wragg_et_al_2022_MER | SRR15173775 | SAMN20257104 | *A. mellifera* | *A. m. iberiensis* | 1 | ESP10 | Iberiensis Spain | Spain Central Transect 1 | HiSeq |
| Wragg_et_al_2022_MER | SRR15173774 | SAMN20257105 | *A. mellifera* | *A. m. iberiensis* | 1 | ESP11 | Iberiensis Spain | Spain Central Transect 1 | HiSeq |
| Wragg_et_al_2022_MER | SRR15173773 | SAMN20257106 | *A. mellifera* | *A. m. iberiensis* | 1 | ESP12 | Iberiensis Spain | Spain Central Transect 1 | HiSeq |
| Wragg_et_al_2022_MER | SRR15173772 | SAMN20257107 | *A. mellifera* | *A. m. iberiensis* | 1 | ESP13 | Iberiensis Spain | Spain Central Transect 1 | HiSeq |
| Wragg_et_al_2022_MER | SRR15173771 | SAMN20257108 | *A. mellifera* | *A. m. iberiensis* | 1 | ESP14 | Iberiensis Spain | Spain Central Transect 1 | HiSeq |
| Wragg_et_al_2022_MER | SRR15173770 | SAMN20257109 | *A. mellifera* | *A. m. iberiensis* | 1 | ESP15 | Iberiensis Spain | Spain Central Transect 1 | HiSeq |

| | | | | | | | | | |
|---|---|---|---|---|---|---|---|---|---|
| Wragg_et_al_2022_MER | SRR15173769 | SAMN20257110 | *A. mellifera* | *A. m. iberiensis* | 1 | ESP16 | Iberiensis Spain | Spain Central Transect 1 | HiSeq |
| Wragg_et_al_2022_MER | SRR15173768 | SAMN20257111 | *A. mellifera* | *A. m. iberiensis* | 1 | ESP17 | Iberiensis Spain | Spain Central Transect 1 | HiSeq |
| Wragg_et_al_2022_MER | SRR15173767 | SAMN20257112 | *A. mellifera* | *A. m. iberiensis* | 1 | ESP18 | Iberiensis Spain | Spain Central Transect 1 | HiSeq |
| Wragg_et_al_2022_MER | SRR15173762 | SAMN20257116 | *A. mellifera* | *A. m. iberiensis* | 1 | ESP21 | Iberiensis Spain | Spain Central Transect 9 | HiSeq |
| Wragg_et_al_2022_MER | SRR15173761 | SAMN20257117 | *A. mellifera* | *A. m. iberiensis* | 1 | ESP22 | Iberiensis Spain | Spain Central Transect 9 | HiSeq |
| Wragg_et_al_2022_MER | SRR15173760 | SAMN20257118 | *A. mellifera* | *A. m. iberiensis* | 1 | ESP23 | Iberiensis Spain | Spain Central Transect 9 | HiSeq |
| Wragg_et_al_2022_MER | SRR15173759 | SAMN20257119 | *A. mellifera* | *A. m. iberiensis* | 1 | ESP24 | Iberiensis Spain | Spain Central Transect 9 | HiSeq |
| Wragg_et_al_2022_MER | SRR15173758 | SAMN20257120 | *A. mellifera* | *A. m. iberiensis* | 1 | ESP25 | Iberiensis Spain | Spain Central Transect 9 | HiSeq |
| Wragg_et_al_2022_MER | SRR15173757 | SAMN20257121 | *A. mellifera* | *A. m. iberiensis* | 1 | ESP26 | Iberiensis Spain | Spain Central Transect 9 | HiSeq |
| Wragg_et_al_2022_MER | SRR15173756 | SAMN20257122 | *A. mellifera* | *A. m. iberiensis* | 1 | ESP27 | Iberiensis Spain | Spain Central Transect 9 | HiSeq |
| Wragg_et_al_2022_MER | SRR15173754 | SAMN20257123 | *A. mellifera* | *A. m. iberiensis* | 1 | ESP28 | Iberiensis Spain | Spain Central Transect 9 | HiSeq |
| Wragg_et_al_2022_MER | SRR15173753 | SAMN20257124 | *A. mellifera* | *A. m. iberiensis* | 1 | ESP29 | Iberiensis Spain | Spain Central Transect 9 | HiSeq |
| Wragg_et_al_2022_MER | SRR15173751 | SAMN20257126 | *A. mellifera* | *A. m. iberiensis* | 1 | ESP30 | Iberiensis Spain | Spain Central Transect 9 | HiSeq |
| Wragg_et_al_2022_MER | SRR15173776 | SAMN20257103 | *A. mellifera* | *A. m. iberiensis* | 1 | ESP1 | Iberiensis Spain | Spain Mediterranean Transect 1 | HiSeq |
| Wragg_et_al_2022_MER | SRR15173765 | SAMN20257113 | *A. mellifera* | *A. m. iberiensis* | 1 | ESP19 | Iberiensis Spain | Spain Mediterranean Transect 1 | HiSeq |
| Wragg_et_al_2022_MER | SRR15173764 | SAMN20257114 | *A. mellifera* | *A. m. iberiensis* | 1 | ESP2 | Iberiensis Spain | Spain Mediterranean Transect 1 | HiSeq |
| Wragg_et_al_2022_MER | SRR15173763 | SAMN20257115 | *A. mellifera* | *A. m. iberiensis* | 1 | ESP20 | Iberiensis Spain | Spain Mediterranean Transect 1 | HiSeq |
| Wragg_et_al_2022_MER | SRR15173752 | SAMN20257125 | *A. mellifera* | *A. m. iberiensis* | 1 | ESP3 | Iberiensis Spain | Spain Mediterranean Transect 1 | HiSeq |
| Wragg_et_al_2022_MER | SRR15173750 | SAMN20257127 | *A. mellifera* | *A. m. iberiensis* | 1 | ESP4 | Iberiensis Spain | Spain Mediterranean Transect 1 | HiSeq |
| Wragg_et_al_2022_MER | SRR15173749 | SAMN20257128 | *A. mellifera* | *A. m. iberiensis* | 1 | ESP5 | Iberiensis Spain | Spain Mediterranean Transect 1 | HiSeq |
| Wragg_et_al_2022_MER | SRR15173748 | SAMN20257129 | *A. mellifera* | *A. m. iberiensis* | 1 | ESP6 | Iberiensis Spain | Spain Mediterranean Transect 1 | HiSeq |
| Wragg_et_al_2022_MER | SRR15173747 | SAMN20257130 | *A. mellifera* | *A. m. iberiensis* | 1 | ESP7 | Iberiensis Spain | Spain Mediterranean Transect 1 | HiSeq |
| Wragg_et_al_2022_MER | SRR15173746 | SAMN20257131 | *A. mellifera* | *A. m. iberiensis* | 1 | ESP8 | Iberiensis Spain | Spain Mediterranean Transect 1 | HiSeq |
| Wragg_et_al_2022_MER | SRR15173703 | SAMN20257170 | *A. mellifera* | Unknown | 1 | ITSAP105 | Isere 1 Breeder | Isere, France | HiSeq |
| Wragg_et_al_2022_MER | SRR15173690 | SAMN20257181 | *A. mellifera* | Unknown | 1 | ITSAP18 | Isere 1 Breeder | Isere, France | HiSeq |
| Wragg_et_al_2022_MER | SRR15173642 | SAMN20257224 | *A. mellifera* | Unknown | 1 | ITSAP67 | Isere 1 Breeder | Isere, France | HiSeq |
| Wragg_et_al_2022_MER | SRR15173641 | SAMN20257225 | *A. mellifera* | Unknown | 1 | ITSAP68 | Isere 1 Breeder | Isere, France | HiSeq |
| Wragg_et_al_2022_MER | SRR15173640 | SAMN20257226 | *A. mellifera* | Unknown | 1 | ITSAP69 | Isere 1 Breeder | Isere, France | HiSeq |
| Wragg_et_al_2022_MER | SRR15173610 | SAMN20257253 | *A. mellifera* | Unknown | 1 | ITSAP98 | Isere 1 Breeder | Isere, France | HiSeq |
| Wragg_et_al_2022_MER | SRR15173710 | SAMN20257163 | *A. mellifera* | Unknown | 1 | ITSAP1 | Isere 2 Breeder | Isere, France | HiSeq |
| Wragg_et_al_2022_MER | SRR15173680 | SAMN20257190 | *A. mellifera* | Unknown | 1 | ITSAP29 | Isere 2 Breeder | Isere, France | HiSeq |
| Wragg_et_al_2022_MER | SRR15173673 | SAMN20257196 | *A. mellifera* | Unknown | 1 | ITSAP35 | Isere 2 Breeder | Isere, France | HiSeq |
| Wragg_et_al_2022_MER | SRR15173672 | SAMN20257197 | *A. mellifera* | Unknown | 1 | ITSAP36 | Isere 2 Breeder | Isere, France | HiSeq |
| Wragg_et_al_2022_MER | SRR15173669 | SAMN20257200 | *A. mellifera* | Unknown | 1 | ITSAP39 | Isere 2 Breeder | Isere, France | HiSeq |
| Wragg_et_al_2022_MER | SRR15173665 | SAMN20257203 | *A. mellifera* | Unknown | 1 | ITSAP43 | Isere 2 Breeder | Isere, France | HiSeq |
| Wragg_et_al_2022_MER | SRR15173662 | SAMN20257206 | *A. mellifera* | Unknown | 1 | ITSAP47 | Isere 2 Breeder | Isere, France | HiSeq |

| | | | | | | | | | |
|---|---|---|---|---|---|---|---|---|---|
| Wragg_et_al_2022_MER | SRR15173620 | SAMN20257244 | *A. mellifera* | Unknown | 1 | ITSAP89 | Isere 2 Breeder | Isere, France | HiSeq |
| Wragg_et_al_2022_MER | SRR15173619 | SAMN20257245 | *A. mellifera* | Unknown | 1 | ITSAP8B | Isere 2 Breeder | Isere, France | HiSeq |
| Wragg_et_al_2022_MER | SRR15173615 | SAMN20257249 | *A. mellifera* | Unknown | 1 | ITSAP92 | Isere 2 Breeder | Isere, France | HiSeq |
| Wragg_et_al_2022_MER | SRR15173614 | SAMN20257250 | *A. mellifera* | Unknown | 1 | ITSAP93 | Isere 2 Breeder | Isere, France | HiSeq |
| Wragg_et_al_2022_MER | SRR15173732 | SAMN20257143 | *A. mellifera* | *A. m. ligustica* | 1 | ITA10A | Ligustica Italy | Italy | HiSeq |
| Wragg_et_al_2022_MER | SRR6236948 | SAMN07956273 | *A. mellifera* | *A. m. ligustica* | 1 | ITA11A | Ligustica Italy | Italy | HiSeq |
| Wragg_et_al_2022_MER | SRR6236947 | SAMN07956274 | *A. mellifera* | *A. m. ligustica* | 1 | ITA12A | Ligustica Italy | Italy | HiSeq |
| Wragg_et_al_2022_MER | SRR15173731 | SAMN20257144 | *A. mellifera* | *A. m. ligustica* | 1 | ITA13A | Ligustica Italy | Italy | HiSeq |
| Wragg_et_al_2022_MER | SRR15173730 | SAMN20257145 | *A. mellifera* | *A. m. ligustica* | 1 | ITA14A | Ligustica Italy | Italy | HiSeq |
| Wragg_et_al_2022_MER | SRR15173729 | SAMN20257146 | *A. mellifera* | *A. m. ligustica* | 1 | ITA15A | Ligustica Italy | Italy | HiSeq |
| Wragg_et_al_2022_MER | SRR15173728 | SAMN20257147 | *A. mellifera* | *A. m. ligustica* | 1 | ITA16A | Ligustica Italy | Italy | HiSeq |
| Wragg_et_al_2022_MER | SRR15173727 | SAMN20257148 | *A. mellifera* | *A. m. ligustica* | 1 | ITA17A | Ligustica Italy | Italy | HiSeq |
| Wragg_et_al_2022_MER | SRR6236950 | SAMN07956275 | *A. mellifera* | *A. m. ligustica* | 1 | ITA18A | Ligustica Italy | Italy | HiSeq |
| Wragg_et_al_2022_MER | SRR15173726 | SAMN20257149 | *A. mellifera* | *A. m. ligustica* | 1 | ITA19A | Ligustica Italy | Italy | HiSeq |
| Wragg_et_al_2022_MER | SRR6236949 | SAMN07956276 | *A. mellifera* | *A. m. ligustica* | 1 | ITA1A | Ligustica Italy | Italy | HiSeq |
| Wragg_et_al_2022_MER | SRR15173725 | SAMN20257150 | *A. mellifera* | *A. m. ligustica* | 1 | ITA20A | Ligustica Italy | Italy | HiSeq |
| Wragg_et_al_2022_MER | SRR15173724 | SAMN20257151 | *A. mellifera* | *A. m. ligustica* | 1 | ITA21A | Ligustica Italy | Italy | HiSeq |
| Wragg_et_al_2022_MER | SRR15173723 | SAMN20257152 | *A. mellifera* | *A. m. ligustica* | 1 | ITA22A | Ligustica Italy | Italy | HiSeq |
| Wragg_et_al_2022_MER | SRR15173721 | SAMN20257153 | *A. mellifera* | *A. m. ligustica* | 1 | ITA23A | Ligustica Italy | Italy | HiSeq |
| Wragg_et_al_2022_MER | SRR6236952 | SAMN07956277 | *A. mellifera* | *A. m. ligustica* | 1 | ITA24A | Ligustica Italy | Italy | HiSeq |
| Wragg_et_al_2022_MER | SRR6236951 | SAMN07956278 | *A. mellifera* | *A. m. ligustica* | 1 | ITA25A | Ligustica Italy | Italy | HiSeq |
| Wragg_et_al_2022_MER | SRR6236954 | SAMN07956279 | *A. mellifera* | *A. m. ligustica* | 1 | ITA26A | Ligustica Italy | Italy | HiSeq |
| Wragg_et_al_2022_MER | SRR15173720 | SAMN20257154 | *A. mellifera* | *A. m. ligustica* | 1 | ITA27A | Ligustica Italy | Italy | HiSeq |
| Wragg_et_al_2022_MER | SRR6236953 | SAMN07956280 | *A. mellifera* | *A. m. ligustica* | 1 | ITA28A | Ligustica Italy | Italy | HiSeq |
| Wragg_et_al_2022_MER | SRR15173719 | SAMN20257155 | *A. mellifera* | *A. m. ligustica* | 1 | ITA29A | Ligustica Italy | Italy | HiSeq |
| Wragg_et_al_2022_MER | SRR15173718 | SAMN20257156 | *A. mellifera* | *A. m. ligustica* | 1 | ITA2A | Ligustica Italy | Italy | HiSeq |
| Wragg_et_al_2022_MER | SRR6236907 | SAMN07956281 | *A. mellifera* | *A. m. ligustica* | 1 | ITA30A | Ligustica Italy | Italy | HiSeq |
| Wragg_et_al_2022_MER | SRR15173717 | SAMN20257157 | *A. mellifera* | *A. m. ligustica* | 1 | ITA3A | Ligustica Italy | Italy | HiSeq |
| Wragg_et_al_2022_MER | SRR15173716 | SAMN20257158 | *A. mellifera* | *A. m. ligustica* | 1 | ITA4A | Ligustica Italy | Italy | HiSeq |
| Wragg_et_al_2022_MER | SRR15173715 | SAMN20257159 | *A. mellifera* | *A. m. ligustica* | 1 | ITA5A | Ligustica Italy | Italy | HiSeq |
| Wragg_et_al_2022_MER | SRR6236908 | SAMN07956282 | *A. mellifera* | *A. m. ligustica* | 1 | ITA6A | Ligustica Italy | Italy | HiSeq |
| Wragg_et_al_2022_MER | SRR15173714 | SAMN20257160 | *A. mellifera* | *A. m. ligustica* | 1 | ITA7A | Ligustica Italy | Italy | HiSeq |
| Wragg_et_al_2022_MER | SRR15173713 | SAMN20257161 | *A. mellifera* | *A. m. ligustica* | 1 | ITA8A | Ligustica Italy | Italy | HiSeq |
| Wragg_et_al_2022_MER | SRR15173712 | SAMN20257162 | *A. mellifera* | *A. m. ligustica* | 1 | ITA9A | Ligustica Italy | Italy | HiSeq |
| Wragg_et_al_2022_MER | SRR15173860 | SAMN20256973 | *A. mellifera* | *A.m.mellifera* | 1 | Ab-PacBio | Ouessant Conservatory | Ouessant, France | NovaSeq |
| Wragg_et_al_2022_MER | SRR3157185 | SAMN04481280 | *A. mellifera* | *A.m.mellifera* | 1 | OUE1 | Ouessant Conservatory | Ouessant, France | HiSeq |
| Wragg_et_al_2022_MER | SRR3157175 | SAMN04481270 | *A. mellifera* | *A.m.mellifera* | 1 | OUE10 | Ouessant Conservatory | Ouessant, France | HiSeq |
| Wragg_et_al_2022_MER | SRR3157176 | SAMN04481271 | *A. mellifera* | *A.m.mellifera* | 1 | OUE11 | Ouessant Conservatory | Ouessant, France | HiSeq |
| Wragg_et_al_2022_MER | SRR3157177 | SAMN04481272 | *A. mellifera* | *A.m.mellifera* | 1 | OUE12 | Ouessant Conservatory | Ouessant, France | HiSeq |
| Wragg_et_al_2022_MER | SRR3157178 | SAMN04481273 | *A. mellifera* | *A.m.mellifera* | 1 | OUE13 | Ouessant Conservatory | Ouessant, France | HiSeq |
| Wragg_et_al_2022_MER | SRR3157179 | SAMN04481274 | *A. mellifera* | *A.m.mellifera* | 1 | OUE14 | Ouessant | Ouessant, France | HiSeq |

| | | | | | | | | | |
|---|---|---|---|---|---|---|---|---|---|
| | | | | | | | Conservatory | | |
| Wragg_et_al_2022_MER | SRR3157180 | SAMN04481275 | *A. mellifera* | *A.m.mellifera* | 1 | OUE15 | Ouessant Conservatory | Ouessant, France | HiSeq |
| Wragg_et_al_2022_MER | SRR3157181 | SAMN04481276 | *A. mellifera* | *A.m.mellifera* | 1 | OUE16 | Ouessant Conservatory | Ouessant, France | HiSeq |
| Wragg_et_al_2022_MER | SRR3157182 | SAMN04481277 | *A. mellifera* | *A.m.mellifera* | 1 | OUE17 | Ouessant Conservatory | Ouessant, France | HiSeq |
| Wragg_et_al_2022_MER | SRR3157183 | SAMN04481278 | *A. mellifera* | *A.m.mellifera* | 1 | OUE18 | Ouessant Conservatory | Ouessant, France | HiSeq |
| Wragg_et_al_2022_MER | SRR3157184 | SAMN04481279 | *A. mellifera* | *A.m.mellifera* | 1 | OUE19 | Ouessant Conservatory | Ouessant, France | HiSeq |
| Wragg_et_al_2022_MER | SRR3157196 | SAMN04481291 | *A. mellifera* | *A.m.mellifera* | 1 | OUE2 | Ouessant Conservatory | Ouessant, France | HiSeq |
| Wragg_et_al_2022_MER | SRR3157186 | SAMN04481281 | *A. mellifera* | *A.m.mellifera* | 1 | OUE20 | Ouessant Conservatory | Ouessant, France | HiSeq |
| Wragg_et_al_2022_MER | SRR3157187 | SAMN04481282 | *A. mellifera* | *A.m.mellifera* | 1 | OUE21 | Ouessant Conservatory | Ouessant, France | HiSeq |
| Wragg_et_al_2022_MER | SRR3157188 | SAMN04481283 | *A. mellifera* | *A.m.mellifera* | 1 | OUE22 | Ouessant Conservatory | Ouessant, France | HiSeq |
| Wragg_et_al_2022_MER | SRR3157189 | SAMN04481284 | *A. mellifera* | *A.m.mellifera* | 1 | OUE23 | Ouessant Conservatory | Ouessant, France | HiSeq |
| Wragg_et_al_2022_MER | SRR3157190 | SAMN04481285 | *A. mellifera* | *A.m.mellifera* | 1 | OUE24 | Ouessant Conservatory | Ouessant, France | HiSeq |
| Wragg_et_al_2022_MER | SRR3157191 | SAMN04481286 | *A. mellifera* | *A.m.mellifera* | 1 | OUE25 | Ouessant Conservatory | Ouessant, France | HiSeq |
| Wragg_et_al_2022_MER | SRR3157192 | SAMN04481287 | *A. mellifera* | *A.m.mellifera* | 1 | OUE26 | Ouessant Conservatory | Ouessant, France | HiSeq |
| Wragg_et_al_2022_MER | SRR3157193 | SAMN04481288 | *A. mellifera* | *A.m.mellifera* | 1 | OUE27 | Ouessant Conservatory | Ouessant, France | HiSeq |
| Wragg_et_al_2022_MER | SRR3157194 | SAMN04481289 | *A. mellifera* | *A.m.mellifera* | 1 | OUE28 | Ouessant Conservatory | Ouessant, France | HiSeq |
| Wragg_et_al_2022_MER | SRR3157195 | SAMN04481290 | *A. mellifera* | *A.m.mellifera* | 1 | OUE29 | Ouessant Conservatory | Ouessant, France | HiSeq |
| Wragg_et_al_2022_MER | SRR15173926 | SAMN20257304 | *A. mellifera* | *A.m.mellifera* | 1 | OUE3 | Ouessant Conservatory | Ouessant, France | HiSeq |
| Wragg_et_al_2022_MER | SRR3157197 | SAMN04481292 | *A. mellifera* | *A.m.mellifera* | 1 | OUE30 | Ouessant Conservatory | Ouessant, France | HiSeq |
| Wragg_et_al_2022_MER | SRR3157198 | SAMN04481293 | *A. mellifera* | *A.m.mellifera* | 1 | OUE31 | Ouessant Conservatory | Ouessant, France | HiSeq |
| Wragg_et_al_2022_MER | SRR3157199 | SAMN04481294 | *A. mellifera* | *A.m.mellifera* | 1 | OUE32 | Ouessant Conservatory | Ouessant, France | HiSeq |
| Wragg_et_al_2022_MER | SRR3157200 | SAMN04481295 | *A. mellifera* | *A.m.mellifera* | 1 | OUE33 | Ouessant Conservatory | Ouessant, France | HiSeq |
| Wragg_et_al_2022_MER | SRR3157201 | SAMN04481296 | *A. mellifera* | *A.m.mellifera* | 1 | OUE34 | Ouessant Conservatory | Ouessant, France | HiSeq |
| Wragg_et_al_2022_MER | SRR3157202 | SAMN04481297 | *A. mellifera* | *A.m.mellifera* | 1 | OUE35 | Ouessant Conservatory | Ouessant, France | HiSeq |
| Wragg_et_al_2022_MER | SRR3157203 | SAMN04481298 | *A. mellifera* | *A.m.mellifera* | 1 | OUE36 | Ouessant Conservatory | Ouessant, France | HiSeq |
| Wragg_et_al_2022_MER | SRR3157204 | SAMN04481299 | *A. mellifera* | *A.m.mellifera* | 1 | OUE37 | Ouessant Conservatory | Ouessant, France | HiSeq |
| Wragg_et_al_2022_MER | SRR15173925 | SAMN20257305 | *A. mellifera* | *A.m.mellifera* | 1 | OUE38 | Ouessant Conservatory | Ouessant, France | HiSeq |
| Wragg_et_al_2022_MER | SRR15173924 | SAMN20257306 | *A. mellifera* | *A.m.mellifera* | 1 | OUE39 | Ouessant Conservatory | Ouessant, France | HiSeq |
| Wragg_et_al_2022_MER | SRR15173923 | SAMN20257307 | *A. mellifera* | *A.m.mellifera* | 1 | OUE4 | Ouessant Conservatory | Ouessant, France | HiSeq |
| Wragg_et_al_2022_MER | SRR15173922 | SAMN20257308 | *A. mellifera* | *A.m.mellifera* | 1 | OUE40 | Ouessant Conservatory | Ouessant, France | HiSeq |
| Wragg_et_al_2022_MER | SRR15173921 | SAMN20257309 | *A. mellifera* | *A.m.mellifera* | 1 | OUE5 | Ouessant Conservatory | Ouessant, France | HiSeq |
| Wragg_et_al_2022_MER | SRR15173920 | SAMN20257310 | *A. mellifera* | *A.m.mellifera* | 1 | OUE6 | Ouessant Conservatory | Ouessant, France | HiSeq |
| Wragg_et_al_2022_MER | SRR15173919 | SAMN20257311 | *A. mellifera* | *A.m.mellifera* | 1 | OUE7 | Ouessant Conservatory | Ouessant, France | HiSeq |

| | | | | | | | | | |
|---|---|---|---|---|---|---|---|---|---|
| Wragg_et_al_2022_MER | SRR15173916 | SAMN20257313 | *A. mellifera* | *A.m.mellifera* | 1 | OUE9 | Ouessant Conservatory | Ouessant, France | HiSeq |
| Wragg_et_al_2022_MER | SRR15173577 | SAMN20257333 | *A. mellifera* | *A.m.mellifera* | 1 | POR1 | Porquerolles Conservatory | Porquerolles, France | HiSeq |
| Wragg_et_al_2022_MER | SRR15173576 | SAMN20257334 | *A. mellifera* | *A.m.mellifera* | 1 | POR10 | Porquerolles Conservatory | Porquerolles, France | HiSeq |
| Wragg_et_al_2022_MER | SRR15173575 | SAMN20257335 | *A. mellifera* | *A.m.mellifera* | 1 | POR11 | Porquerolles Conservatory | Porquerolles, France | HiSeq |
| Wragg_et_al_2022_MER | SRR15173574 | SAMN20257336 | *A. mellifera* | *A.m.mellifera* | 1 | POR12 | Porquerolles Conservatory | Porquerolles, France | HiSeq |
| Wragg_et_al_2022_MER | SRR15173573 | SAMN20257337 | *A. mellifera* | *A.m.mellifera* | 1 | POR13 | Porquerolles Conservatory | Porquerolles, France | HiSeq |
| Wragg_et_al_2022_MER | SRR15173572 | SAMN20257338 | *A. mellifera* | *A.m.mellifera* | 1 | POR14 | Porquerolles Conservatory | Porquerolles, France | HiSeq |
| Wragg_et_al_2022_MER | SRR6236956 | SAMN07956291 | *A. mellifera* | *A.m.mellifera* | 1 | POR15 | Porquerolles Conservatory | Porquerolles, France | HiSeq |
| Wragg_et_al_2022_MER | SRR15173571 | SAMN20257339 | *A. mellifera* | *A.m.mellifera* | 1 | POR2 | Porquerolles Conservatory | Porquerolles, France | HiSeq |
| Wragg_et_al_2022_MER | SRR15173570 | SAMN20257340 | *A. mellifera* | *A.m.mellifera* | 1 | POR3 | Porquerolles Conservatory | Porquerolles, France | HiSeq |
| Wragg_et_al_2022_MER | SRR15173569 | SAMN20257341 | *A. mellifera* | *A.m.mellifera* | 1 | POR4 | Porquerolles Conservatory | Porquerolles, France | HiSeq |
| Wragg_et_al_2022_MER | SRR15173568 | SAMN20257342 | *A. mellifera* | *A.m.mellifera* | 1 | POR5 | Porquerolles Conservatory | Porquerolles, France | HiSeq |
| Wragg_et_al_2022_MER | SRR15173566 | SAMN20257343 | *A. mellifera* | *A.m.mellifera* | 1 | POR6 | Porquerolles Conservatory | Porquerolles, France | HiSeq |
| Wragg_et_al_2022_MER | SRR6236955 | SAMN07956292 | *A. mellifera* | *A.m.mellifera* | 1 | POR7 | Porquerolles Conservatory | Porquerolles, France | HiSeq |
| Wragg_et_al_2022_MER | SRR15173565 | SAMN20257344 | *A. mellifera* | *A.m.mellifera* | 1 | POR8 | Porquerolles Conservatory | Porquerolles, France | HiSeq |
| Wragg_et_al_2022_MER | SRR15173564 | SAMN20257345 | *A. mellifera* | *A.m.mellifera* | 1 | POR9 | Porquerolles Conservatory | Porquerolles, France | HiSeq |
| Wragg_et_al_2022_MER | SRR3157089 | SAMN04481184 | *A. mellifera* | Unknown | 1 | FL1 | Royal Jelly France | France | HiSeq |
| Wragg_et_al_2022_MER | SRR15173743 | SAMN20257133 | *A. mellifera* | Unknown | 1 | FL106 | Royal Jelly France | France | HiSeq |
| Wragg_et_al_2022_MER | SRR15173742 | SAMN20257134 | *A. mellifera* | Unknown | 1 | FL109 | Royal Jelly France | France | HiSeq |
| Wragg_et_al_2022_MER | SRR3157085 | SAMN04481180 | *A. mellifera* | Unknown | 1 | FL11 | Royal Jelly France | France | HiSeq |
| Wragg_et_al_2022_MER | SRR3157086 | SAMN04481181 | *A. mellifera* | Unknown | 1 | FL12 | Royal Jelly France | France | HiSeq |
| Wragg_et_al_2022_MER | SRR3157087 | SAMN04481182 | *A. mellifera* | Unknown | 1 | FL13 | Royal Jelly France | France | HiSeq |
| Wragg_et_al_2022_MER | SRR3157088 | SAMN04481183 | *A. mellifera* | Unknown | 1 | FL15 | Royal Jelly France | France | HiSeq |
| Wragg_et_al_2022_MER | SRR15173741 | SAMN20257135 | *A. mellifera* | Unknown | 1 | FL17 | Royal Jelly France | France | HiSeq |
| Wragg_et_al_2022_MER | SRR15173740 | SAMN20257136 | *A. mellifera* | Unknown | 1 | FL1bis | Royal Jelly France | France | HiSeq |
| Wragg_et_al_2022_MER | SRR3157090 | SAMN04481185 | *A. mellifera* | Unknown | 1 | FL2 | Royal Jelly France | France | HiSeq |
| Wragg_et_al_2022_MER | SRR15173739 | SAMN20257137 | *A. mellifera* | Unknown | 1 | FL22 | Royal Jelly France | France | HiSeq |
| Wragg_et_al_2022_MER | SRR3157091 | SAMN04481186 | *A. mellifera* | Unknown | 1 | FL3 | Royal Jelly France | France | HiSeq |
| Wragg_et_al_2022_MER | SRR15173738 | SAMN20257138 | *A. mellifera* | Unknown | 1 | FL3bis | Royal Jelly France | France | HiSeq |
| Wragg_et_al_2022_MER | SRR15173737 | SAMN20257139 | *A. mellifera* | Unknown | 1 | FL61 | Royal Jelly France | France | HiSeq |
| Wragg_et_al_2022_MER | SRR3157092 | SAMN04481187 | *A. mellifera* | Unknown | 1 | FL7 | Royal Jelly France | France | HiSeq |
| Wragg_et_al_2022_MER | SRR3157093 | SAMN04481188 | *A. mellifera* | Unknown | 1 | FL8 | Royal Jelly France | France | HiSeq |
| Wragg_et_al_2022_MER | SRR15173736 | SAMN20257140 | *A. mellifera* | Unknown | 1 | FL86 | Royal Jelly France | France | HiSeq |
| Wragg_et_al_2022_MER | SRR15173735 | SAMN20257141 | *A. mellifera* | Unknown | 1 | FL87 | Royal Jelly France | France | HiSeq |
| Wragg_et_al_2022_MER | SRR3157094 | SAMN04481189 | *A. mellifera* | Unknown | 1 | FL9 | Royal Jelly France | France | HiSeq |
| Wragg_et_al_2022_MER | SRR15173734 | SAMN20257142 | *A. mellifera* | Unknown | 1 | FL9bis | Royal Jelly France | France | HiSeq |
| Wragg_et_al_2022_MER | SRR15173701 | SAMN20257172 | *A. mellifera* | Unknown | 1 | ITSAP11 | Royal Jelly France | France | HiSeq |
| Wragg_et_al_2022_MER | SRR15173686 | SAMN20257184 | *A. mellifera* | Unknown | 1 | ITSAP20 | Royal Jelly France | France | HiSeq |

| | | | | | | | | | |
|---|---|---|---|---|---|---|---|---|---|
| Wragg_et_al_2022_MER | SRR15173683 | SAMN20257187 | *A. mellifera* | Unknown | 1 | ITSAP24 | Royal Jelly France | France | HiSeq |
| Wragg_et_al_2022_MER | SRR15173679 | SAMN20257191 | *A. mellifera* | Unknown | 1 | ITSAP2B | Royal Jelly France | France | HiSeq |
| Wragg_et_al_2022_MER | SRR15173678 | SAMN20257192 | *A. mellifera* | Unknown | 1 | ITSAP3 | Royal Jelly France | France | HiSeq |
| Wragg_et_al_2022_MER | SRR15173675 | SAMN20257194 | *A. mellifera* | Unknown | 1 | ITSAP31 | Royal Jelly France | France | HiSeq |
| Wragg_et_al_2022_MER | SRR15173663 | SAMN20257205 | *A. mellifera* | Unknown | 1 | ITSAP46 | Royal Jelly France | France | HiSeq |
| Wragg_et_al_2022_MER | SRR15173658 | SAMN20257210 | *A. mellifera* | Unknown | 1 | ITSAP50 | Royal Jelly France | France | HiSeq |
| Wragg_et_al_2022_MER | SRR15173653 | SAMN20257214 | *A. mellifera* | Unknown | 1 | ITSAP54 | Royal Jelly France | France | HiSeq |
| Wragg_et_al_2022_MER | SRR15173648 | SAMN20257219 | *A. mellifera* | Unknown | 1 | ITSAP62 | Royal Jelly France | France | HiSeq |
| Wragg_et_al_2022_MER | SRR15173645 | SAMN20257222 | *A. mellifera* | Unknown | 1 | ITSAP65 | Royal Jelly France | France | HiSeq |
| Wragg_et_al_2022_MER | SRR15173643 | SAMN20257223 | *A. mellifera* | Unknown | 1 | ITSAP66 | Royal Jelly France | France | HiSeq |
| Wragg_et_al_2022_MER | SRR15173630 | SAMN20257235 | *A. mellifera* | Unknown | 1 | ITSAP79 | Royal Jelly France | France | HiSeq |
| Wragg_et_al_2022_MER | SRR15173628 | SAMN20257237 | *A. mellifera* | Unknown | 1 | ITSAP81 | Royal Jelly France | France | HiSeq |
| Wragg_et_al_2022_MER | SRR15173617 | SAMN20257247 | *A. mellifera* | Unknown | 1 | ITSAP90 | Royal Jelly France | France | HiSeq |
| Wragg_et_al_2022_MER | SRR3157095 | SAMN04481190 | *A. mellifera* | Unknown | 1 | NM1 | Royal Jelly France | France | HiSeq |
| Wragg_et_al_2022_MER | SRR3157101 | SAMN04481196 | *A. mellifera* | Unknown | 1 | NM2 | Royal Jelly France | France | HiSeq |
| Wragg_et_al_2022_MER | SRR3157096 | SAMN04481191 | *A. mellifera* | Unknown | 1 | NM21 | Royal Jelly France | France | HiSeq |
| Wragg_et_al_2022_MER | SRR3157097 | SAMN04481192 | *A. mellifera* | Unknown | 1 | NM25 | Royal Jelly France | France | HiSeq |
| Wragg_et_al_2022_MER | SRR3157098 | SAMN04481193 | *A. mellifera* | Unknown | 1 | NM26 | Royal Jelly France | France | HiSeq |
| Wragg_et_al_2022_MER | SRR3157099 | SAMN04481194 | *A. mellifera* | Unknown | 1 | NM28 | Royal Jelly France | France | HiSeq |
| Wragg_et_al_2022_MER | SRR3157100 | SAMN04481195 | *A. mellifera* | Unknown | 1 | NM29 | Royal Jelly France | France | HiSeq |
| Wragg_et_al_2022_MER | SRR3157102 | SAMN04481197 | *A. mellifera* | Unknown | 1 | NM3 | Royal Jelly France | France | HiSeq |
| Wragg_et_al_2022_MER | SRR3157103 | SAMN04481198 | *A. mellifera* | Unknown | 1 | NM4 | Royal Jelly France | France | HiSeq |
| Wragg_et_al_2022_MER | SRR3157104 | SAMN04481199 | *A. mellifera* | Unknown | 1 | NM5 | Royal Jelly France | France | HiSeq |
| Wragg_et_al_2022_MER | SRR3157105 | SAMN04481200 | *A. mellifera* | Unknown | 1 | NM6 | Royal Jelly France | France | HiSeq |
| Wragg_et_al_2022_MER | SRR3157106 | SAMN04481201 | *A. mellifera* | Unknown | 1 | NM7 | Royal Jelly France | France | HiSeq |
| Wragg_et_al_2022_MER | SRR3157107 | SAMN04481202 | *A. mellifera* | Unknown | 1 | NM8 | Royal Jelly France | France | HiSeq |
| Wragg_et_al_2022_MER | SRR3157108 | SAMN04481203 | *A. mellifera* | Unknown | 1 | NM9 | Royal Jelly France | France | HiSeq |
| Wragg_et_al_2022_MER | SRR3157110 | SAMN04481205 | *A. mellifera* | Unknown | 1 | XC1 | Royal Jelly France | France | HiSeq |
| Wragg_et_al_2022_MER | SRR3157113 | SAMN04481208 | *A. mellifera* | Unknown | 1 | XC5 | Royal Jelly France | France | HiSeq |
| Wragg_et_al_2022_MER | SRR3157114 | SAMN04481209 | *A. mellifera* | Unknown | 1 | XC9 | Royal Jelly France | France | HiSeq |
| Wragg_et_al_2022_MER | SRR15173494 | SAMN20257408 | *A. mellifera* | Unknown | 1 | YC1 | Royal Jelly France | France | HiSeq |
| Wragg_et_al_2022_MER | SRR15173493 | SAMN20257409 | *A. mellifera* | Unknown | 1 | YC10 | Royal Jelly France | France | HiSeq |
| Wragg_et_al_2022_MER | SRR15173492 | SAMN20257410 | *A. mellifera* | Unknown | 1 | YC2 | Royal Jelly France | France | HiSeq |
| Wragg_et_al_2022_MER | SRR15173491 | SAMN20257411 | *A. mellifera* | Unknown | 1 | YC3 | Royal Jelly France | France | HiSeq |
| Wragg_et_al_2022_MER | SRR15173490 | SAMN20257412 | *A. mellifera* | Unknown | 1 | YC4 | Royal Jelly France | France | HiSeq |
| Wragg_et_al_2022_MER | SRR15173488 | SAMN20257413 | *A. mellifera* | Unknown | 1 | YC5 | Royal Jelly France | France | HiSeq |
| Wragg_et_al_2022_MER | SRR15173487 | SAMN20257414 | *A. mellifera* | Unknown | 1 | YC6 | Royal Jelly France | France | HiSeq |
| Wragg_et_al_2022_MER | SRR15173486 | SAMN20257415 | *A. mellifera* | Unknown | 1 | YC7 | Royal Jelly France | France | HiSeq |
| Wragg_et_al_2022_MER | SRR15173485 | SAMN20257416 | *A. mellifera* | Unknown | 1 | YC8 | Royal Jelly France | France | HiSeq |
| Wragg_et_al_2022_MER | SRR15173484 | SAMN20257417 | *A. mellifera* | Unknown | 1 | YC9 | Royal Jelly France | France | HiSeq |
| Wragg_et_al_2022_MER | SRR15173563 | SAMN20257346 | *A. mellifera* | Unknown | 1 | Sar1 | Sarthe Breeder | Sarthe, France | HiSeq |
| Wragg_et_al_2022_MER | SRR15173562 | SAMN20257347 | *A. mellifera* | Unknown | 1 | Sar11 | Sarthe Breeder | Sarthe, France | HiSeq |
| Wragg_et_al_2022_MER | SRR15173561 | SAMN20257348 | *A. mellifera* | Unknown | 1 | Sar13 | Sarthe Breeder | Sarthe, France | HiSeq |

| | | | | | | | | | |
|---|---|---|---|---|---|---|---|---|---|
| Wragg_et_al_2022_MER | SRR15173560 | SAMN20257349 | *A. mellifera* | Unknown | 1 | Sar15 | Sarthe Breeder | Sarthe, France | HiSeq |
| Wragg_et_al_2022_MER | SRR15173559 | SAMN20257350 | *A. mellifera* | Unknown | 1 | Sar17 | Sarthe Breeder | Sarthe, France | HiSeq |
| Wragg_et_al_2022_MER | SRR15173558 | SAMN20257351 | *A. mellifera* | Unknown | 1 | Sar19 | Sarthe Breeder | Sarthe, France | HiSeq |
| Wragg_et_al_2022_MER | SRR15173557 | SAMN20257352 | *A. mellifera* | Unknown | 1 | Sar21 | Sarthe Breeder | Sarthe, France | HiSeq |
| Wragg_et_al_2022_MER | SRR15173555 | SAMN20257353 | *A. mellifera* | Unknown | 1 | Sar23 | Sarthe Breeder | Sarthe, France | HiSeq |
| Wragg_et_al_2022_MER | SRR15173554 | SAMN20257354 | *A. mellifera* | Unknown | 1 | Sar25 | Sarthe Breeder | Sarthe, France | HiSeq |
| Wragg_et_al_2022_MER | SRR15173553 | SAMN20257355 | *A. mellifera* | Unknown | 1 | Sar27 | Sarthe Breeder | Sarthe, France | HiSeq |
| Wragg_et_al_2022_MER | SRR15173552 | SAMN20257356 | *A. mellifera* | Unknown | 1 | Sar29 | Sarthe Breeder | Sarthe, France | HiSeq |
| Wragg_et_al_2022_MER | SRR15173551 | SAMN20257357 | *A. mellifera* | Unknown | 1 | Sar3 | Sarthe Breeder | Sarthe, France | HiSeq |
| Wragg_et_al_2022_MER | SRR15173550 | SAMN20257358 | *A. mellifera* | Unknown | 1 | Sar31 | Sarthe Breeder | Sarthe, France | HiSeq |
| Wragg_et_al_2022_MER | SRR15173549 | SAMN20257359 | *A. mellifera* | Unknown | 1 | Sar33 | Sarthe Breeder | Sarthe, France | HiSeq |
| Wragg_et_al_2022_MER | SRR15173548 | SAMN20257360 | *A. mellifera* | Unknown | 1 | Sar5 | Sarthe Breeder | Sarthe, France | HiSeq |
| Wragg_et_al_2022_MER | SRR15173547 | SAMN20257361 | *A. mellifera* | Unknown | 1 | Sar7 | Sarthe Breeder | Sarthe, France | HiSeq |
| Wragg_et_al_2022_MER | SRR15173546 | SAMN20257362 | *A. mellifera* | Unknown | 1 | Sar9 | Sarthe Breeder | Sarthe, France | HiSeq |
| Wragg_et_al_2022_MER | SRR5021018 | SAMN06017588 | *A. mellifera* | *A.m.mellifera* | 1 | SavA10 | Savoy Conservatory | Savoy, France | HiSeq |
| Wragg_et_al_2022_MER | SRR5021009 | SAMN06017590 | *A. mellifera* | *A.m.mellifera* | 1 | SavA11 | Savoy Conservatory | Savoy, France | HiSeq |
| Wragg_et_al_2022_MER | SRR5021027 | SAMN06017591 | *A. mellifera* | *A.m.mellifera* | 1 | SavA12 | Savoy Conservatory | Savoy, France | HiSeq |
| Wragg_et_al_2022_MER | SRR5021036 | SAMN06017593 | *A. mellifera* | *A.m.mellifera* | 1 | SavA14 | Savoy Conservatory | Savoy, France | HiSeq |
| Wragg_et_al_2022_MER | SRR5021038 | SAMN06017595 | *A. mellifera* | *A.m.mellifera* | 1 | SavA15 | Savoy Conservatory | Savoy, France | HiSeq |
| Wragg_et_al_2022_MER | SRR5021023 | SAMN06017597 | *A. mellifera* | *A.m.mellifera* | 1 | SavA16 | Savoy Conservatory | Savoy, France | HiSeq |
| Wragg_et_al_2022_MER | SRR5021031 | SAMN06017599 | *A. mellifera* | *A.m.mellifera* | 1 | SavA17 | Savoy Conservatory | Savoy, France | HiSeq |
| Wragg_et_al_2022_MER | SRR5021013 | SAMN06017601 | *A. mellifera* | *A.m.mellifera* | 1 | SavA18 | Savoy Conservatory | Savoy, France | HiSeq |
| Wragg_et_al_2022_MER | SRR5021032 | SAMN06017579 | *A. mellifera* | *A.m.mellifera* | 1 | SavA2 | Savoy Conservatory | Savoy, France | HiSeq |
| Wragg_et_al_2022_MER | SRR5021017 | SAMN06017581 | *A. mellifera* | *A.m.mellifera* | 1 | SavA4 | Savoy Conservatory | Savoy, France | HiSeq |
| Wragg_et_al_2022_MER | SRR5021037 | SAMN06017583 | *A. mellifera* | *A.m.mellifera* | 1 | SavA7 | Savoy Conservatory | Savoy, France | HiSeq |
| Wragg_et_al_2022_MER | SRR5021019 | SAMN06017585 | *A. mellifera* | *A.m.mellifera* | 1 | SavA8 | Savoy Conservatory | Savoy, France | HiSeq |
| Wragg_et_al_2022_MER | SRR5021015 | SAMN06017586 | *A. mellifera* | *A.m.mellifera* | 1 | SavA9 | Savoy Conservatory | Savoy, France | HiSeq |
| Wragg_et_al_2022_MER | SRR5021029 | SAMN06017589 | *A. mellifera* | *A.m.mellifera* | 1 | SavB10 | Savoy Conservatory | Savoy, France | HiSeq |
| Wragg_et_al_2022_MER | SRR5021008 | SAMN06017592 | *A. mellifera* | *A.m.mellifera* | 1 | SavB13 | Savoy Conservatory | Savoy, France | HiSeq |
| Wragg_et_al_2022_MER | SRR5021016 | SAMN06017594 | *A. mellifera* | *A.m.mellifera* | 1 | SavB14 | Savoy Conservatory | Savoy, France | HiSeq |
| Wragg_et_al_2022_MER | SRR5021035 | SAMN06017596 | *A. mellifera* | *A.m.mellifera* | 1 | SavB15 | Savoy Conservatory | Savoy, France | HiSeq |
| Wragg_et_al_2022_MER | SRR5021024 | SAMN06017598 | *A. mellifera* | *A.m.mellifera* | 1 | SavB16 | Savoy Conservatory | Savoy, France | HiSeq |
| Wragg_et_al_2022_MER | SRR5021028 | SAMN06017600 | *A. mellifera* | *A.m.mellifera* | 1 | SavB17 | Savoy Conservatory | Savoy, France | HiSeq |
| Wragg_et_al_2022_MER | SRR5021025 | SAMN06017602 | *A. mellifera* | *A.m.mellifera* | 1 | SavB18 | Savoy Conservatory | Savoy, France | HiSeq |
| Wragg_et_al_2022_MER | SRR5021022 | SAMN06017603 | *A. mellifera* | *A.m.mellifera* | 1 | SavB20 | Savoy Conservatory | Savoy, France | HiSeq |
| Wragg_et_al_2022_MER | SRR5021026 | SAMN06017604 | *A. mellifera* | *A.m.mellifera* | 1 | SavB21 | Savoy Conservatory | Savoy, France | HiSeq |
| Wragg_et_al_2022_MER | SRR5021030 | SAMN06017605 | *A. mellifera* | *A.m.mellifera* | 1 | SavB23 | Savoy Conservatory | Savoy, France | HiSeq |
| Wragg_et_al_2022_MER | SRR5021034 | SAMN06017582 | *A. mellifera* | *A.m.mellifera* | 1 | SavB5 | Savoy Conservatory | Savoy, France | HiSeq |
| Wragg_et_al_2022_MER | SRR5021020 | SAMN06017584 | *A. mellifera* | *A.m.mellifera* | 1 | SavB7 | Savoy Conservatory | Savoy, France | HiSeq |
| Wragg_et_al_2022_MER | SRR5021010 | SAMN06017587 | *A. mellifera* | *A.m.mellifera* | 1 | SavB9 | Savoy Conservatory | Savoy, France | HiSeq |
| Wragg_et_al_2022_MER | SRR5021021 | SAMN06017606 | *A. mellifera* | *A.m.mellifera* | 1 | SavC34 | Savoy Conservatory | Savoy, France | HiSeq |
| Wragg_et_al_2022_MER | SRR5021033 | SAMN06017607 | *A. mellifera* | *A.m.mellifera* | 1 | SavC36 | Savoy Conservatory | Savoy, France | HiSeq |
| Wragg_et_al_2022_MER | SRR5021012 | SAMN06017608 | *A. mellifera* | *A.m.mellifera* | 1 | SavC40 | Savoy Conservatory | Savoy, France | HiSeq |

| | | | | | | | | | |
|---|---|---|---|---|---|---|---|---|---|
| Wragg_et_al_2022_MER | SRR6236970 | SAMN07956303 | *A. mellifera* | *A.m.mellifera* | 1 | SOL1 | Sollies Conservatory | Sollies, France | HiSeq |
| Wragg_et_al_2022_MER | SRR15173542 | SAMN20257365 | *A. mellifera* | *A.m.mellifera* | 1 | SOL10 | Sollies Conservatory | Sollies, France | HiSeq |
| Wragg_et_al_2022_MER | SRR15173541 | SAMN20257366 | *A. mellifera* | *A.m.mellifera* | 1 | SOL11 | Sollies Conservatory | Sollies, France | HiSeq |
| Wragg_et_al_2022_MER | SRR15173540 | SAMN20257367 | *A. mellifera* | *A.m.mellifera* | 1 | SOL12 | Sollies Conservatory | Sollies, France | HiSeq |
| Wragg_et_al_2022_MER | SRR15173539 | SAMN20257368 | *A. mellifera* | *A.m.mellifera* | 1 | SOL13 | Sollies Conservatory | Sollies, France | HiSeq |
| Wragg_et_al_2022_MER | SRR15173538 | SAMN20257369 | *A. mellifera* | *A.m.mellifera* | 1 | SOL14 | Sollies Conservatory | Sollies, France | HiSeq |
| Wragg_et_al_2022_MER | SRR15173537 | SAMN20257370 | *A. mellifera* | *A.m.mellifera* | 1 | SOL2 | Sollies Conservatory | Sollies, France | HiSeq |
| Wragg_et_al_2022_MER | SRR15173536 | SAMN20257371 | *A. mellifera* | *A.m.mellifera* | 1 | SOL3 | Sollies Conservatory | Sollies, France | HiSeq |
| Wragg_et_al_2022_MER | SRR6236961 | SAMN07956304 | *A. mellifera* | *A.m.mellifera* | 1 | SOL4 | Sollies Conservatory | Sollies, France | HiSeq |
| Wragg_et_al_2022_MER | SRR15173535 | SAMN20257372 | *A. mellifera* | *A.m.mellifera* | 1 | SOL5 | Sollies Conservatory | Sollies, France | HiSeq |
| Wragg_et_al_2022_MER | SRR15173532 | SAMN20257373 | *A. mellifera* | *A.m.mellifera* | 1 | SOL6 | Sollies Conservatory | Sollies, France | HiSeq |
| Wragg_et_al_2022_MER | SRR15173531 | SAMN20257374 | *A. mellifera* | *A.m.mellifera* | 1 | SOL7 | Sollies Conservatory | Sollies, France | HiSeq |
| Wragg_et_al_2022_MER | SRR15173530 | SAMN20257375 | *A. mellifera* | *A.m.mellifera* | 1 | SOL8 | Sollies Conservatory | Sollies, France | HiSeq |
| Wragg_et_al_2022_MER | SRR15173529 | SAMN20257376 | *A. mellifera* | *A.m.mellifera* | 1 | SOL9 | Sollies Conservatory | Sollies, France | HiSeq |
| Wragg_et_al_2022_MER | SRR15173708 | SAMN20257165 | *A. mellifera* | Unknown | 1 | ITSAP100 | Tarn 1 Breeder | Tarn, France | HiSeq |
| Wragg_et_al_2022_MER | SRR15173707 | SAMN20257166 | *A. mellifera* | Unknown | 1 | ITSAP101 | Tarn 1 Breeder | Tarn, France | HiSeq |
| Wragg_et_al_2022_MER | SRR15173706 | SAMN20257167 | *A. mellifera* | Unknown | 1 | ITSAP102 | Tarn 1 Breeder | Tarn, France | HiSeq |
| Wragg_et_al_2022_MER | SRR15173705 | SAMN20257168 | *A. mellifera* | Unknown | 1 | ITSAP103 | Tarn 1 Breeder | Tarn, France | HiSeq |
| Wragg_et_al_2022_MER | SRR15173698 | SAMN20257173 | *A. mellifera* | Unknown | 1 | ITSAP12 | Tarn 1 Breeder | Tarn, France | HiSeq |
| Wragg_et_al_2022_MER | SRR15173694 | SAMN20257177 | *A. mellifera* | Unknown | 1 | ITSAP15B | Tarn 1 Breeder | Tarn, France | HiSeq |
| Wragg_et_al_2022_MER | SRR15173668 | SAMN20257201 | *A. mellifera* | Unknown | 1 | ITSAP40 | Tarn 1 Breeder | Tarn, France | HiSeq |
| Wragg_et_al_2022_MER | SRR15173664 | SAMN20257204 | *A. mellifera* | Unknown | 1 | ITSAP45 | Tarn 1 Breeder | Tarn, France | HiSeq |
| Wragg_et_al_2022_MER | SRR15173637 | SAMN20257229 | *A. mellifera* | Unknown | 1 | ITSAP70 | Tarn 1 Breeder | Tarn, France | HiSeq |
| Wragg_et_al_2022_MER | SRR15173635 | SAMN20257231 | *A. mellifera* | Unknown | 1 | ITSAP72 | Tarn 1 Breeder | Tarn, France | HiSeq |
| Wragg_et_al_2022_MER | SRR15173632 | SAMN20257233 | *A. mellifera* | Unknown | 1 | ITSAP74 | Tarn 1 Breeder | Tarn, France | HiSeq |
| Wragg_et_al_2022_MER | SRR15173631 | SAMN20257234 | *A. mellifera* | Unknown | 1 | ITSAP78 | Tarn 1 Breeder | Tarn, France | HiSeq |
| Wragg_et_al_2022_MER | SRR15173621 | SAMN20257243 | *A. mellifera* | Unknown | 1 | ITSAP88 | Tarn 1 Breeder | Tarn, France | HiSeq |
| Wragg_et_al_2022_MER | SRR15173612 | SAMN20257252 | *A. mellifera* | Unknown | 1 | ITSAP95 | Tarn 1 Breeder | Tarn, France | HiSeq |
| Wragg_et_al_2022_MER | SRR3157125 | SAMN04481220 | *A. mellifera* | Unknown | 1 | JFM1 | Tarn 1 Breeder | Tarn, France | HiSeq |
| Wragg_et_al_2022_MER | SRR3157115 | SAMN04481210 | *A. mellifera* | Unknown | 1 | JFM10 | Tarn 1 Breeder | Tarn, France | HiSeq |
| Wragg_et_al_2022_MER | SRR3157116 | SAMN04481211 | *A. mellifera* | Unknown | 1 | JFM11 | Tarn 1 Breeder | Tarn, France | HiSeq |
| Wragg_et_al_2022_MER | SRR3157117 | SAMN04481212 | *A. mellifera* | Unknown | 1 | JFM12 | Tarn 1 Breeder | Tarn, France | HiSeq |
| Wragg_et_al_2022_MER | SRR3157118 | SAMN04481213 | *A. mellifera* | Unknown | 1 | JFM13 | Tarn 1 Breeder | Tarn, France | HiSeq |
| Wragg_et_al_2022_MER | SRR3157119 | SAMN04481214 | *A. mellifera* | Unknown | 1 | JFM14 | Tarn 1 Breeder | Tarn, France | HiSeq |
| Wragg_et_al_2022_MER | SRR3157120 | SAMN04481215 | *A. mellifera* | Unknown | 1 | JFM15 | Tarn 1 Breeder | Tarn, France | HiSeq |
| Wragg_et_al_2022_MER | SRR3157121 | SAMN04481216 | *A. mellifera* | Unknown | 1 | JFM16 | Tarn 1 Breeder | Tarn, France | HiSeq |
| Wragg_et_al_2022_MER | SRR3157122 | SAMN04481217 | *A. mellifera* | Unknown | 1 | JFM17 | Tarn 1 Breeder | Tarn, France | HiSeq |
| Wragg_et_al_2022_MER | SRR3157123 | SAMN04481218 | *A. mellifera* | Unknown | 1 | JFM18 | Tarn 1 Breeder | Tarn, France | HiSeq |
| Wragg_et_al_2022_MER | SRR3157124 | SAMN04481219 | *A. mellifera* | Unknown | 1 | JFM19 | Tarn 1 Breeder | Tarn, France | HiSeq |
| Wragg_et_al_2022_MER | SRR3157135 | SAMN04481230 | *A. mellifera* | Unknown | 1 | JFM2 | Tarn 1 Breeder | Tarn, France | HiSeq |
| Wragg_et_al_2022_MER | SRR3157126 | SAMN04481221 | *A. mellifera* | Unknown | 1 | JFM20 | Tarn 1 Breeder | Tarn, France | HiSeq |
| Wragg_et_al_2022_MER | SRR3157128 | SAMN04481223 | *A. mellifera* | Unknown | 1 | JFM22 | Tarn 1 Breeder | Tarn, France | HiSeq |
| Wragg_et_al_2022_MER | SRR3157129 | SAMN04481224 | *A. mellifera* | Unknown | 1 | JFM23 | Tarn 1 Breeder | Tarn, France | HiSeq |

| | | | | | | | | | |
|---|---|---|---|---|---|---|---|---|---|
| Wragg_et_al_2022_MER | SRR3157131 | SAMN04481226 | *A. mellifera* | Unknown | 1 | JFM25 | Tarn 1 Breeder | Tarn, France | HiSeq |
| Wragg_et_al_2022_MER | SRR3157132 | SAMN04481227 | *A. mellifera* | Unknown | 1 | JFM26 | Tarn 1 Breeder | Tarn, France | HiSeq |
| Wragg_et_al_2022_MER | SRR3157133 | SAMN04481228 | *A. mellifera* | Unknown | 1 | JFM28 | Tarn 1 Breeder | Tarn, France | HiSeq |
| Wragg_et_al_2022_MER | SRR3157134 | SAMN04481229 | *A. mellifera* | Unknown | 1 | JFM29 | Tarn 1 Breeder | Tarn, France | HiSeq |
| Wragg_et_al_2022_MER | SRR3157136 | SAMN04481231 | *A. mellifera* | Unknown | 1 | JFM31 | Tarn 1 Breeder | Tarn, France | HiSeq |
| Wragg_et_al_2022_MER | SRR3157137 | SAMN04481232 | *A. mellifera* | Unknown | 1 | JFM33 | Tarn 1 Breeder | Tarn, France | HiSeq |
| Wragg_et_al_2022_MER | SRR3157139 | SAMN04481234 | *A. mellifera* | Unknown | 1 | JFM4 | Tarn 1 Breeder | Tarn, France | HiSeq |
| Wragg_et_al_2022_MER | SRR3157141 | SAMN04481236 | *A. mellifera* | Unknown | 1 | JFM6 | Tarn 1 Breeder | Tarn, France | HiSeq |
| Wragg_et_al_2022_MER | SRR3157142 | SAMN04481237 | *A. mellifera* | Unknown | 1 | JFM7 | Tarn 1 Breeder | Tarn, France | HiSeq |
| Wragg_et_al_2022_MER | SRR3157143 | SAMN04481238 | *A. mellifera* | Unknown | 1 | JFM8 | Tarn 1 Breeder | Tarn, France | HiSeq |
| Wragg_et_al_2022_MER | SRR3157144 | SAMN04481239 | *A. mellifera* | Unknown | 1 | JFM9 | Tarn 1 Breeder | Tarn, France | HiSeq |
| Wragg_et_al_2022_MER | SRR15173879 | SAMN20257273 | *A. mellifera* | Unknown | 1 | KF1 | Tarn 2 Breeder | Tarn, France | HiSeq |
| Wragg_et_al_2022_MER | SRR15173959 | SAMN20257274 | *A. mellifera* | Unknown | 1 | KF10 | Tarn 2 Breeder | Tarn, France | HiSeq |
| Wragg_et_al_2022_MER | SRR15173958 | SAMN20257275 | *A. mellifera* | Unknown | 1 | KF11 | Tarn 2 Breeder | Tarn, France | HiSeq |
| Wragg_et_al_2022_MER | SRR15173957 | SAMN20257276 | *A. mellifera* | Unknown | 1 | KF12 | Tarn 2 Breeder | Tarn, France | HiSeq |
| Wragg_et_al_2022_MER | SRR15173956 | SAMN20257277 | *A. mellifera* | Unknown | 1 | KF13 | Tarn 2 Breeder | Tarn, France | HiSeq |
| Wragg_et_al_2022_MER | SRR15173955 | SAMN20257278 | *A. mellifera* | Unknown | 1 | KF14 | Tarn 2 Breeder | Tarn, France | HiSeq |
| Wragg_et_al_2022_MER | SRR15173954 | SAMN20257279 | *A. mellifera* | Unknown | 1 | KF15 | Tarn 2 Breeder | Tarn, France | HiSeq |
| Wragg_et_al_2022_MER | SRR15173953 | SAMN20257280 | *A. mellifera* | Unknown | 1 | KF16 | Tarn 2 Breeder | Tarn, France | HiSeq |
| Wragg_et_al_2022_MER | SRR15173952 | SAMN20257281 | *A. mellifera* | Unknown | 1 | KF17 | Tarn 2 Breeder | Tarn, France | HiSeq |
| Wragg_et_al_2022_MER | SRR15173951 | SAMN20257282 | *A. mellifera* | Unknown | 1 | KF18 | Tarn 2 Breeder | Tarn, France | HiSeq |
| Wragg_et_al_2022_MER | SRR15173949 | SAMN20257283 | *A. mellifera* | Unknown | 1 | KF19 | Tarn 2 Breeder | Tarn, France | HiSeq |
| Wragg_et_al_2022_MER | SRR15173948 | SAMN20257284 | *A. mellifera* | Unknown | 1 | KF2 | Tarn 2 Breeder | Tarn, France | HiSeq |
| Wragg_et_al_2022_MER | SRR15173947 | SAMN20257285 | *A. mellifera* | Unknown | 1 | KF20 | Tarn 2 Breeder | Tarn, France | HiSeq |
| Wragg_et_al_2022_MER | SRR15173945 | SAMN20257287 | *A. mellifera* | Unknown | 1 | KF22 | Tarn 2 Breeder | Tarn, France | HiSeq |
| Wragg_et_al_2022_MER | SRR15173944 | SAMN20257288 | *A. mellifera* | Unknown | 1 | KF23 | Tarn 2 Breeder | Tarn, France | HiSeq |
| Wragg_et_al_2022_MER | SRR15173943 | SAMN20257289 | *A. mellifera* | Unknown | 1 | KF24 | Tarn 2 Breeder | Tarn, France | HiSeq |
| Wragg_et_al_2022_MER | SRR15173942 | SAMN20257290 | *A. mellifera* | Unknown | 1 | KF25 | Tarn 2 Breeder | Tarn, France | HiSeq |
| Wragg_et_al_2022_MER | SRR15173941 | SAMN20257291 | *A. mellifera* | Unknown | 1 | KF26 | Tarn 2 Breeder | Tarn, France | HiSeq |
| Wragg_et_al_2022_MER | SRR15173940 | SAMN20257292 | *A. mellifera* | Unknown | 1 | KF27 | Tarn 2 Breeder | Tarn, France | HiSeq |
| Wragg_et_al_2022_MER | SRR15173938 | SAMN20257293 | *A. mellifera* | Unknown | 1 | KF3 | Tarn 2 Breeder | Tarn, France | HiSeq |
| Wragg_et_al_2022_MER | SRR15173937 | SAMN20257294 | *A. mellifera* | Unknown | 1 | KF4 | Tarn 2 Breeder | Tarn, France | HiSeq |
| Wragg_et_al_2022_MER | SRR15173936 | SAMN20257295 | *A. mellifera* | Unknown | 1 | KF41 | Tarn 2 Breeder | Tarn, France | HiSeq |
| Wragg_et_al_2022_MER | SRR15173935 | SAMN20257296 | *A. mellifera* | Unknown | 1 | KF42 | Tarn 2 Breeder | Tarn, France | HiSeq |
| Wragg_et_al_2022_MER | SRR15173934 | SAMN20257297 | *A. mellifera* | Unknown | 1 | KF43 | Tarn 2 Breeder | Tarn, France | HiSeq |
| Wragg_et_al_2022_MER | SRR15173933 | SAMN20257298 | *A. mellifera* | Unknown | 1 | KF5 | Tarn 2 Breeder | Tarn, France | HiSeq |
| Wragg_et_al_2022_MER | SRR15173932 | SAMN20257299 | *A. mellifera* | Unknown | 1 | KF6 | Tarn 2 Breeder | Tarn, France | HiSeq |
| Wragg_et_al_2022_MER | SRR15173931 | SAMN20257300 | *A. mellifera* | Unknown | 1 | KF7 | Tarn 2 Breeder | Tarn, France | HiSeq |
| Wragg_et_al_2022_MER | SRR15173930 | SAMN20257301 | *A. mellifera* | Unknown | 1 | KF8 | Tarn 2 Breeder | Tarn, France | HiSeq |
| Wragg_et_al_2022_MER | SRR15173929 | SAMN20257302 | *A. mellifera* | Unknown | 1 | KF9 | Tarn 2 Breeder | Tarn, France | HiSeq |
| Wragg_et_al_2022_MER | SRR15173927 | SAMN20257303 | *A. mellifera* | Unknown | 1 | KFA | Tarn 2 Breeder | Tarn, France | HiSeq |
| Wragg_et_al_2022_MER | SRR15173779 | SAMN20257101 | *A. mellifera* | Unknown | 1 | DAN2a | Unknown | Unknown | HiSeq |
| Wragg_et_al_2022_MER | SRR15173778 | SAMN20257102 | *A. mellifera* | Unknown | 1 | DAN2b | Unknown | Unknown | HiSeq |

| | | | | | | | | | |
|---|---|---|---|---|---|---|---|---|---|
| Wragg_et_al_2022_MER | SRR15173702 | SAMN20257171 | *A. mellifera* | Unknown | 1 | ITSAP106 | Unknown | Unknown | HiSeq |
| Wragg_et_al_2022_MER | SRR15173697 | SAMN20257174 | *A. mellifera* | Unknown | 1 | ITSAP13 | Unknown | Unknown | HiSeq |
| Wragg_et_al_2022_MER | SRR15173695 | SAMN20257176 | *A. mellifera* | Unknown | 1 | ITSAP15 | Unknown | Unknown | HiSeq |
| Wragg_et_al_2022_MER | SRR15173692 | SAMN20257179 | *A. mellifera* | Unknown | 1 | ITSAP17 | Unknown | Unknown | HiSeq |
| Wragg_et_al_2022_MER | SRR15173689 | SAMN20257182 | *A. mellifera* | Unknown | 1 | ITSAP19 | Unknown | Unknown | HiSeq |
| Wragg_et_al_2022_MER | SRR15173687 | SAMN20257183 | *A. mellifera* | Unknown | 1 | ITSAP2 | Unknown | Unknown | HiSeq |
| Wragg_et_al_2022_MER | SRR15173674 | SAMN20257195 | *A. mellifera* | Unknown | 1 | ITSAP32 | Unknown | Unknown | HiSeq |
| Wragg_et_al_2022_MER | SRR15173651 | SAMN20257216 | *A. mellifera* | Unknown | 1 | ITSAP6 | Unknown | Unknown | HiSeq |
| Wragg_et_al_2022_MER | SRR15173629 | SAMN20257236 | *A. mellifera* | Unknown | 1 | ITSAP8 | Unknown | Unknown | HiSeq |
| Wragg_et_al_2022_MER | SRR15173528 | SAMN20257377 | *A. mellifera* | Unknown | 1 | TES14 | Unknown | Unknown | HiSeq |
| Wragg_et_al_2022_MER | SRR15173526 | SAMN20257379 | *A. mellifera* | Unknown | 1 | TES22 | Unknown | Unknown | HiSeq |
| Wragg_et_al_2022_MER | SRR15173801 | SAMN20257081 | *A. mellifera* | Unknown | 1 | CIReine-M-Durapi-1 | Vaucluse Breeder | Vaucluse, France | HiSeq |
| Wragg_et_al_2022_MER | SRR15173800 | SAMN20257082 | *A. mellifera* | Unknown | 1 | CIReine-M-Durapi-10 | Vaucluse Breeder | Vaucluse, France | HiSeq |
| Wragg_et_al_2022_MER | SRR15173798 | SAMN20257083 | *A. mellifera* | Unknown | 1 | CIReine-M-Durapi-11 | Vaucluse Breeder | Vaucluse, France | HiSeq |
| Wragg_et_al_2022_MER | SRR15173797 | SAMN20257084 | *A. mellifera* | Unknown | 1 | CIReine-M-Durapi-12 | Vaucluse Breeder | Vaucluse, France | HiSeq |
| Wragg_et_al_2022_MER | SRR15173796 | SAMN20257085 | *A. mellifera* | Unknown | 1 | CIReine-M-Durapi-13 | Vaucluse Breeder | Vaucluse, France | HiSeq |
| Wragg_et_al_2022_MER | SRR15173795 | SAMN20257086 | *A. mellifera* | Unknown | 1 | CIReine-M-Durapi-14 | Vaucluse Breeder | Vaucluse, France | HiSeq |
| Wragg_et_al_2022_MER | SRR15173794 | SAMN20257087 | *A. mellifera* | Unknown | 1 | CIReine-M-Durapi-15 | Vaucluse Breeder | Vaucluse, France | HiSeq |
| Wragg_et_al_2022_MER | SRR15173793 | SAMN20257088 | *A. mellifera* | Unknown | 1 | CIReine-M-Durapi-16 | Vaucluse Breeder | Vaucluse, France | HiSeq |
| Wragg_et_al_2022_MER | SRR15173792 | SAMN20257089 | *A. mellifera* | Unknown | 1 | CIReine-M-Durapi-17 | Vaucluse Breeder | Vaucluse, France | HiSeq |
| Wragg_et_al_2022_MER | SRR15173791 | SAMN20257090 | *A. mellifera* | Unknown | 1 | CIReine-M-Durapi-18 | Vaucluse Breeder | Vaucluse, France | HiSeq |
| Wragg_et_al_2022_MER | SRR15173790 | SAMN20257091 | *A. mellifera* | Unknown | 1 | CIReine-M-Durapi-19 | Vaucluse Breeder | Vaucluse, France | HiSeq |
| Wragg_et_al_2022_MER | SRR15173789 | SAMN20257092 | *A. mellifera* | Unknown | 1 | CIReine-M-Durapi-2 | Vaucluse Breeder | Vaucluse, France | HiSeq |
| Wragg_et_al_2022_MER | SRR15173787 | SAMN20257093 | *A. mellifera* | Unknown | 1 | CIReine-M-Durapi-20 | Vaucluse Breeder | Vaucluse, France | HiSeq |
| Wragg_et_al_2022_MER | SRR15173786 | SAMN20257094 | *A. mellifera* | Unknown | 1 | CIReine-M-Durapi-3 | Vaucluse Breeder | Vaucluse, France | HiSeq |
| Wragg_et_al_2022_MER | SRR15173785 | SAMN20257095 | *A. mellifera* | Unknown | 1 | CIReine-M-Durapi-4 | Vaucluse Breeder | Vaucluse, France | HiSeq |
| Wragg_et_al_2022_MER | SRR15173784 | SAMN20257096 | *A. mellifera* | Unknown | 1 | CIReine-M-Durapi-5 | Vaucluse Breeder | Vaucluse, France | HiSeq |
| Wragg_et_al_2022_MER | SRR15173783 | SAMN20257097 | *A. mellifera* | Unknown | 1 | CIReine-M-Durapi-6 | Vaucluse Breeder | Vaucluse, France | HiSeq |
| Wragg_et_al_2022_MER | SRR15173782 | SAMN20257098 | *A. mellifera* | Unknown | 1 | CIReine-M-Durapi-7 | Vaucluse Breeder | Vaucluse, France | HiSeq |
| Wragg_et_al_2022_MER | SRR15173781 | SAMN20257099 | *A. mellifera* | Unknown | 1 | CIReine-M-Durapi-8 | Vaucluse Breeder | Vaucluse, France | HiSeq |
| Wragg_et_al_2022_MER | SRR15173780 | SAMN20257100 | *A. mellifera* | Unknown | 1 | CIReine-M-Durapi- | Vaucluse Breeder | Vaucluse, France | HiSeq |

| | | | | | | | | | |
|---|---|---|---|---|---|---|---|---|---|
| | | | | | | 9 | | | |
| Cao_et_al_2023_GBE | SRR23343482 | SAMN33097019 | *A. dorsata* | *A. dorsata* | 2 | D10 | XS | Xishuangbanna, Yunnan, China | NovaSeq |
| Cao_et_al_2023_GBE | SRR23343440 | SAMN33097024 | *A. dorsata* | *A. dorsata* | 2 | D12 | PE | Puer, Yunnan, China | NovaSeq |
| Cao_et_al_2023_GBE | SRR23343438 | SAMN33097026 | *A. dorsata* | *A. dorsata* | 2 | D14 | PE | Puer, Yunnan, China | NovaSeq |
| Cao_et_al_2023_GBE | SRR23343441 | SAMN33097023 | *A. dorsata* | *A. dorsata* | 2 | D18 | XS | Xishuangbanna, Yunnan, China | NovaSeq |
| Cao_et_al_2023_GBE | SRR23343474 | SAMN33097044 | *A. dorsata* | *A. dorsata* | 2 | D23 | NN | Nanning, Guangxi, China | NovaSeq |
| Cao_et_al_2023_GBE | SRR23343479 | SAMN33097039 | *A. dorsata* | *A. dorsata* | 2 | D27 | CZ | Chongzuo, Guangxi, China | NovaSeq |
| Cao_et_al_2023_GBE | SRR23343477 | SAMN33097041 | *A. dorsata* | *A. dorsata* | 2 | D29 | CZ | Chongzuo, Guangxi, China | NovaSeq |
| Cao_et_al_2023_GBE | SRR23343475 | SAMN33097043 | *A. dorsata* | *A. dorsata* | 2 | D7 | NN | Nanning, Guangxi, China | NovaSeq |
| Cao_et_al_2023_GBE | SRR23343458 | SAMN33097058 | *A. laboriosa* | *A. laboriosa* | 2 | HD14 | BS | Baoshan, Yunnan, China | NovaSeq |
| Cao_et_al_2023_GBE | SRR23343454 | SAMN33097062 | *A. laboriosa* | *A. laboriosa* | 2 | HD18 | DQ | Diqing, Yunnan | NovaSeq |
| Cao_et_al_2023_GBE | SRR23343453 | SAMN33097063 | *A. laboriosa* | *A. laboriosa* | 2 | HD191 | DQ | Diqing, Yunnan | NovaSeq |
| Cao_et_al_2023_GBE | SRR23343445 | SAMN33097070 | *A. laboriosa* | *A. laboriosa* | 2 | HD22 | RK | Rikaze, Tibet | NovaSeq |
| Cao_et_al_2023_GBE | SRR23343442 | SAMN33097073 | *A. laboriosa* | *A. laboriosa* | 2 | HD25 | RK | Rikaze, Tibet | NovaSeq |
| Cao_et_al_2023_GBE | SRR23343452 | SAMN33097064 | *A. laboriosa* | *A. laboriosa* | 2 | HD26 | LZ | Linzhi, Tibet | NovaSeq |
| Cao_et_al_2023_GBE | SRR23343450 | SAMN33097066 | *A. laboriosa* | *A. laboriosa* | 2 | HD28 | LZ | Linzhi, Tibet | NovaSeq |
| Cao_et_al_2023_GBE | SRR23343462 | SAMN33097055 | *A. laboriosa* | *A. laboriosa* | 2 | HD2 | BS | Baoshan, Yunnan, China | NovaSeq |
| Cao_et_al_2023_GBE | SRR23343473 | SAMN33097045 | *A. laboriosa* | *A. laboriosa* | 2 | HD4 | HH | Honghe, Yunnan, China | NovaSeq |
| Cao_et_al_2023_GBE | SRR23343468 | SAMN33097049 | *A. laboriosa* | *A. laboriosa* | 2 | HD8 | HH | Honghe, Yunnan, China | NovaSeq |
| Chen_et_al_2018_MBE | SRR6301378 | SAMN08038300 | *A. cerana* | *A. cerana* | 2 | AK04 | AK | Ankang, Shanxi | HiSeq |
| Chen_et_al_2018_MBE | SRR6301375 | SAMN08038305 | *A. cerana* | *A. cerana* | 2 | AK09 | AK | Ankang, Shanxi | HiSeq |
| Chen_et_al_2018_MBE | SRR6301346 | SAMN08038371 | *A. cerana* | *A. cerana* | 2 | BM05 | BM | Bomi, Tibet | HiSeq |
| Chen_et_al_2018_MBE | SRR6301345 | SAMN08038374 | *A. cerana* | *A. cerana* | 2 | BM08 | BM | Bomi, Tibet | HiSeq |
| Chen_et_al_2018_MBE | SRR6301311 | SAMN08038389 | *A. cerana* | *A. cerana* | 2 | DQ03 | DQ | Diqing, Yunnan | HiSeq |
| Chen_et_al_2018_MBE | SRR6301306 | SAMN08038392 | *A. cerana* | *A. cerana* | 2 | DQ06 | DQ | Diqing, Yunnan | HiSeq |
| Chen_et_al_2018_MBE | SRR6301297 | SAMN08038318 | *A. cerana* | *A. cerana* | 2 | GY02 | GY | Guyuan, Ningxia | HiSeq |
| Chen_et_al_2018_MBE | SRR6301296 | SAMN08038319 | *A. cerana* | *A. cerana* | 2 | GY03 | GY | Guyuan, Ningxia | HiSeq |
| Chen_et_al_2018_MBE | SRR6301437 | SAMN08038408 | *A. cerana* | *A. cerana* | 2 | HC02 | HC | Hechi, Guangxi | HiSeq |
| Chen_et_al_2018_MBE | SRR6301438 | SAMN08038413 | *A. cerana* | *A. cerana* | 2 | HC07 | HC | Hechi, Guangxi | HiSeq |
| Chen_et_al_2018_MBE | SRR6301400 | SAMN08038431 | *A. cerana* | *A. cerana* | 2 | HK05 | HK | Haikou, Hainan | HiSeq |
| Chen_et_al_2018_MBE | SRR6301395 | SAMN08038436 | *A. cerana* | *A. cerana* | 2 | HK10 | HK | Haikou, Hainan | HiSeq |
| Chen_et_al_2018_MBE | SRR6301320 | SAMN08038348 | *A. cerana* | *A. cerana* | 2 | JZ02 | JZ | Jiuzhai, Sichuan | HiSeq |
| Chen_et_al_2018_MBE | SRR6301314 | SAMN08038354 | *A. cerana* | *A. cerana* | 2 | JZ08 | JZ | Jiuzhai, Sichuan | HiSeq |
| Chen_et_al_2018_MBE | SRR6301294 | SAMN08038339 | *A. cerana* | *A. cerana* | 2 | ME03 | ME | Maerkang, Sichuan | HiSeq |
| Chen_et_al_2018_MBE | SRR6301285 | SAMN08038346 | *A. cerana* | *A. cerana* | 2 | ME10 | ME | Maerkang, Sichuan | HiSeq |
| Chen_et_al_2018_MBE | SRR6301270 | SAMN08038267 | *A. cerana* | *A. cerana* | 2 | MX01 | MX | Minxian, Gansu | HiSeq |
| Chen_et_al_2018_MBE | SRR6301266 | SAMN08038271 | *A. cerana* | *A. cerana* | 2 | MX05 | MX | Minxian, Gansu | HiSeq |
| Chen_et_al_2018_MBE | SRR6301409 | SAMN08038308 | *A. cerana* | *A. cerana* | 2 | MY02 | MY | Mengyin, Shandong | HiSeq |
| Chen_et_al_2018_MBE | SRR6301407 | SAMN08038310 | *A. cerana* | *A. cerana* | 2 | MY04 | MY | Mengyin, Shandong | HiSeq |
| Chen_et_al_2018_MBE | SRR6301329 | SAMN08038378 | *A. cerana* | *A. cerana* | 2 | NY02 | NY | Nayong, Guizhou | HiSeq |
| Chen_et_al_2018_MBE | SRR6301328 | SAMN08038379 | *A. cerana* | *A. cerana* | 2 | NY03 | NY | Nayong, Guizhou | HiSeq |
| Chen_et_al_2018_MBE | SRR6301325 | SAMN08038264 | *A. cerana* | *A. cerana* | 2 | QY08 | QY | Qingyuan, Liaoning | HiSeq |

| | | | | | | | | | |
|---|---|---|---|---|---|---|---|---|---|
| Chen_et_al_2018_MBE | SRR6301364 | SAMN08038265 | *A. cerana* | *A. cerana* | 2 | QY09 | QY | Qingyuan, Liaoning | HiSeq |
| Chen_et_al_2018_MBE | SRR6301355 | SAMN08038332 | *A. cerana* | *A. cerana* | 2 | SJ06 | SJ | Shennongjia, Hubei | HiSeq |
| Chen_et_al_2018_MBE | SRR6301353 | SAMN08038336 | *A. cerana* | *A. cerana* | 2 | SJ10 | SJ | Shennongjia, Hubei | HiSeq |
| Chen_et_al_2018_MBE | SRR6301368 | SAMN08038364 | *A. cerana* | *A. cerana* | 2 | SN08 | SN | Suining, Sichuan | HiSeq |
| Chen_et_al_2018_MBE | SRR6301362 | SAMN08038366 | *A. cerana* | *A. cerana* | 2 | SN10 | SN | Suining, Sichuan | HiSeq |
| Chen_et_al_2018_MBE | SRR6301417 | SAMN08038288 | *A. cerana* | *A. cerana* | 2 | XA02 | XA | Xian, Shanxi | HiSeq |
| Chen_et_al_2018_MBE | SRR6301422 | SAMN08038293 | *A. cerana* | *A. cerana* | 2 | XA07 | XA | Xian, Shanxi | HiSeq |
| Chen_et_al_2018_MBE | SRR6301280 | SAMN08038404 | *A. cerana* | *A. cerana* | 2 | XS08 | XS | Xishuangbanna, Yunnan, China | HiSeq |
| Chen_et_al_2018_MBE | SRR6301283 | SAMN08038405 | *A. cerana* | *A. cerana* | 2 | XS09 | XS | Xishuangbanna, Yunnan, China | HiSeq |
| Chen_et_al_2018_MBE | SRR6301393 | SAMN08038277 | *A. cerana* | *A. cerana* | 2 | YL01 | YL | Yulin, Shanxi | HiSeq |
| Chen_et_al_2018_MBE | SRR6301391 | SAMN08038279 | *A. cerana* | *A. cerana* | 2 | YL03 | YL | Yulin, Shanxi | HiSeq |
| Chen_et_al_2018_MBE | SRR6301426 | SAMN08038422 | *A. cerana* | *A. cerana* | 2 | ZZ06 | ZZ | Zhangzhou, Fujian | HiSeq |
| Chen_et_al_2018_MBE | SRR6301425 | SAMN08038423 | *A. cerana* | *A. cerana* | 2 | ZZ07 | ZZ | Zhangzhou, Fujian | HiSeq |

*Table S2: List of samples excluded as a result of high within-population structure or kinship.*

| Population | Individual | Reason | Method |
|---|---|---|---|
| Buckfast | ITSAP3 | Within-group population structure (Axis1 10.22% of the inertia) | PCA (SNPRelate) |
| Buckfast | ITSAP6B | Within-group population structure (Axis1 10.22% of the inertia) | PCA (SNPRelate) |
| Buckfast | ITSAP7 | Within-group population structure (Axis1 10.22% of the inertia) | PCA (SNPRelate) |
| Buckfast | ITSAP10 | Within-group population structure (Axis1 10.22% of the inertia) | PCA (SNPRelate) |
| Buckfast | ITSAP71 | Within-group population structure (Axis1 10.22% of the inertia) | PCA (SNPRelate) |
| Buckfast | CIReine-M-Durapi-16 | Within-group population structure (Axis1 10.22% of the inertia) | PCA (SNPRelate) |
| Caucasia | BR36 | Within-group population structure (Axis1: 11.22% of the inertia) | PCA (SNPRelate) |
| Caucasia | CIReine-M-Durapi-18 | Within-group population structure (Axis2: 10.68% of the inertia) | PCA (SNPRelate) |
| Ligustica | ITA8A | Within-group population structure (Axis1: 8.83% of the inertia) | PCA (SNPRelate) |
| Ligustica | ITA15A | Within-group population structure (Axis 2: 6.76% of the inertia) | PCA (SNPRelate) |
| Mellifera | POR11 | Within-group population structure (Axis1: 8.52% of the inertia) | PCA (SNPRelate) |
| Mellifera | POR13 | Within-group population structure (Axis2: 7.86% of the inertia) | PCA (SNPRelate) |
| Buckfast | DAN2b | Involved in one close-related pair and has lower call-rate (99.7%) than mate | IBD (hmmIBD) |
| Buckfast | ITSAP78 | Involved in one close-related pair and has lower call-rate (99.9%) than mate | IBD (hmmIBD) |
| Buckfast | ITSAP101 | Involved in one close-related pair and has lower call-rate (99.9%) than mate | IBD (hmmIBD) |
| Carnica (Germany) | BER6 | Involved in one close-related pair and has lower call-rate (99.0%) than mate | IBD (hmmIBD) |
| Carnica | SLO1 | Involved in 2 close-related pairs | IBD (hmmIBD) |
| Carnica | SLO19 | Involved in 2 close-related pairs | IBD (hmmIBD) |
| Carnica | ITSAP16 | Involved in one close-related pair and has lower call-rate (99.9%) than mate | IBD (hmmIBD) |
| Carnica | POL23 | Involved in one close-related pair and has lower call-rate (99.7%) than mate | IBD (hmmIBD) |
| Carnica | SLO10 | Involved in one close-related pair and has lower call-rate (99.7%) than mate | IBD (hmmIBD) |
| Carnica | SLO15 | Involved in one close-related pair and has lower call-rate (99.5%) than mate | IBD (hmmIBD) |
| Carnica | SLO7 | Involved in one close-related pair and has lower call-rate (99.3%) than mate | IBD (hmmIBD) |
| Carnica | SLO2 | Involved in one close-related pair and has lower call-rate (99.5%) than mate | IBD (hmmIBD) |
| Colonsay | UK5A | Involved in one close-related pair and has lower call-rate (99.6%) than mate | IBD (hmmIBD) |
| Colonsay | UK4A | Involved in one close-related pair and has lower call-rate (99.5%) than mate | IBD (hmmIBD) |
| Corsica | AOC4 | Involved in one close-related pair and has lower call-rate (81.9%) than mate | IBD (hmmIBD) |
| Mellifera | POR7 | Involved in one close-related pair and has lower call-rate (99.9%) than mate | IBD (hmmIBD) |
| Mellifera | SOL14 | Involved in one close-related pair and has lower call-rate (99.9%) than mate | IBD (hmmIBD) |
| Mellifera | POR12 | Involved in one close-related pair and has lower call-rate (99.8%) than mate | IBD (hmmIBD) |
| Ouessant | OUE15 | Involved in one close-related pair and has lower call-rate (98.6%) than mate | IBD (hmmIBD) |
| Ouessant | OUE1 | Involved in one close-related pair and has lower call-rate (95.2%) than mate | IBD (hmmIBD) |
| Ouessant | OUE18 | Involved in one close-related pair and has lower call-rate (96.0%) than mate | IBD (hmmIBD) |
| Ouessant | OUE34 | Involved in one close-related pair and has lower call-rate (99.9%) than mate | IBD (hmmIBD) |
| Ouessant | OUE29 | Involved in one close-related pair and has lower call-rate (99.8%) than mate | IBD (hmmIBD) |
| RoyalJelly | YC6 | Involved in 4 close-related pairs | IBD (hmmIBD) |
| RoyalJelly | ITSAP11 | Involved in 3 close-related pairs | IBD (hmmIBD) |
| RoyalJelly | ITSAP31 | Involved in 2 close-related pairs | IBD (hmmIBD) |
| RoyalJelly | ITSAP65 | Involved in 2 close-related pairs | IBD (hmmIBD) |
| RoyalJelly | YC1 | Involved in 2 close-related pairs | IBD (hmmIBD) |
| RoyalJelly | ITSAP81 | Involved in one close-related pair and has lower call-rate (99.9%) than mate | IBD (hmmIBD) |
| RoyalJelly | ITSAP24 | Involved in one close-related pair and has lower call-rate (99.8%) than mate | IBD (hmmIBD) |
| RoyalJelly | FL3 | Involved in one close-related pair and has lower call-rate (91.6%) than mate | IBD (hmmIBD) |
| RoyalJelly | TES22 | Involved in one close-related pair and has lower call-rate (99.0%) than mate | IBD (hmmIBD) |
| RoyalJelly | YC3 | Involved in one close-related pair and has lower call-rate (99.9%) than mate | IBD (hmmIBD) |
| RoyalJelly | YC2 | Involved in one close-related pair and has lower call-rate (99.9%) than mate | IBD (hmmIBD) |

*Table S3: List of the 267 A. mellifera samples included in the study*

| Population | Individual |
|---|---|
| Buckfast | CIReine-M-Durapi-12 |
| Buckfast | ITSAP30 |
| Buckfast | ITSAP82 |
| Buckfast | ITSAP88 |
| Buckfast | JFM19 |
| Buckfast | JFM2 |
| Buckfast | JFM25 |
| Buckfast | JFM29 |
| Buckfast | JFM31 |
| Buckfast | JFM33 |
| Buckfast | JFM6 |
| Buckfast | JFM7 |
| Buckfast | JFM9 |
| Buckfast | DAN2a |
| Buckfast | ITSAP12 |
| Buckfast | ITSAP79 |
| Carnica (German subgroup) | BER10 |
| Carnica (German subgroup) | BER11 |
| Carnica (German subgroup) | BER12 |
| Carnica (German subgroup) | BER13 |
| Carnica (German subgroup) | BER14 |
| Carnica (German subgroup) | BER15 |
| Carnica (German subgroup) | BER18 |
| Carnica (German subgroup) | BER2 |
| Carnica (German subgroup) | BER4 |
| Carnica (German subgroup) | BER7 |
| Carnica (German subgroup) | BER8 |
| Carnica (German subgroup) | BER9 |
| Carnica (German subgroup) | BER19 |
| Carnica (non-German subgroup) | BER16 |
| Carnica (non-German subgroup) | BER5 |
| Carnica (non-German subgroup) | CAR10 |
| Carnica (non-German subgroup) | CAR11 |
| Carnica (non-German subgroup) | CAR12 |
| Carnica (non-German subgroup) | CAR13 |
| Carnica (non-German subgroup) | CAR14 |
| Carnica (non-German subgroup) | CAR15 |
| Carnica (non-German subgroup) | CAR17 |
| Carnica (non-German subgroup) | CAR18 |
| Carnica (non-German subgroup) | CAR19 |
| Carnica (non-German subgroup) | CAR2 |
| Carnica (non-German subgroup) | CAR20 |
| Carnica (non-German subgroup) | CAR21 |
| Carnica (non-German subgroup) | CAR22 |
| Carnica (non-German subgroup) | CAR23 |
| Carnica (non-German subgroup) | CAR25 |
| Carnica (non-German subgroup) | CAR26 |
| Carnica (non-German subgroup) | CAR28 |
| Carnica (non-German subgroup) | CAR29 |
| Carnica (non-German subgroup) | CAR3 |
| Carnica (non-German subgroup) | CAR30 |
| Carnica (non-German subgroup) | CAR31 |

| | |
|---|---|
| Carnica (non-German subgroup) | CAR34 |
| Carnica (non-German subgroup) | CAR4 |
| Carnica (non-German subgroup) | CAR5 |
| Carnica (non-German subgroup) | CAR6 |
| Carnica (non-German subgroup) | CAR7 |
| Carnica (non-German subgroup) | ITSAP106 |
| Carnica (non-German subgroup) | ITSAP17B |
| Carnica (non-German subgroup) | ITSAP23 |
| Carnica (non-German subgroup) | ITSAP38 |
| Carnica (non-German subgroup) | ITSAP42 |
| Carnica (non-German subgroup) | ITSAP48 |
| Carnica (non-German subgroup) | ITSAP55 |
| Carnica (non-German subgroup) | ITSAP6 |
| Carnica (non-German subgroup) | ITSAP60 |
| Carnica (non-German subgroup) | ITSAP61 |
| Carnica (non-German subgroup) | ITSAP63 |
| Carnica (non-German subgroup) | ITSAP64 |
| Carnica (non-German subgroup) | ITSAP9 |
| Carnica (non-German subgroup) | JFM11 |
| Carnica (non-German subgroup) | JFM13 |
| Carnica (non-German subgroup) | JFM26 |
| Carnica (non-German subgroup) | POL15 |
| Carnica (non-German subgroup) | POL2 |
| Carnica (non-German subgroup) | POL5 |
| Carnica (non-German subgroup) | POL6 |
| Carnica (non-German subgroup) | POL7 |
| Carnica (non-German subgroup) | POL9 |
| Carnica (non-German subgroup) | SLO12 |
| Carnica (non-German subgroup) | SLO13 |
| Carnica (non-German subgroup) | SLO14 |
| Carnica (non-German subgroup) | SLO16 |
| Carnica (non-German subgroup) | SLO17 |
| Carnica (non-German subgroup) | SLO18 |
| Carnica (non-German subgroup) | SLO20 |
| Carnica (non-German subgroup) | SLO3 |
| Carnica (non-German subgroup) | SLO5 |
| Carnica (non-German subgroup) | SLO8 |
| Carnica (non-German subgroup) | ITSAP49 |
| Carnica (non-German subgroup) | POL28 |
| Carnica (non-German subgroup) | SLO11 |
| Carnica (non-German subgroup) | SLO4 |
| Carnica (non-German subgroup) | SLO6 |
| Carnica (non-German subgroup) | SLO9 |
| Caucasia | CAU10 |
| Caucasia | CAU11 |
| Caucasia | CAU12 |
| Caucasia | CAU14 |
| Caucasia | CAU15A |
| Caucasia | CAU16A |
| Caucasia | CAU17 |
| Caucasia | CAU17A |
| Caucasia | CAU18A |
| Caucasia | CAU19A |
| Caucasia | CAU20 |

| | |
|---|---|
| Caucasia | CAU21 |
| Caucasia | CAU6 |
| Caucasia | CAU7 |
| Caucasia | KF43 |
| Colonsay | UK10A |
| Colonsay | UK11A |
| Colonsay | UK12A |
| Colonsay | UK13A |
| Colonsay | UK14A |
| Colonsay | UK15A |
| Colonsay | UK16A |
| Colonsay | UK17A |
| Colonsay | UK18A |
| Colonsay | UK19A |
| Colonsay | UK1A |
| Colonsay | UK20A |
| Colonsay | UK21A |
| Colonsay | UK22A |
| Colonsay | UK23A |
| Colonsay | UK25A |
| Colonsay | UK27A |
| Colonsay | UK28A |
| Colonsay | UK29A |
| Colonsay | UK2A |
| Colonsay | UK3A |
| Colonsay | UK7A |
| Colonsay | UK8A |
| Colonsay | UK9A |
| Colonsay | UK26A |
| Colonsay | UK6A |
| Iberiensis | BR15 |
| Iberiensis | BR5 |
| Iberiensis | ESP11 |
| Iberiensis | ESP14 |
| Iberiensis | ESP16 |
| Iberiensis | ESP18 |
| Iberiensis | ESP21 |
| Iberiensis | ESP22 |
| Iberiensis | ESP23 |
| Iberiensis | ESP24 |
| Iberiensis | ESP25 |
| Iberiensis | ESP26 |
| Iberiensis | ESP27 |
| Iberiensis | ESP28 |
| Iberiensis | ESP29 |
| Iberiensis | ESP30 |
| Ligustica | ITA10A |
| Ligustica | ITA11A |
| Ligustica | ITA12A |
| Ligustica | ITA13A |
| Ligustica | ITA14A |
| Ligustica | ITA16A |
| Ligustica | ITA17A |
| Ligustica | ITA18A |

| | |
|---|---|
| Ligustica | ITA19A |
| Ligustica | ITA1A |
| Ligustica | ITA20A |
| Ligustica | ITA22A |
| Ligustica | ITA24A |
| Ligustica | ITA25A |
| Ligustica | ITA26A |
| Ligustica | ITA27A |
| Ligustica | ITA29A |
| Ligustica | ITA2A |
| Ligustica | ITA30A |
| Ligustica | ITA3A |
| Ligustica | ITA4A |
| Ligustica | ITA5A |
| Ligustica | ITA6A |
| Ligustica | ITA7A |
| Ligustica | ITA9A |
| Mellifera | POR1 |
| Mellifera | POR10 |
| Mellifera | POR14 |
| Mellifera | POR15 |
| Mellifera | POR2 |
| Mellifera | POR3 |
| Mellifera | POR4 |
| Mellifera | POR5 |
| Mellifera | POR8 |
| Mellifera | POR9 |
| Mellifera | SOL1 |
| Mellifera | SOL2 |
| Mellifera | SOL6 |
| Mellifera | SOL7 |
| Mellifera | POR6 |
| Mellifera | SOL13 |
| Mellifera | SOL5 |
| Ouessant | Ab-PacBio |
| Ouessant | OUE10 |
| Ouessant | OUE11 |
| Ouessant | OUE12 |
| Ouessant | OUE13 |
| Ouessant | OUE14 |
| Ouessant | OUE16 |
| Ouessant | OUE17 |
| Ouessant | OUE19 |
| Ouessant | OUE2 |
| Ouessant | OUE21 |
| Ouessant | OUE22 |
| Ouessant | OUE23 |
| Ouessant | OUE24 |
| Ouessant | OUE25 |
| Ouessant | OUE26 |
| Ouessant | OUE27 |
| Ouessant | OUE28 |
| Ouessant | OUE30 |
| Ouessant | OUE31 |

| | |
|---|---|
| Ouessant | OUE32 |
| Ouessant | OUE33 |
| Ouessant | OUE36 |
| Ouessant | OUE38 |
| Ouessant | OUE39 |
| Ouessant | OUE4 |
| Ouessant | OUE40 |
| Ouessant | OUE5 |
| Ouessant | OUE6 |
| Ouessant | OUE7 |
| Ouessant | OUE20 |
| Ouessant | OUE3 |
| Ouessant | OUE35 |
| Ouessant | OUE37 |
| Ouessant | OUE9 |
| RoyalJelly | FL1 |
| RoyalJelly | FL106 |
| RoyalJelly | FL11 |
| RoyalJelly | FL12 |
| RoyalJelly | FL13 |
| RoyalJelly | FL15 |
| RoyalJelly | FL17 |
| RoyalJelly | FL1bis |
| RoyalJelly | FL22 |
| RoyalJelly | FL61 |
| RoyalJelly | FL7 |
| RoyalJelly | FL87 |
| RoyalJelly | FL9 |
| RoyalJelly | FL9bis |
| RoyalJelly | ITSAP102 |
| RoyalJelly | ITSAP20 |
| RoyalJelly | ITSAP2B |
| RoyalJelly | ITSAP62 |
| RoyalJelly | ITSAP66 |
| RoyalJelly | NM1 |
| RoyalJelly | NM2 |
| RoyalJelly | NM21 |
| RoyalJelly | NM26 |
| RoyalJelly | NM28 |
| RoyalJelly | NM5 |
| RoyalJelly | NM7 |
| RoyalJelly | NM9 |
| RoyalJelly | XC5 |
| RoyalJelly | YC10 |
| RoyalJelly | YC4 |
| RoyalJelly | YC5 |
| RoyalJelly | YC9 |
| RoyalJelly | ITSAP50 |
| RoyalJelly | ITSAP90 |
| RoyalJelly | NM3 |
| RoyalJelly | ITSAP54 |
| RoyalJelly | YC7 |
| RoyalJelly | YC8 |
